# Resistance to Pharmacological Ascorbate, Thioredoxin Reductase Inhibition, and Radiation in KEAP1 Mutant Lung Cancer Cells is Mediated by NRF2

**DOI:** 10.64898/2026.11.17.767416

**Authors:** S Sho, AM Tomanek-Chalkley, BA Wagner, DL Marren, SA Wasimi, AN Lengel, MQM Gentles, TJ Nels, CF Pulliam, MA Fath, TIA Evans, M Furqan, BG Allen, EB Taylor, AJ Dupuy, DR Spitz

## Abstract

Kelch-like ECH-associated protein 1 (KEAP1) mutations occur in 11-27% of non-small cell lung cancers (NSCLCs), leading to increased activity of nuclear factor erythroid 2 (NRF2), upregulating thiol and peroxide metabolism and promoting cancer therapy resistance. Inhibiting NRF2 target gene, thioredoxin reductase, with FDA approved Auranofin (AUR) inhibits thioredoxin-mediated H_2_O_2_ metabolism and pharmacological ascorbate (P-AscH^-^) selectively increases H_2_O_2_ in NSCLCs.

The current study shows KEAP1 mutant NSCLCs (A549, H460, H2030) were resistant to P-AscH^-^ or AUR, relative to KEAP1 WT (H1299T, Calu6, H292). AUR pretreatment synergistically enhanced P-AscH⁻ cytotoxicity in all NSCLCs but not in primary human lung epithelial cells (HBEpCs). N-Acetyl-L-Cysteine (NAC) or catalase overexpression protected NSCLCs from toxicity induced by AUR + P-AscH⁻ and NAC inhibited glutathione disulfide (GSSG) accumulation demonstrating the involvement of thiol oxidation and H_2_O_2_ in the mechanism.

Knockout (KO) of KEAP1 in H1299T WT increased nuclear NRF2, NRF2 target genes, and increased resistance to radiation or P-AscH^-^. P-AscH^-^ combined with radiation significantly sensitized KEAP1 KO H1299Ts. KO of NRF2 in KEAP1 mutant A549 sensitized them to P-AscH^-^, AUR, and radiation. Significant changes in levels of GSH, GSSG, GSH/thioredoxin metabolic enzymes, and NRF2 target genes occurred in both KEAP1 and NRF2 KO cells. NRF2 KOs also demonstrated significant decreases in NADPH, NADP^+^, mitochondrial function, and Coenzyme A. These findings show the KEAP1–NRF2 pathway is a key mediator of redox-dependent metabolism and therapeutic resistance in NSCLC, supporting targeting thiol/peroxide metabolism using Aur + P-AscH^-^ to enhance treatment efficacy.

**Graphical Abstract:** 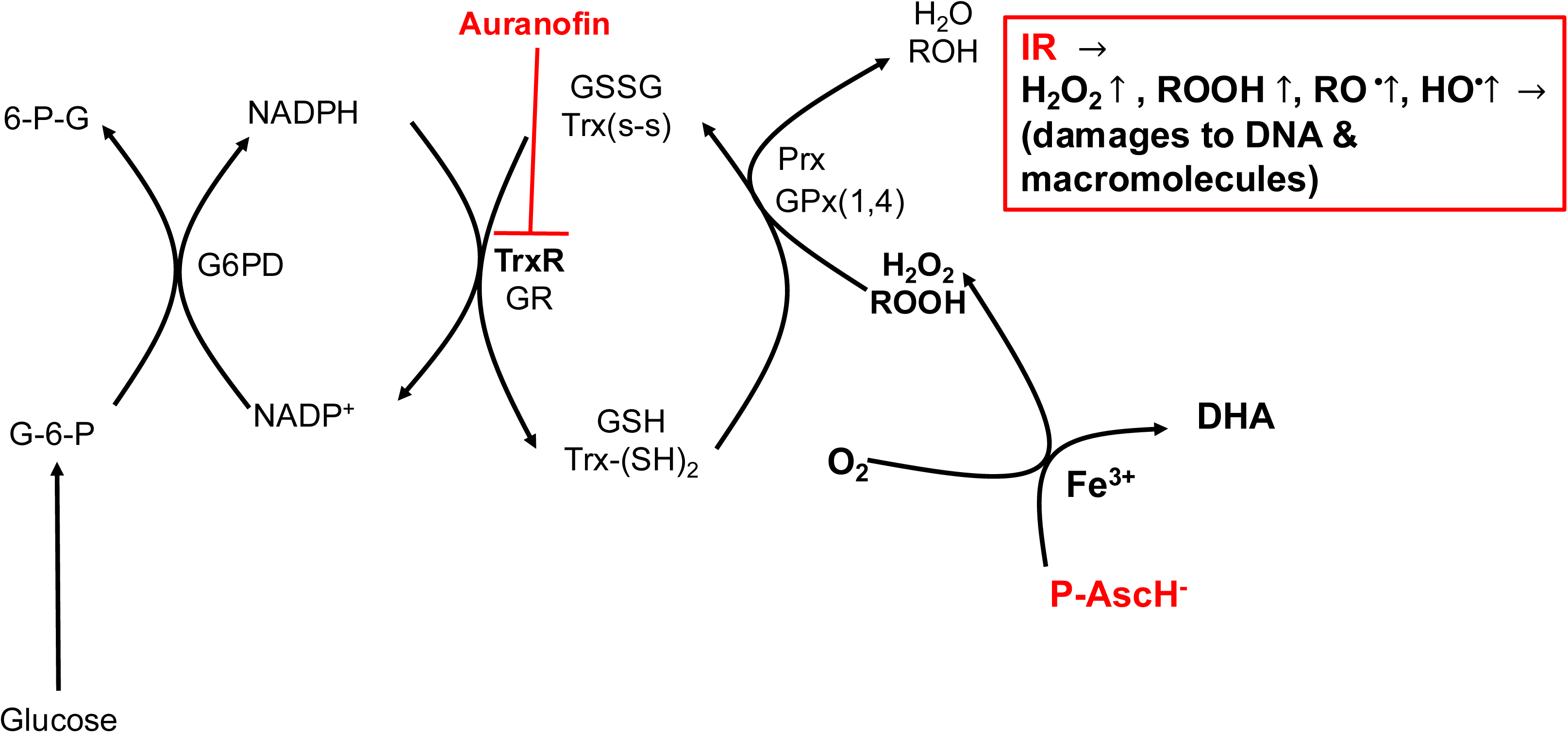

## Introduction

Lung cancer is one of the most common and deadliest cancers in the United States with ∼125,000 estimated deaths each year, accounting for ∼20 % of all cancer deaths with 80-85% of these cases being non-small cell lung cancer (NSCLC) [1]. Despite the significant improvement in survival with the introduction of immune check point inhibitors, only 20-30 % of NSCLC patients respond to this therapy [2, 3]. Furthermore, a large study of NSCLC identified mutations in tumor suppressor KEAP1 (Kelch-like ECH-associated protein 1) in ∼11% of patients [4] and these mutants were refractive to all therapy interventions.

Mechanistically, KEAP1 homodimerizes and interacts with Cullin 3 at its N-terminal BTB domain, and acts as substrate recognition subunit of the functional Cullin 3-RING E3 ubiquitin ligase complex, which targets nuclear factor-erythroid factor 2 (NRF2) transcription factor for degradation under un stressed conditions [4]. KEAP1 mutations result in constitutively active NRF2, upregulation of thioredoxin/glutathione metabolism, alterations in iron metabolism, production of NADPH, and increased resistance to radio-chemotherapy, immunotherapy, and tyrosine kinase inhibitors [5–10].

Pharmacological ascorbate (P-AscH^-^, given intravenously, plasma concentration of ≥ 20 mM) is known to specifically cause cytotoxicity in cancer cells [11–13]. This P-AscH^-^ induced cancer cell specific toxicity is accompanied by sustained increases in steady-state levels of reactive oxygen species (ROS) and labile iron levels due to changes in mitochondrial oxidative metabolism [11, 13]. Furthermore, in normal cells, P-AscH^-^ acts as a donor antioxidant [12, 14].

The prooxidant mechanism of P-AscH^-^ involves the reduction of ferric iron (Fe^3+^) to labile ferrous iron (Fe^2+^) causing release from ferritin (and other Fe containing proteins) increasing labile iron pools in cancer cells enhancing treatment efficacy when combined with radio-chemotherapy [15–18]. Phase I/II clinical trials have been completed in glioblastoma, pancreatic cancer, sarcoma, and stage IV NSCLC (NCT01752491; NCT02344355; NCT02420314; NCT02905578; NCT02905591; NCT03146962; NCT03541486; NCT04033107). The published phase II trial in stage IV NSCLC demonstrated that 34.2 % of the patients showed partial response to 4 cycles of P-AscH^-^ IV administration as an adjuvant (75 g twice/week) of the platinum doublet chemotherapy (Carboplatin + Paclitaxel every three weeks), and improvement in OS, PFS, and the disease control rate (stable disease and partial response) was also observed [17]. Moreover, the randomized phase II trial in stage IV metastatic pancreatic ductal adenocarcinoma (PDAC) in which 75 g P-AscH^-^ was given three times weekly as an adjuvant with chemotherapy (gemcitabine and paclitaxel) also showed significant increases in OS and PFS, and no increase in treatment-related toxicity [16].

Auranofin, the gold phosphine compound that was originally FDA-approved for rheumatoid arthritis in 1985, is also suggested to cause cancer specific toxicity by inhibiting thioredoxin reductase (TrxR) and glutathione peroxidase (GPx), altering thiol and H_2_O_2_ metabolism [19–21]. Pilot feasibility studies and phase I/II clinical trials have been pursued with auranofin as a monotherapy or in combination with sirolimus in NSCLC, ovarian cancer, peritoneal cancer, and small cell lung cancer, but limited tumor responses were observed (NCT01747798; NCT02126527).

The current study demonstrates that KEAP1 mutant NSCLCs (A549, H460, H2030) were resistant to P-AscH^-^ or AUR, relative to KEAP1 WT (H1299T, Calu-6, H292). AUR pretreatment synergistically enhanced P-AscH⁻ cytotoxicity in all NSCLCs but not in primary human lung epithelial cells (HBEpCs). The thiol antioxidant N-Acetyl-L-Cysteine (NAC) [22] or the overexpression of catalase which scavenges H_2_O_2_ inhibited the toxicity induced by AUR + P-AscH⁻ in NSCLCs and NAC also inhibited glutathione disulfide (GSSG) accumulation demonstrating the involvement of thiol oxidation and H_2_O_2_ in the mechanism causing cancer cell specific toxicity.

CRISPR knockout (KO) of KEAP1 in H1299T WT increased nuclear NRF2, NRF2 target genes, and increased resistance to radiation or P-AscH^-^. P-AscH^-^ combined with radiation *in vitro* significantly sensitized KEAP1 KO H1299Ts. CRISPR KO of NRF2 in KEAP1 mutant A549 sensitized them to P-AscH^-^, AUR, and radiation. Biochemical and transcriptomic analysis demonstrated significant changes in levels of GSH, GSSG, GSH/thioredoxin metabolic enzymes, and many other antioxidant NRF2 target genes occurred in both KEAP1 and NRF2 KO cells. Furthermore, metabolomic and Seahorse-XF analysis of NRF2 KOs also demonstrated significant decreases in NADPH, NADP+, mitochondrial function, and Coenzyme A. These findings show the KEAP1–NRF2 pathway is a key mediator of redox-dependent oxidative metabolism and therapeutic resistance in NSCLC, supporting the hypothesis that targeting thiol/peroxide metabolism using Aur + P-AscH^-^ could be an effective adjuvant for enhancing cancer cells specific therapeutic efficacy.

## Materials and Methods

### Cell Lines and Reagents

A549, Calu-6, H1299, H2030, H292, H460 cells were purchased from ATCC (CCL-185, HTB-56, CRL-5803, CRL-5914, CRL-1848, HTB-177). H1299T cells were derived from H1299 as described previously [23]. All NSCLC cell lines were maintained in RPMI 1640 media (#11875093; Life Technologies) supplemented with 10% Fetal Bovine Serum (FBS) (FBS; Atlanta Biologicals, Flowery Brach, GA, USA) in a humidified 37°C incubator with 5% CO_2_, 4 (physiological) or 21% (normoxic) O_2_ [22, 24]. A549CAT2 and H1299TCAT15 doxy-inducible catalase overexpressing cells were generated by performing creation of lentivirus utilizing HEK293/HEK293FT cells, followed by viral transduction and cloning [25, 26]. H1299T KEAP1 Knockout (KO) cells and A549 NRF2 KO cells were generated from H1299T cells and A549 cells respectively, as described below. Normal lung epithelial cells (HBEpC) were purchased from ATCC (PCS-300-010) and were maintained in Bronchial/Tracheal Epithelial Cell Growth Medium (# 511-500; Cell Applications, Inc., San Diego, CA, USA) in a humidified 37°C incubator, 5% CO_2_, room air, and 4% O_2_.

### Drug Delivery

Auranofin (# BML-EI206-0100; Enzo Life Sciences Inc., Farmingdale, NY, USA) was dissolved in 200 proof ethanol (# 2716; Decon Labs, Inc., King of Prussia, PA, USA) to make 1 or 10 mM solution. L-Ascorbic Acid solution (approximately 1 M) was prepared in Nanopure Type 1 water, and the pH was adjusted to 7.0 with NaOH. Precise concentration was determined spectrophotometrically as previously described (ε265 = 14,500 M^−1^ cm^−1^) [27–30]. 1M N-Acetyl-L-Cysteine (NAC) solution was prepared fresh each use by combining 0.4896 g with 0.2840 g NaHCO_3_ in water and sterilizing via filtration. For combination treatments, cells were exposed to auranofin (or vehicle control) for 3 or 24 hours followed by P-AscH^-^ for 1 hour (**exposure schemes shown in Fig. S2C, S3F, and S11C**). In the case of doxy-inducible catalase, cells were exposed to 1 µg/mL doxy for 24 hours prior to exposure to P-AscH^-^ or auranofin (**exposure scheme shown in Figure S5G).** In the case of testing the thiol antioxidant, cells were treated with NAC (15 mM) for 3 hours prior to auranofin for 21 hours followed by P-AscH^-^ for 1 hour (**exposure scheme shown in Figure S5G**).

For the NSCLCs that were maintained in 21% O_2_, P-AscH^-^ was delivered from 1 M stock in FBS-free RPMI 1640 media pre-equilibrated to 21% O_2_ for 1 hour. Since HBEpC cells were maintained in 4% O_2_ + 5% CO_2_ to increase their growth and clonogenic survival, for comparison of HBEpCs to NSCLCs, all cultures were maintained and cloned in 4% O_2_ as well as P-AscH^-^ or auranofin were delivered to all cell lines in Bronchial/Tracheal Epithelial Cell Growth Medium pre-equilibrated to 4% O_2_ for 1 hour.

### Ionizing Radiation

Ionizing radiation (IR) was delivered at the Radiation and Free Radical Research core facility at University of Iowa, with an Xstrahl CIX3 Cabinet irradiator, which operates with maximum voltage of 320 kV, maximum field size of 51.3 cm diameter at 70 cm FSD (Focus-to-Skin Distance), and dose rate of 71.4 cGy per minute. Cells were irradiated in 60 mm cell culture dishes.

### Clonogenic Survival Assay

Approximately 40,000-150,000 were plated in 60 mm dishes and were allowed to grow for 24-72 hours. For dose response experiments, 14 NSCLC cells (A549, Calu-6, H1299, H1299T, H2030, H292, H460, H1299T single guide RNA control (sg CTRL), H1299T KEAP1 KO1, KEAP1 KO2, A549 sgCTRL1, A549 sgCTRL2, A549 NRF2 Complete KO, and A549 NRF2 Isoform Specific KO) were maintained in 21% O_2_ and treated with 0 - 40 pmol/cell (0 - 12.5 mM) P-AscH^-^ for 1 hour, auranofin (0 - 8 µM) for 24 hours, or irradiated 0 - 14 Gy. HBEpC cells and 2 NSCLC cells (A549, H1299T) maintained in 4% O_2_ were treated with 1 – 5 µM auranofin for 3 hours and followed by 0 -160 pmol/cell (0 - 40 mM) P-AscH^-^ for 1 hour. In some experiments, 10 NSCLC cells (A549, Calu-6, H1299, H1299T, H2030, H292, H460, H1299T sgCTRL, H1299T KEAP1 KO1, H1299T KEAP1 KO2) were treated with auranofin (0.5 or 1 µM) for 24 hours followed by P-AscH^-^ for 1 hour (5 - 40 pmol/cell [2.5-12.5 mM]), and A549, A549CAT2 and H1299TCAT15 cells were treated with 15 mM NAC for 3 hours prior to the 21-hour auranofin treatment (0.5 or 1 µM), followed by 1-hour P-AscH^-^ treatment (5 or 40 pmol/cell [2.5 or 12.5 mM]). The doses of auranofin and P-AscH^-^ were chosen to yield 60 – 80% survival and 40 – 60% survival, respectively, based on the dose response curves. For Catalase induction in A549CAT2 and H1299TCAT15 cells, 1 µg/mL doxy was added 24 hours prior to the treatments with auranofin and P-AscH^-^. H1299T sgCTRL, H1299T KEAP1 KO1, and H1299T KEAP1 KO2 were also treated with P-AscH^-^ for 1 hour and were irradiated 2 or 4 Gy (Timeline is shown in **Fig. S9**). The doses of radiation were chosen to yield 60 – 80% survival based on the dose response curves. After treatments, both floating and attached cells were collected, and total number of cells were counted from each 60 mm cell culture dishes using a Beckman Coulter Counter (Beckman Coulter Inc., Hialeah, FL, USA). 150-200,000 cells from each cell culture dish were seeded into 6-well cell culture plates in triplicates for quantification. After 7 - 14 days of incubation at 37°C in complete media with 0.1% gentamycin or 50-100 units/mL penicillin & 50-100 µg/mL streptomycin, cells were fixed with 70% ethanol and were stained with Coomassie Blue. Colonies of more than 50 cells were counted and plotted to calculate plating efficiency, which is defined as:

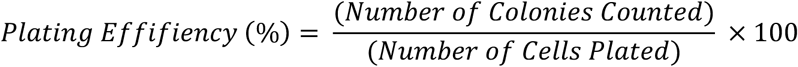

The plating efficiency of experimental groups were normalized to that of control group to calculate the normalized survival fractions. In the experiments where 1 µg/mL doxy was added, plating efficiencies of experimental groups were normalized to the control group that was treated with 1 µg/mL doxy. The doubling time of the cells were calculated based on the number of cells plated and total cell counts from control groups after they were grown in culture for 24-72 hours using the formula:

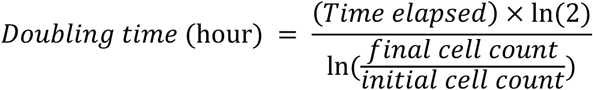

In the experiments with HBEpC cells, 75,000 irradiated (25 Gy) Chinese hamster ovary (CHO) B1 feeder cells were plated into 6-well cell culture plates [31, 32].

**Catalase Activity Assay:**

Cell pellets were homogenized in potassium phosphate buffer (50 mM, pH 7.0), sonicated in water bath sonicator (Sonics Vibra-Cell (Sonics & Materials, Inc., Newtown, CT, USA) at low power three times, and protein concentration was determined using Pierce^TM^ BCA Protein Assay Kit (#23225; Thermo Fisher Scientific, Waltham, MA, USA). Catalase activity was determined at 25 °C as previously described [33, 34]. Briefly, 75-100 µL of samples were added into 4 mL K_2_HPO_4_ buffer and were separated into two quartz cuvettes and were defined as blank and sample groups. 1 mL K_2_HPO_4_ buffer was mixed as the blank. 1 mL K_2_HPO_4_ buffer + 10 mM H_2_O_2_ was added into sample group, and the rate of disappearance of absorption at 240 nm was measured. All activity assays were normalized per mg protein in cell samples.

### Total Glutathione and Glutathione Disulfide Assay

A549 sgCTRL1, A549 sgCTRL2, A549 NRF2 Complete KO, A549 NRF2 Isoform Specific KO, H1299T sgCTRL, H1299T KEAP1 KO1, and H1299T KEAP1 KO2 were plated in 60 mm dishes, allowed to grow for 48 hours. A549 and H1299TCAT15 were plated in 60 mm dishes, allowed to grow for 24 hours, and were either treated with 1 µg/mL doxy for 24 hours or 15 mM NAC for 3 hours, followed by 21-hour auranofin treatment and 1-hour P-AscH^-^ treatment. Then the cultures were washed with DPBS (#14190144; Life Technologies, Carlsbad, CA), and the cells were scraped into 5% 5-sulfosalicylic acid (#S-7408; Sigma-Aldrich, St. Louis, MO) solution and stored in -80 °C for maximum of three weeks. The samples were thawed and centrifuged, and the supernatant was used for total (GSH + GSSG) and oxidized (GSSG) glutathione assay. Protein pellets were dissolved in 0.1N NaOH and 1% SDS in ddH_2_O at 37 °C for overnight before determining the protein concentration using Pierce^TM^ BCA Protein Assay Kit. Total glutathione and glutathione disulfide were determined as previously described [35, 36].

### Thioredoxin Reductase Assay

TrxR activity was determined spectrophotometrically as previously described [37]. Briefly, cell pellets were homogenized in Diethylenetriaminepentaacetic acid (DETAPAC, #D651810; Sigma-Aldrich), sonicated in water bath sonicator at low power three times, and protein concentration was determined using Pierce^TM^ BCA Protein Assay Kit. The formation of TNB in the presence of NADPH in samples were measured with or without inhibitor (#ab83463; Abcam, Waltham, MA, USA). The increase of absorbance at 412 nm/minute from the sample + inhibitor group was subtracted from that of the sample group and was divided by the protein concentration to calculate the amount of TrxR activity in each sample.

### Glutathione Peroxidase 1 Activity Assay

Cell pellets were homogenized in DETAPAC, sonicated in water bath sonicator at low power three times, and protein concentration was determined using Pierce^TM^ BCA Protein Assay Kit. GPx1 activity was determined at 25 °C as previously described [38–40] using H_2_O_2_ as substrate.

### Generation of CRISPR Knockout Cell Lines

sgRNA oligos targeting KEAP1 and NRF2 (sequences listed below) and Alt-R^TM^ CRISPR-Cas9 Control Kit (Human) (#1072554) were purchased from Integrated DNA Technologies (IDT, Coralville, IA, USA). To create the CRISPR knockout cell lines, CRISPR ribonucleoprotein (RNP) complexes were delivered using cationic lipid transfection following the protocol entitled “Alt-R CRISPR-Cas9 System: cationic lipid delivery of CRISPR ribonucleoprotein complexes into mammalian cells,” which was provided by IDT. Briefly, to form the negative control sgRNA oligo, 1 µL each of 100 µM negative control crRNA (#1079138; IDT) and 100 µM tracrRNA (#1073189; IDT) from Alt-R^TM^ CRISPR-Cas9 Control Kit were added into 98 µL Nuclease-Free Duplex Buffer (#11-01-03-01; IDT), and the solution was incubated in 95 °C for 5 minutes before it was allowed to cool to 25 °C. Other sgRNA oligos were suspended in Nuclease-Free Duplex Buffer to make it 1 µM solution. Then, RNP complexes was formed by combining the sgRNA oligos, 1 µM Alt-R^TM^ S.p. Cas9-GFP V3 (#10008100; Integrated DNA Technoligies, Coralville, IA, USA), and Opti-MEM media (#31985062; Life Technologies), and incubating the solution in 25 °C for 5 minutes to allow assembly. The RNP complexes were then combined with RNAiMAX transfection reagent (#13778150; Thermo Fisher Scientific) and were incubated in 25 °C for 20 minutes to form the complexes, followed by transfection. The efficacy of transfection in each cell was determined based on the presence of GFP signals in cells within 24-48 hours after transfection. Cells were then plated in 100 mm dishes with 300 cells per dish, clones were grown for 7 – 9 days, and several colonies were selected and expanded. Confirmation of the successful knockouts was achieved with western blotting and gel electrophoresis of the cDNA library from each cell line. As a clarification, A549 was transfected with the negative control sgRNA and sgRNA 1.AD (Hs.Cas9.NFE2L2.1.AD) targeting NRF2 on the same day to create A549 sgCTRL1 and A549 NRF2 Complete KO cells, and was transfected with the negative control sgRNA and sgRNA 1.AE (Hs.Cas9.NFE2L2.1.AE) targeting NRF2 on other day to create A549 sgCTRL2 and A549 NRF2 Isoform Specific KO cells. NRF2 sgRNA 1.AE, which was utilized to generate the Isoform Specific KO cell line, does not target transcript isoforms NM.001313903.2 and NM.001313904.1 (**Fig 7A** **and S6**), which are translated into NRF2 isoform 5 and 6 (NP_001300832.1, 532 amino acids long, and NP_001300833.1, 505 amino acids long). These isoforms are thought to be truncated at the N-terminal domain that contains KEAP1-interacting Neh2 domain, but could retain the C-terminal basic leucine zipper domain, which supports the hypothesis that DNA-binding capacity is retained [NCBI RefSeq, NP_001300832.1, NP_001300833.1; [4]]. H1299T were transfected with negative control sgRNA, and sgRNA 1.AA (Hs.Cas9.KEAP1.1.AA) and 1.AD (Hs.Cas9.KEAP1.1.AD) targeting KEAP1 to create H1299T sgCTRL, H1299T KEAP1 KO1, and H1299T KEAP1 KO2 cells, respectively.

### Transcriptome Profiling by RNA-seq

RNA was isolated from H1299T, H1299T sgCTRL, H1299T KEAP1 KO1, H1299T KEAP1 KO2, A549, A549 sgCTRL1, A549 sgCTRL2, A549 NRF2 Complete KO, and A549 NRF2 Isoform Specific KO cells, utilizing RNeasy Plus Mini Kit (#74134; Qiagen, Hilden, Germany) and were sent for the quality control and mRNA-seq (sequencer: Element Aviti24 (short read), flow cell type: HIGH output, kit length: 150 cycle (2x75bp)) at Iowa Institute of Human Genetics, Carver College of Medicine. The data was mapped using the Kallisto pseudoalignment tool [41], and differential gene expression analysis was performed using Sleuth [42].

### Western blotting

Cells were pelleted and were subjected for step-by-step lysis of cells, using NE-PER™ Nuclear and Cytoplasmic Extraction Reagents (#78833; Thermo Fisher Scientific). Briefly, cytoplasmic components were separated from intact nuclei, and nuclear protein lysate was isolated from DNA and mRNA. 25 µg of cytoplasmic protein lysates and 15 µg of nuclear protein lysates were loaded onto a NuPAGE^TM^ 4-12% Bis-Tris Gel (Invitrogen, Eugene, Oregon, USA) and was run at 150 V for approximately 1.5 hours. Proteins were transferred to 0.2 µm Immun-Blot PVDF Membranes (#1620177; Bio-RAD, Hercules, CA, USA) for 1.5 hours at 200 mA. The membrane was then blocked with 5% milk before incubation with KEAP1 (D6B12) rabbit monoclonal antibody (mAb) (#8047; Cell Signaling Technology, Danvers, MA, USA) at [1:500], NRF2 (E3J1V) rabbit mAb (#33649; Cell Signaling Technology) at [1:1000], *β*-actin mAb (#3700; Cell Signaling Technology) at [1:3000], ALDH3A2 polyclonal antibody (pAb) (#15090-1-AP; Proteintech, Rosemont, IL) at [1:1000], ferritin heavy chain 1 (FtH1) mAb (#4393; Cell Signaling Technology) at [1:1000], glutamate-cysteine ligase catalytic subunit (GCLC) mAb (#52183; Cell Signaling Technology) [1:1000], glutamate-cysteine ligase modifier subunit (GCLM) mAb (Antibody Registry ID: AB_1553694; Developmental Studies Hybridoma Bank, deposited by Clinical Proteomics Technologies for Cancer, National Cancer Institute, and maintained at The University of Iowa, Department of Biology, Iowa City, IA 52242) at [1:1000], Heme oxygenase 1 (HMOX1) mAb (#NBP1-97507; Novus Biologicals, Centennial, CO, USA) at [1:1000], peroxiredoxin 1 (Prx1) mAb (#8499; Cell Signaling Technology) at [1:1000], TrxR1 mAb (# 6925; Cell Signaling Technology) at [1:500], thioredoxin (Trx1) mAb (#2429; Cell Signaling Technology) at [1:500], heterochromatin protein 1*α* (HP1*α*) mAb (#ab109028; Abcam) at [1:1000], and Lamin B1 mAb (# 13435; Cell Signaling Technology) at [1:1000] overnight at 4 °C. Following the incubation, the membrane was washed and was incubated in anti-rabbit or anti-mouse IgG HRP-linked secondary antibody at [1:5000] (#7074S and 7076S, respectively; Cell Signaling Technology) for 40 minutes at room temperature before visualization using SuperSignal^TM^ West Pico PLUS Chemiluminescent Substrate (#34580; Thermo Fisher Scientific).

### Seahorse-XF Real-time ATP Rate Assay

For H1299T sgCTRL, H1299T KEAP1 KO, A549 sgCTRL, and A549 NRF2 KO cells, Seahorse-XF Real-time ATP rate assay was performed as described previously with some modifications [43, 44]. Briefly, approximately 3000 – 6000 cells were plated in Seahorse XF Pro M Cell Culture Microplates (Agilent Technologies, Santa Clara, CA) and were allowed to grow for 48 – 72 hours in RPMI 1640 media supplemented with 10% FBS. Prior to the assay, growth medium was replaced with Seahorse assay media (Dubecco’s Modified Eagle’s Medium (DMEM) (#D5030; Sigma-Aldrich)) that was supplemented with 5 mM HEPES (#15630080; Gibco, Thermo Fisher Scientific), 10 mM D-glucose (#A2494001; Gibco, Thermo Fisher Scientific), 2 mM L-glutamine (#A2916801; Gibco, Thermo Fisher Scientific), and 1 mM sodium pyruvate (#11360070; Gibco, Thermo Fisher Scientific), with the pH adjusted to 7.4 at 37 °C, using an XF Prep station and incubator (Agilent Technologies). The supplemented assay media was prepared fresh and formulated to the proper pH right before the assay. Cells were then allowed to equilibrate and incubate for 30 – 45 minutes at 37 °C in the Prep Station CO_2_-free incubator. The assay consisted of sequential injections of 2.5 µM Oligomycin A (#75351; Sigma-Aldrich) and a combination of 2 µM each Rotenone (#R8875; Sigma-Aldrich) and Antimycin A (#A8674; Sigma-Aldrich). After calibration using Seahorse XF Pro Controller software, analysis was initiated in an Agilent-Seahorse XF Pro Bioenergetic Flux Analyzer (Agilent Technologies). Following the experiment, the cells were treated with 10 µM bisbenzimide H33342 trihydochloride (#B2261; Sigma-Aldrich) in the assay media and were analyzed using the BioTek Cytation 1 imaging reader for cell counts and normalization. Mitochondrial and glycolytic ATP production rates (amol (10^-18^) ATP/cell/s) were calculated based on the oxygen consumption rates (OCR) and proton efflux rate (PER), which was converted from extracellular acidification rate (ECAR), following the manufacturer’s protocol. Proper buffer factor and CO_2_ contribution factors were added into the Real-Time ATP Rate assay report generator, and a phosphate-to-oxygen ratio of 2.75 was used to calculate the mitochondrial ATP production rates.

### Measurement of mitochondrial membrane potential utilizing JC-1

H1299T sgCTRL and KEAP1 KO cells, and A549 sgCTRL and NRF2 KO cells were plated in 60 mm dishes and were allowed to grow for 48 – 72 hours to reach 1 – 1.5 million cells prior to the experiment. From the MitoProbe^TM^ JC-1 Assay Kit for Flow Cytometry (M34152; Life Technologies), utilizing the 30 µg JC-1 (5’,6,6’-tetrachloro-1,1’,3,3’-tetraethylbenzimidazolylcarbocyanine iodide) powder, 200 µM solution was made in DMSO (D2438; Sigma Aldrich). For the experiment, cells were treated with 2 µM JC-1 and 50 µM CCCP (Carbonyl cyanide m-chloro phenyl hydrazone; 500 µM solution was provided in the kit), the mitochondrial uncoupler, in fresh media, and were incubated at 37 °C for 15 minutes with intermittent inversion. After the incubation, cells were collected, centrifuged, and resuspended in PBS. Then cells were filtered through the Sheet Nylon 30 mesh (#146506; VWR International Incorporated, Radnor, PA, USA), collected in 12x75 mm Polystyrene Round-Bottom Tube (#352054; Coning Inc., Corning, NY, USA) for detection of fluorescence, utilizing FACScan (Waters Biosciences, Milform, MA, USA; excitation: 514 nm, emission: 529 nm (green) and 590 nm (red)

### Measurement of Mitochondrial Mass utilizing MitoTracker Green

H1299T sgCTRL and KEAP1 KO cells, and A549 sgCTRL and NRF2 KO cells were plated in 60 mm dishes and were allowed to grow for 48 – 72 hours to reach 1 – 1.5 million cells prior to the experiment. 1 mM MitoTracker Green (MTG) stock solution was prepared from a vial of MitoTracker^TM^ Green FM (M7514; Invitrogen) in DMSO and was diluted 1:10 with PBS to make working stocks. To make the staining buffer, 1 µL working stock was added per 1 mL PBS for the experiment. Cells were collected, centrifuged, and washed with PBS before staining. For staining, cells were resuspended in 500 µL PBS/MTG staining buffer and were incubated at 37 °C for 15 minutes with intermittent inversion. Cells were then placed on ice to stop the reaction, were filtered through the mesh, and were collected in 12x75 mm Polystyrene Round-Bottom Tube for detection of fluorescence, utilizing FACScan (excitation (Ex) maxima: 490 nm, emission (Em) maxima: 516 nm).

### Metabolomics profiling

A549 sgCTRL and A549 NRF2 KO cells were plated in 6-well cell culture plates and were allowed to grow for 48 – 72 hours to reach 0.8 – 1 million cells. Cells were then washed twice with ice-cold PBS and twice with ice-cold sterile ddH_2_O and were flash-frozen by floating the plates above the liquid nitrogen to preserve the metabolic profiles. The frozen plates were immediately placed on dry ice to maintain the temperature while being transported to Metabolomics Core Facility, Carver College of Medicine, for LC-MS profiling.

For sample processing, cells were scraped using 1 mL of ice-cold extraction solvent (acetonitrile:methanol:water, 2:2:1) spiked with 9 heavy internal standards. The scraped cells were transferred to 1.5 mL Eppendorf tubes and snap-frozen in liquid nitrogen. The frozen cell extracts were sonicated for 10 minutes, rotated at −20°C for 1 hour, and then centrifuged at the highest speed for 10 minutes. A 100 µL aliquot of the supernatant was transferred to a new microcentrifuge tube. The extracts were then dried using a SpeedVac apparatus. Dried extracts were reconstituted in 50 µL acetonitrile/water (1:1 v/v) vortexed well, kept in -20C overnight, centrifuged and the supernatant was transferred to LC-MS autosampler vials for analysis. LC-MS data were acquired on a Thermo Q Exactive hybrid quadrupole Orbitrap mass spectrometer with a Vanquish Flex UHPLC system. The LC column used was a Millipore SeQuant ZIC-pHILIC (2.1 x 150 mm, 5 µm particle size) with a ZIC-pHILIC guard column (20 x 2.1 mm). The LC column used was a Millipore SeQuant ZIC-pHILIC (2.1 x 150 mm, 5 µm particle size) with a ZIC-pHILIC guard column (20 x 2.1 mm). The injection volume was 8 µL.

Mobile phase is as follows: Solvent A: 20 mM ammonium carbonate [(NH4)_2_CO_3_] and 0.1% Ammonium Hydroxide (v/v) [NH_4_OH] [Note: pH is ∼9.1]; Solvent B: Acetonitrile. The method was run at a flow rate of 0.150 mL/min. The gradient starts at 80% B and decreasing to 20% B over 20 minutes; returning to 80% B in 0.5 minutes; and held there for 7 minutes [45]. There was a 2-minute pre-equilibration time prior to sample injection and during re-equilibration from 24.5 to 26.5 minutes flow is increased to 0.3 ml/min. The mass spectrometer was operated in full-scan, polarity-switching mode from 1 to 20 minutes, with the spray voltage set to 3.0 kV, the heated capillary held at 275 °C, and the HESI probe held at 350 °C. The sheath gas flow was set to 40 units, the auxiliary gas flow was set to 15 units, and the sweep gas flow was set to 1 unit. MS data acquisition was performed in a range of m/z 70–1,000, with the resolution set at 70,000, the AGC target at 1 × 10^6^, and the maximum injection time at 200 ms [45].

Acquired LC-MS data were processed by Thermo Scientific TraceFinder 5.2 software, and metabolites were identified based on the University of Iowa Metabolomics Core facility standard-confirmed, inhouse library. Signal drift was corrected by local polynomial regression against pooled QC samples [46]. The drift corrected data was normalized to the total ion current or to the total sum of metabolite values on a per sample basis, as specified, converted into log_2_ values, and then analyzed using GraphPad Prism® (GraphPad Software, Inc., San Diego, CA, USA).

### Measurements of Intracellular and Mitochondrial ROS

H1299T sgCTRL and H1299T KEAP1 KO cells, and A549 sgCTRL and A549 NRF2 KO cells were plated in 60 mm dishes and were allowed to grow for 48 – 72 hours to reach 1 – 1.5 million cells prior to the experiment. We referred to the vendor’s instructions and the previously published protocol [47] for CellROX Deep Red Staining, and for dihydroethidium (DHE)/MitoSOX oxidation [33]. 10 mM Antimycin A solution was prepared in DMSO for the positive control in DHE and MitoSOX Labeling Assay, and 72.24 mM tert-butyl hydroperoxide (TBH) (#458139; Sigma-Aldrich) for the positive control in CellROX Deep Red experiments. Cells were washed in DPBS and were collected, centrifuged, and resuspended in 500 µL of 5 mM sodium pyruvate (100 mM stock, #11360070; Life Technologies) containing DPBS or HBSS -/-/-(#14175095; Life Technologies) before staining with DHE/MitoSOX or CellROX Deep Red, respectively. For DHE/MitoSOX labeling, cells were incubated with 10 µM DHE or 2 µM MitoSOX, 10 µM Antimycin A, and DMSO (vehicles) in pyruvate containing DPBS. For CellROX Deep Red staining, cells were incubated in 5 µM CellROX Deep Red, and 400 µM TBH. All the incubations were in dark at 37 °C for 15 – 30 minutes with intermittent inversion. To stop the reaction, the stained samples were placed on ice, were filtered through the mesh, and were collected in 12x75 mm Polystyrene Round-Bottom Tube for detection of fluorescence. The samples were analyzed utilizing FACScan (DHE/MitoSOX Ex/Em: 488/585 (nm), CellROX Deep Red Ex/Em maxima: 640/665 (nm)). Median fluorescence intensity of 10000 cells was calculated in each cell line for statistics.

### Cell Cycle Analysis by Propidium Iodide (PI) staining

H1299T sgCTRL and H1299T KEAP1 KO cells, and A549 sgCTRL and A549 NRF2 KO cells were plated in 60 mm dishes and were allowed to grow for 48 – 72 hours to reach 1 – 1.5 million cells prior to the experiment. PI staining for cell cycle analysis was performed as previously described [48]. Briefly, cells were collected, and centrifuged, and washed in DPBS, before the fixation in 70% ethanol at least overnight. After fixation, cells were centrifuged and washed with DPBS. Samples were then incubated in 100 µL RNAse A solution (1mg/mL stock was made in ddH_2_O from #158153; Qiagen) for 30 minutes, followed by addition of 1 mL of 35 µg/mL stock PI (#P4170; Sigma-Aldrich) solution, incubation in room temperature for 1 hour in dark, and filtration through the mesh, and were collected in 12x75 mm Polystyrene Round-Bottom Tube for detection of fluorescence, utilizing Flow Cytometry. DNA quantity was measured by FACScan, and percentage of cells in G1, S, or G2 phases were calculated with CellQuest Pro (Waters Biosciences) and MODFIT software (Verity Software House, Topsham, ME, USA) and the data was utilized for statistics.

### Statistical Analysis

Quantification of western blot Images was performed using ImageJ (v1.53k; U.S. National Institutes of Health, Bethesda, Maryland, USA; https://imagej.net/ij/). Analysis of dose response curves, clonogenic survivals, quantification from western blot images, antioxidant enzyme activities, cell cycle, mitochondrial and glycolytic ATP production rates, % OXPHOS, % Glycolysis, JC-1 Red/Green fluorescence ratio, mitochondrial mass, intracellular and mitochondrial ROS, and metabolomics data were performed with GraphPad Prism. Concentrations of drugs were converted into log_10_ values before the quantification of area under curves was performed. The drift corrected data from metabolomics profiling was normalized to the total ion current or to the total sum of metabolite values on a per sample basis, as specified, converted into log_2_ values, and then analyzed using GraphPad Prism®. For two-group analysis, Welsh’s T-test (unequal variances) or Student’s T-test (equal variances) was applied. For analysis of three or more groups, statistical test was selected based on the scale of each dataset. For targeted experimental assays, one-way or two-way ANOVA with Tukey’s multiple comparison test was used to strictly control the family-wise error rate across all the pairwise comparisons. For high-dimensional metabolomics profiling, one-way ANOVA with original false discovery rate (FDR) method of Benjamini and Hochberg (FDR threshold: Q < 0.05) was used to account for large-scale multiple testing while preserving statistical power. For each experiment, *n* = number of biological replicates performed on different days. Errors represent ± SEM, and significance are shown as: *p ≤ 0.05, **p ≤ 0.01, ***p ≤ 0.001, ****p ≤ 0.0001. Outliers were removed utilizing ROUT (Q = 1%) method and were excluded from the statistical analysis. Further analysis of drug dose response data was performed using a web application, SynergyFinder+ [49]. Non-linear curve and goodness of fit tests were used for dose response curves. Irradiation dose response curves were generated using IGOR Pro (WaveMetrics, Inc., Portland, OR, USA), and significance are shown as *p ≤ 0.05, **p ≤ 0.01, ***p ≤ 0.001, ****p ≤ 0.0001.

## Results

### KEAP1 Mutant NSCLC cells and Normal Lung Epithelial Cells Show Enhanced Resistance to P-AscH^-^ or Auranofin, Relative to KEAP1 WTs

Inactivating mutations in *KEAP1* stabilize and constitutively activate NRF2, leading to upregulation of thioredoxin/glutathione metabolism, alteredaltered iron metabolism, and increased resistance to multiple therapeutic modalities [5–10]. Because *KRAS* and *STK11* mutations frequently co-occur with *KEAP1* mutations in NSCLCs [5, 50–52], the mutational status of KRAS, STK11, NRF2 were obtained from DepMap and are summarized in **Fig. S1** [53]. To determine the impact of KEAP1 status on treatment response, *KEAP1* mutant cell lines (A549, H2030, H460) and *KEAP1* WT cell lines (Calu-6, H1299, H1299T, H292) were treated with auranofin (AUR, 0-2 µM for 24 hr) or P-AscH^-^ (0-160 pmol/cell for 1 hr) followed by clonogenic survival assay under standard culture conditions (21 % O_2_, **Fig 1A-D**). In parallel, primary lung epithelial cells (HBEpC) were treated with P-AscH^-^ (0-160 pmol/cell for 1 hr) to provide a normal reference to A549 and H1299T NSCLCs under more physiologically relevant oxygen conditions (4% O_2_, **Fig S3A**). As expected, all KEAP1 mutant cells were more resistant to P-AscH^-^ and AUR, relative to KEAP1 WT at 21% O_2_ (**Fig. 1A-D**, as expected). Moreover, under 4% O_2_, both HBEpCs and KEAP1 mutant A549 remained more resistant to P-AscH^-^relative to H1299T (**Fig. S3AB**). Notably, both A549 and H1299T were more resistant to P-AscH^-^ at 4% vs. 21% O_2_, indicating that P-AscH^-^ cytotoxicity is oxygen dependent (**compare Fig1A to Fig S3A**). Consistent with the expected mechanism of action, auranofin inhibited TrxR activity in A549, H1299T, and HBEpCs (**Fig. 1E** and **Fig. S4**). These results demonstrate that KEAP1 mutant NSCLCs cells are comparatively more resistant to P-AscH^-^ induced cell killing, which is strongly influenced by oxygen tension, and HBEpCs were relatively more resistant (vs. NSCLCs) to the combination of P-AscH^-^ + AUR at physiologic oxygen conditions.

**Figure 1.**
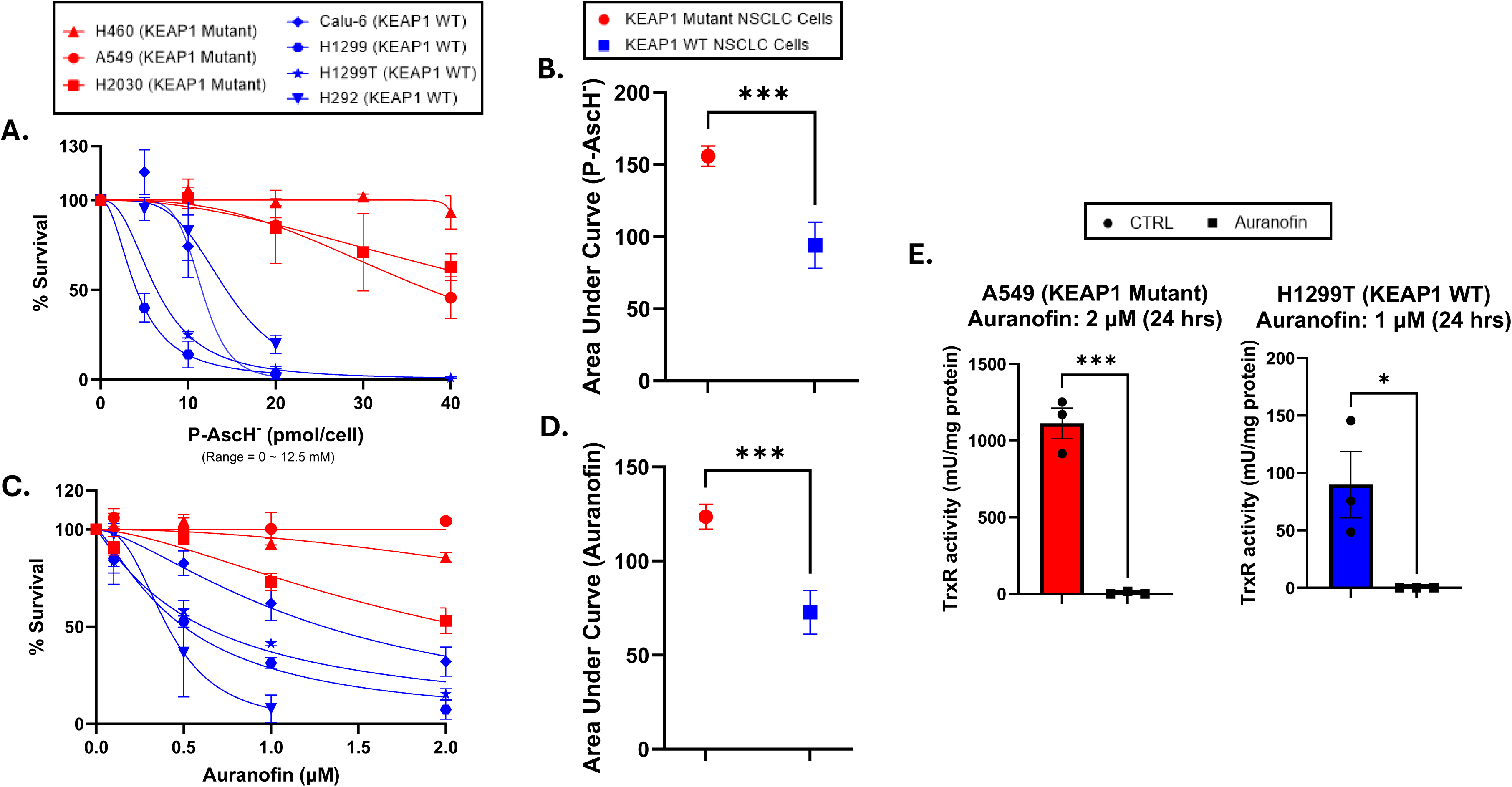
**KEAP1 mutant NSCLC cells show enhanced resistance to P-AscH^-^ and Auranofin, relative to KEAP1 WTs**. (A) Dose response curves of H460, A549, H2030, Calu-6, H292, H1299, and H1299T cells following 1-hour P-AscH^-^ treatments at the indicated concentrations. (B) Quantification of area under curve from P-AscH^-^ dose-response curves in KEAP1 mutant and WT NSCLC cells. (C) Dose response curves of H460, A549, H2030, Calu-6, H292, H1299, and H1299T cells following treatment with Auranofin for 24 hours at the indicated concentrations. (D) Quantification of area under curve from Auranofin dose-response curves in KEAP1 mutant and WT NSCLC cells. (E) TrxR activities in H1299T (KEAP1 WT) and A549 (KEAP1 Mutant) cells with or without Auranofin (1 – 2 µM, 24-hour treatment). Cells were maintained and treated in 5% CO_2_, 21% O_2_, and humidified in a 37°C incubator. Concentrations of P-AscH^-^ and Auranofin were converted to log_10_ values before the calculation of the area under curves of KEAP1 mutant and WT groups, representing overall sensitivity. Lower area under the curve values indicate higher sensitivity. Errors represent ± SEM, *n =* 3. *p ≤ 0.05, **p ≤ 0.01, ***p ≤ 0.001, ****p ≤ 0.0001 by Welch’s T-test. For TrxR activity assays, errors represent ± SEM, *n =* 3. *p ≤ 0.05, **p ≤ 0.01, ***p ≤ 0.001, ****p ≤ 0.0001 by Student’s T-test.

### Pretreatment with Auranofin Synergistically Sensitizes KEAP1 Mutant and WT NSCLC Cells but not Normal Lung Epithelial Cells to P-AscH^-^

To determine if auranofin pretreatment sensitizes the KEAP1 mutant and KEAP1 WT NSCLC cells to P-AscH^-^, seven of the NSCLC cells (A549, H2030, H460, Calu-6, H1299, H1299T, and H292) were pretreated with auranofin (0.5 - 5 µM for 24 hours) followed by treatment with P-AscH^-^ (5 - 160 pmol/cell for 1 hour) and clonogenic survival assay. Auranofin synergistically sensitized both KEAP1 mutant and KEAP1 WT NSCLC cells to P-AscH^-^ (**Fig. 2AC, and Fig. S3CD**), which was further confirmed through Loewe synergy scores of >10 from analysis of drug dose response data with SynergyFinder+ (**Fig. 2BD, S2**). In contrast, auranofin did not sensitize HBEpC to P-AscH^-^ (**Fig. S3E**). These findings demonstrate that the combination of auranofin and P-AscH^-^ enhanced sensitivity in NSCLCs regardless of their KEAP1 mutational status, while being less toxic to HBEpCs.

**Figure 2.**
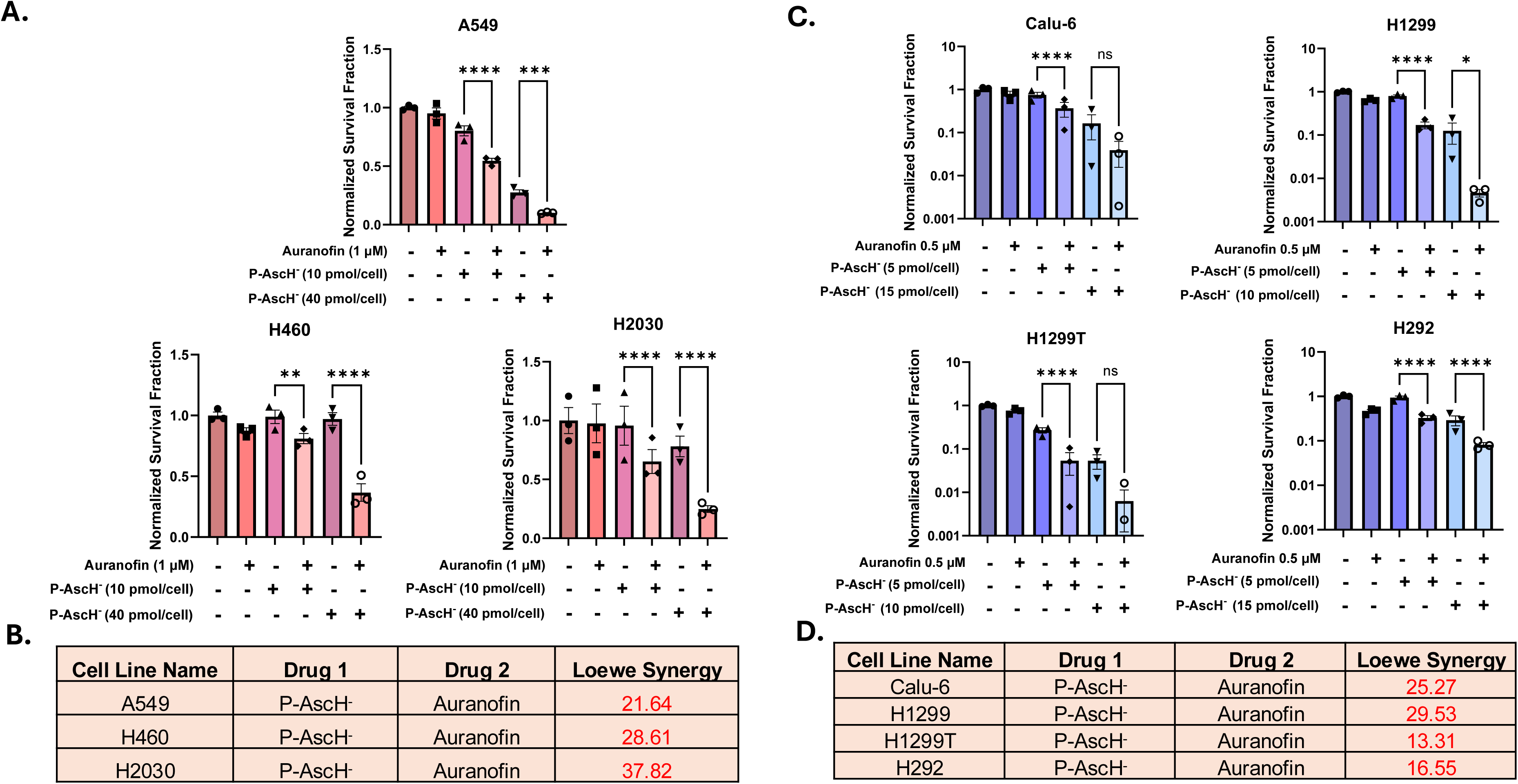
Pretreatment with Auranofin synergistically sensitizes KEAP1 Mutant and WT NSCLC cells to P-AscH^-^. (A) Clonogenic Survival assay of A549, H2030, and H460 cells after treatment with 1 µM auranofin for 24 hours followed by treatment with P-AscH^-^ (1 hour). (B) Loewe synergy scores representing the synergistic effect of Auranofin (1 μM) plus P-AscH^-^ (5 mM (10 pmol/cell), 12.5 mM (40 pmol/cell)) in A549, H460, and H2030 cells. (C) Clonogenic Survival assay of Calu-6, H1299, H1299T, and H292 cells after treatment with 0.5 µM auranofin for 24 hours followed by treatment with P-AscH^-^ (1 hour). (D) Loewe synergy scores representing the synergistic effect of Auranofin (0.5 μM) plus P-AscH^-^ (2.5 mM (5 pmol/cell), 6.25 mM (15 pmol/cell), and 7.5 mM (20 pmol/cell)) in Calu-6, H1299, H1299T, and H292 cells. Cells were maintained and treated in 5% CO_2_, 21% O_2_, in a humidified 37°C incubator. Errors represent ± SEM, *n* = 3. *p ≤ 0.05, **p ≤ 0.01, ***p ≤ 0.001, ****p ≤ 0.0001 by Two-Way ANOVA. Synergistic scores below -10 are considered antagonistic, the scores from -10 to 10 are considered additive, and the scores above 10 are considered synergistic interactions [49].

### N-Acetyl-L-Cysteine and Catalase Overexpression Inhibit the Cytotoxicity of Auranofin Combined with P-AscH^-^

To determine whether thiol oxidation and H₂O₂ contribute to the cytotoxic effects of combined auranofin and P-AscH^-^ treatment, we employed both chemical and enzymatic approaches to modulate cellular redox stress. N-acetyl-L-cysteine (NAC), a thiol-containing antioxidant and precursor for glutathione synthesis, was used to replenish intracellular thiols and limit oxidative stress. A549CAT2 and H1299TCAT15 cells were generated previously to express doxycycline-inducible human catalase, allowing selective scavenging of intracellular H₂O₂ following doxycycline treatment [25, 26]. A549 (KEAP1 mutant) and H1299TCAT15 (KEAP1 WT) cells were treated with NAC for 3 hours prior to exposure to auranofin (0.5 µM or 1µM1µM, 21 hours) and P-AscH^-^ treatments (5 and 40 pmol/cell, 1 hour) followed by clonogenic survival assays. **Fig 3AE** shows that NAC treatment significantly attenuated the cytotoxicity of P-AscH^-^ + AUR combination in both cell lines. Consistent with protection against oxidative thiol stress, NAC also prevented the treatment-induced increases in % glutathione disulfide (%GSSG) (**Fig 3BF**). To directly investigate the role of H_2_O_2_, A549CAT2 and H1299TCAT15 cells, were treated with 1 µg/mL doxycycline to induce catalase expression 24 hours prior to auranofin + P-AscH^-^ treatments followed by clonogenic survival and catalase activity assays. Catalase overexpression significantly inhibited clonogenic cell killing induced with auranofin + P-AscH^-^ in A549CAT2 and H1299TCAT15 (**Fig. 3CDEH**). Interestingly, NAC pretreatment reduced %GSSG levels in AUR + P-AscH^-^-treated A549 and H1299TCAT15 cells without altering total GSH content, suggesting that the protective effect of NAC is attributable to suppression of thiol oxidation rather than changes in overall glutathione abundance. (**Fig S5A-F**). Overall, these data support the hypothesis that auranofin + P-AscH^-^ sensitizes both WT and mutant KEAP1 NSCLCs through both a thiol-oxidation and H_2_O_2_-dependent mechanism.

**Figure 3.**
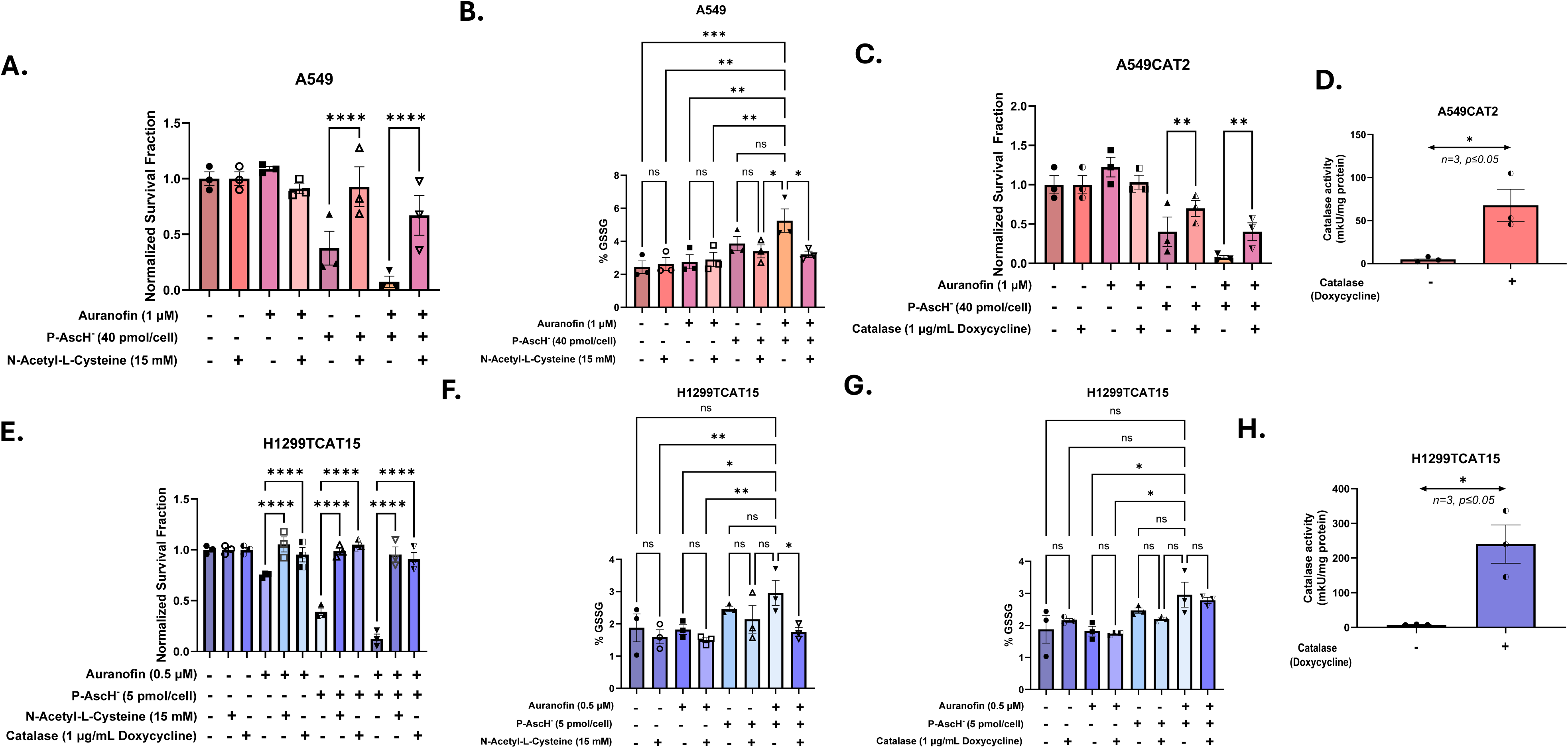
N-Acetyl-L-Cysteine and Catalase overexpression inhibits the toxicity of Auranofin combined with P-AscH^-^. (A) Clonogenic Survival assay of A549 cells after treatment with 15 mM NAC for 24 hours and 1 µM auranofin for 21 hours, followed by treatment with AscH^-^ (1 hour). (B) %GSSG in A549 cells that were treated with NAC (15 mM, 24 hours), Auranofin (1 µM, 21 hours), and P-AscH^-^ (40 pmol/cell, 1 hour). (C) Clonogenic Survival assay of A549CAT2 cells after treatment with 1 µg/mL doxycycline for 48 hours and 1 µM Auranofin for 21 hours, followed by treatment with P-AscH^-^ (1 hour). (D) Catalase activity measured after 48 hours of doxycycline treatment in A549CAT2 by catalase activity assay. (E) Clonogenic Survival assay of H1299T cells after treatment with 1µg/mL doxycycline for 48 hours, 15 mM NAC for 24 hours and 1 µM Auranofin for 21 hours, followed by treatment with P-AscH^-^ (1 hour). (F and G) %GSSG in H1299TCAT15 cells that were treated with NAC (15 mM, 24 hours) or doxycycline (1 µg/mL, 48 hours), Auranofin (0.5 µM, 21 hours), and P-AscH^-^ (5 pmol/cell, 1 hour). (H) Catalase activity measured after 48 hours of doxycycline treatment in H1299TCAT15 by catalase activity. For Clonogenic Survival assay and total and oxidized GSH activity assay, errors represent ± SEM, *n =* 3. *p ≤ 0.05, **p ≤ 0.01, ***p ≤ 0.001, ****p ≤ 0.0001 by Two-Way ANOVA. For catalase activity assay, errors represent ± SEM, *n =*3. *p ≤ 0.05, **p ≤ 0.01, ***p ≤ 0.001, ****p ≤ 0.0001 by Student’s T-test.

CRISPR-mediated Knockout of KEAP1 in H1299T Results in Enhanced Resistance to P-AscH^-^ and Irradiation but not to Auranofin, while CRISPR-mediated Knockout of NRF2 in A549 Results in Sensitization to All the Treatments Since KEAP1 mutant NSCLCs (i.e. A549), also carry KRAS and STK11 mutations that also regulate metabolic pathways and antioxidant expression [54–57], the relative contribution of loss of KEAP1 function and NRF2 function was determined using CRISPR-mediated KEAP1 KO in KEAP1 WT NSCLCs (H1299T) as well as CRISPR-mediated NRF2 KO in KEAP1 mutant NSCLCs (A549). Dose response experiments were performed after P-AscH^-^ or auranofin treatment, or after ionizing radiation in CRISPR-mediated KO of KEAP1 (H1299T) or NRF2 (A549) and appropriate guide RNA controls (sgCTRL) (**Fig 4 and 7**).

**Figure 4.**
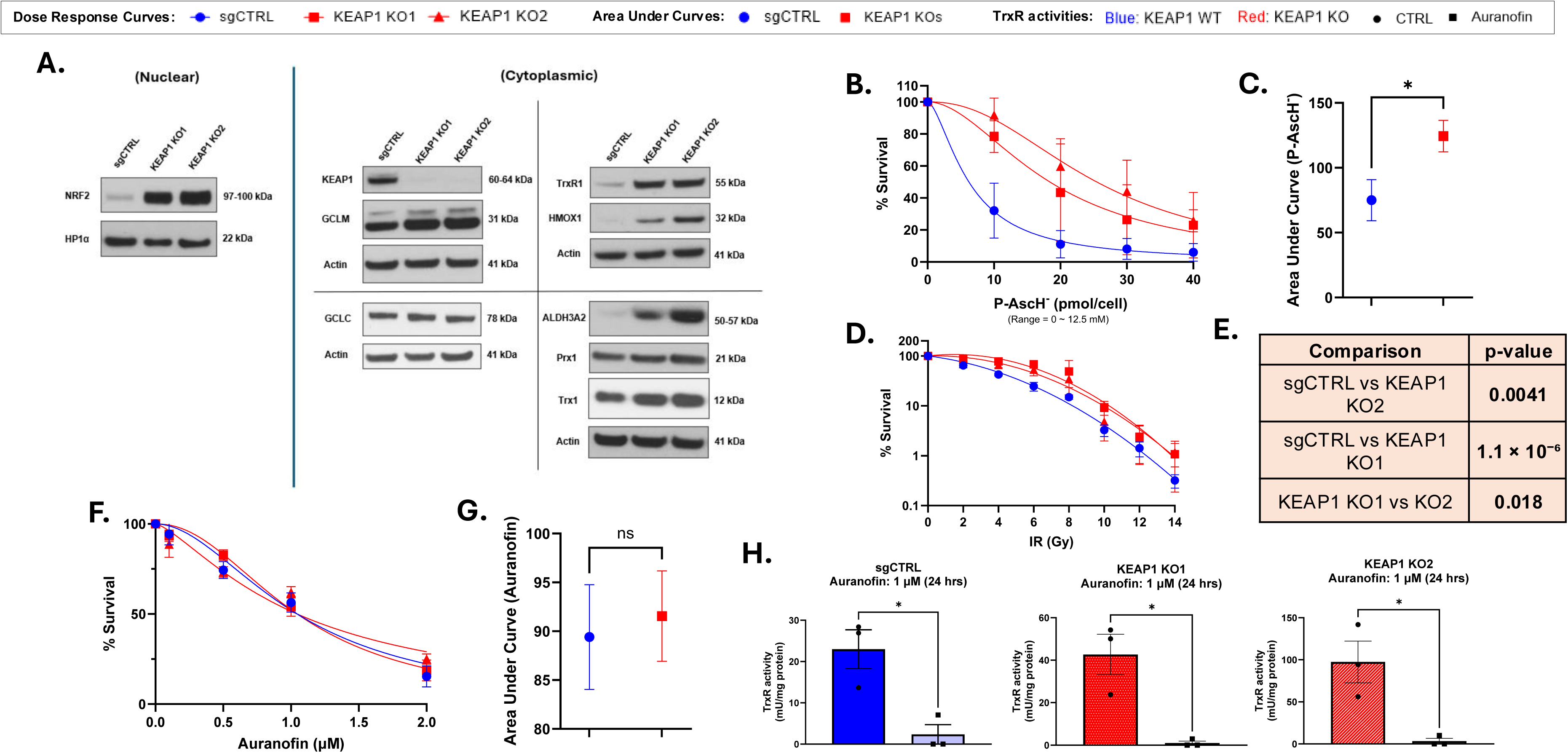
knockout (KO) of KEAP1 in H1299T results in enhanced resistance to P-AscH^-^ and radiation, but not to Auranofin. (A) Western Blot confirmation of KEAP1 KO and increased expressions of NRF2 and its targets. (B) Dose response curves of H1299T negative control and H1299T KEAP1 KO cells after treatment with P-AscH^-^ for 1 hour at the indicated concentrations. (C) Quantification of area under curve from P-AscH^-^ dose-response curves in H1299T negative controls and H1299T KEAP1 KO cells. (D) Dose response curves of H1299T negative controls and H1299T KEAP1 KO cells after treatment after irradiation of indicated doses. (E) The statistical analysis of the curve performed using IGOR Pro. (F) Dose response curves of H1299T negative control and H1299T KEAP1 KO cells after treatment with Auranofin for 24 hours at the indicated concentrations. (G) Quantification of area under curve from Auranofin dose-response curves in H1299T negative controls and H1299T KEAP1 KO cells. (H) TrxR activities in H1299T KEAP1 KO cells with or without Auranofin 1 µM (24-hour treatment). Cells were maintained and treated in 5% CO_2_, 21% O_2_, and humidified in a 37°C incubator. For area under curves, concentrations of Auranofin and P-AscH^-^ were converted to log_10_ values, and area under curve was calculated for each group, representing overall sensitivity to the drugs. Errors represent ± SEM, *n =* 3. *p ≤ 0.05, **p ≤ 0.01, ***p ≤ 0.001, ****p ≤ 0.0001 by Welch’s T-test. The statistical analysis of the radiation dose response curves were performed using IGOR Pro. Errors represent ± SEM, *n* = 3 – 4, *p ≤ 0.05, **p ≤ 0.01, ***p ≤ 0.001, ****p ≤ 0.0001. For TrxR activity assays errors represent ± SEM, *n =* 3. *p ≤ 0.05, **p ≤ 0.01, ***p ≤ 0.001, ****p ≤ 0.0001 by Student’s T-test.

KEAP1 KOs (KO1 and KO2) demonstrated loss of KEAP1 protein and increased nuclear localization of NRF2 (**Fig 4A**), and NRF2 KOs showed loss of NRF2 in the nuclear extracts (**Fig 7A**). The expression of NRF2 targets (GCLC, GCLM, TrxR1, ALDH3A2, HMOX1, Prx1, Trx1) were assessed across the cell lines using western blotting and transcriptome analysis (**Fig 4A, 7A, S7, and S8**). Based on western blotting, A549 NRF2 KO3 cells retained modest residual NRF2 expression (**Fig 7A**) and retained some expression of NRF2 targets, relative to the complete KO (KO2) (**Fig 7A** **and S8**). Transcriptome analysis on both A549 and HT1299T parental cell lines along with KEAP1 targeted (H1299T) clones, NRF2 targeted clones (A549), and their respective sgRNA controls were performed (**Table S1**). Differential gene expression analysis was performed in each cell line by comparing expression between two independently targeted clones with parental and control populations (**Fig. S7**). Deletion of KEAP1 in H1299T cells significantly altered the expression of 192 genes (FDR ≤ 0.001, **Fig. S7a**, **Table S2**). NRF2 deletion in A549 cells was associated with differential expression of 1736 genes (FDR ≤ 0.001, **Fig. S7b**, **Table S3**). Comparison of these two gene sets identified a signature of 31 genes that showed increased expression with KEAP1 deletion and decreased expression with NRF2 loss (**Fig. S7c**). As expected, this gene signature is enriched with genes involved with cellular redox homeostasis (GO:0045454, FDR = 6.12e-07), oxidoreductase activity (GO:0016491, FDR = 4.12e-07).

Additionally, H1299T KEAP1 KO cells (KO1 and KO2) demonstrated similar plating efficiency, doubling times, and cell cycle distribution, relative to H1299T sgCTRL cells, (**Fig S9 A-D, Fig S10 A,B**). Bur interestingly, H1299T KEAP1 KO cells were significantly more resistant to clonogenic cell killing by P-AscH^-^ and radiation, but not to auranofin (**Fig 4B-G)** relative to H1299T sgCTRL cells (**Fig S9 A-D, Fig S10 A,B**). These results demonstrate that loss of KEAP1 in H1299T does not affect cell growth but can increase NRF2 activity in the nucleus and recapitulate much of the gene expression and resistance phenotypes seen in KEAP1 mutant A549.

In contrast, A549 cells with either complete (KO2) or isoform specific (KO3) NRF2 knockout demonstrated different plating efficiencies, doubling time, and cell cycle distribution (**Fig S9 E-H, Fig S10 C, D**.). Both clones were significantly more sensitive to auranofin, P-AscH^-^, and radiation, relative to their respective isoform specific sgCTRLs (sgCTRL1 and sgCTRL2) (**Fig 7B-G**), with KO2 being more sensitive to all agents relative to KO3. Interestingly, when looking at NRF2 target genes, the complete NRF2 KO2 showed greater reductions in TrxR1, HMOX1, GCLC, GCLM, ALDH3A1, and Prx1, relative to the isoform specific KO3. Auranofin treatments (**Fig. 4H and 7H**) uniformly inhibited TrxR activity but did not consistently inhibit GPx1 activity in either KOs (**Fig. S13**). Overall, these results support the hypothesis that complete KO of NRF2 in KEAP1 mutant cells significantly decelerates the growth characteristics of the cells, reduces expression of NRF2 target genes and activities, and sensitizes the cells to P-AscH^-^, AUR, and radiation. Furthermore, the isoform specific NRF2 knockout cells were more resistant than the complete KO, as well as showing less effects on cell growth, expression of NRF2 targets, and less sensitivity to P-AscH^-^ and AUR, suggesting that NRF2 isoform 5/6 is functional. Overall, these results support the hypothesis that KEAP1-NRF2 pathway promotes resistance to auranofin, P-AscH^-^, and radiation.

### Pretreatment with Auranofin Synergistically Sensitizes H1299T sgCTRL and H1299T KEAP1 KO cells to P-AscH^-^

To further assess if auranofin pretreatment sensitizes NSCLC cells to P-AscH^-^ regardless of KEAP1 status, H1299T sgCTRL and H1299T KEAP1 KO cells were pretreated with auranofin (0.5 µM for 24 hours) followed by treatment with P-AscH^-^ (5 - 40 pmol/cell for 1 hour) and clonogenic survival assay. Auranofin synergistically sensitized both H1299T sgCTRL and H1299T KEAP1 KO cells to P-AscH^-^ (**Fig. 5AB**), which was further confirmed through Loewe synergy scores of >10 from analysis of drug dose response data with SynergyFinder+ (**Fig. 5C** **and S11AB**). These findings further support the hypothesis that auranofin + P-AscH^-^ may improve responses to radiotherapy in NSCLCs regardless of their KEAP1 mutational status.

**Figure 5.**
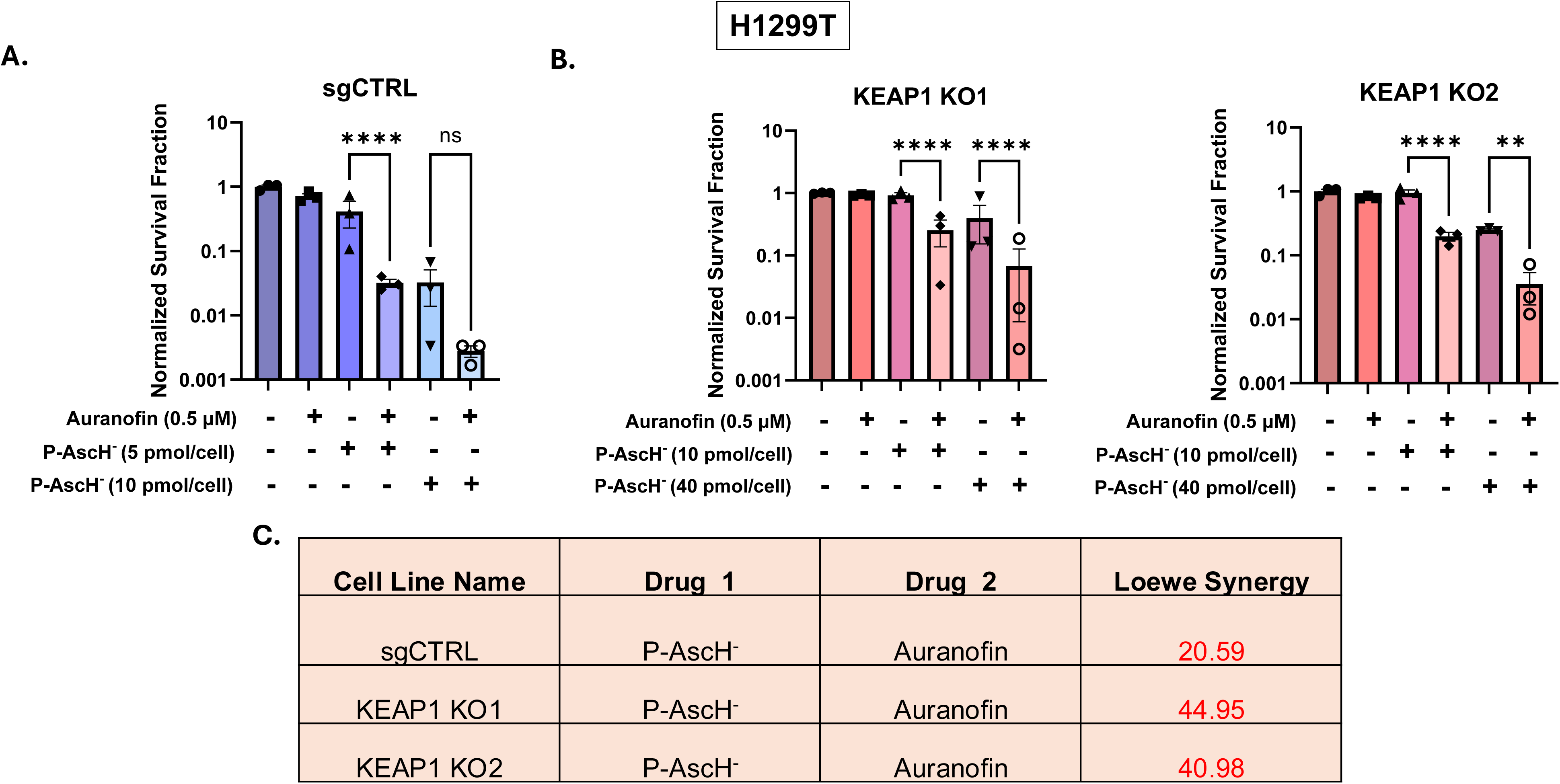
Pretreatment with Auranofin synergistically sensitizes H1299T sgCTRL and H1299T KEAP1 KO cells to P-AscH^-^. (A) Clonogenic Survival Assays of H1299T sgCTRL cells after treatment with 0.5 µM Auranofin for 24 hours followed by treatments with P-AscH^-^ (1 hour). (B) Clonogenic Survival Assays of H1299T KEAP1 KO1 or KO2 cells after treatment with 0.5 µM Auranofin for 24 hours followed by treatments with P-AscH^-^ (1 hour). (C) Loewe synergy scores representing the synergistic effect of Auranofin (0.5 μM) plus P-AscH^-^ (2.5 mM (5 pmol/cell), 5 mM (10 pmol/cell), and12.5 mM (40 pmol/cell)). Cells were maintained and treated in 5% CO_2_, 21% O_2_, and humidified in a 37°C incubator. Errors represent ± SEM, *n =* 3. *p ≤ 0.05, **p ≤ 0.01, ***p ≤ 0.001, ****p ≤ 0.0001 by Two-Way ANOVA. Synergistic scores below -10 are considered antagonistic, the scores from -10 to 10 are considered additive, and the scores above 10 are considered synergistic interactions [49].

### P-AscH^-^ Synergistically Sensitizes H1299T sgCTRL Cells and H1299T KEAP1 KO Cells to Ionizing Radiation

To assess if P-AscH^-^ pretreatment sensitizes NSCLC cells to radiation regardless of KEAP1 status, H1299T sgCTRL and H1299T KEAP1 KO cells were pretreated with P-AscH^-^ (5 - 30 pmol/cell for 1 hour) followed by radiation (2 or 4 Gy) and clonogenic survival assay. The timeline of the treatments is shown in **Figure S12C**. P-AscH^-^ sensitized both H1299T sgCTRL and H1299T KEAP1 KO cells to radiation (**Fig. 6AB**), which was further confirmed through Loewe synergy scores of >10 from analysis of drug dose response data with SynergyFinder+ (**Fig. 6C** **and S12AB**).

**Figure 6.**
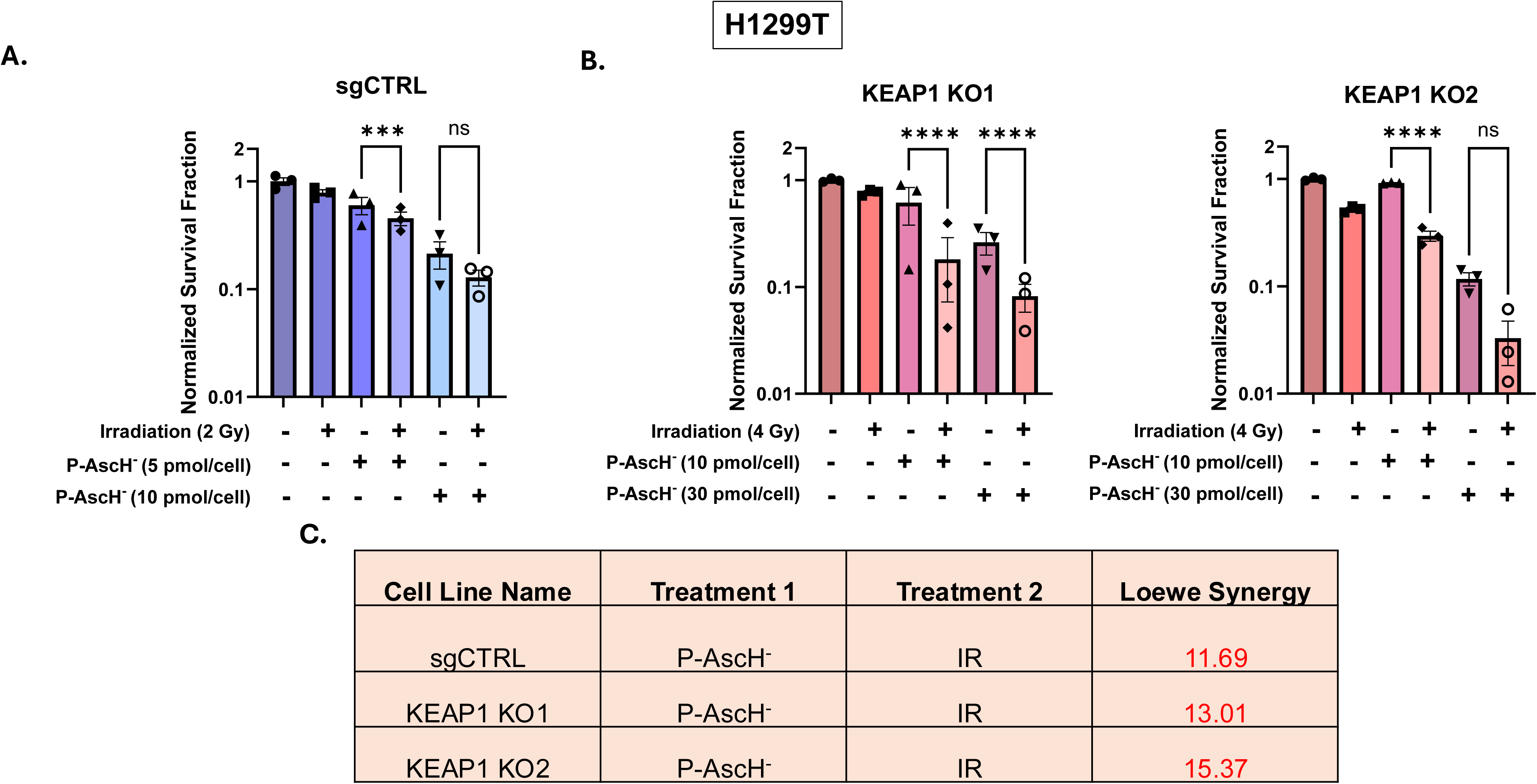
P-AscH^-^ synergistically sensitizes H1299T sgCTRL cells and H1299T KEAP1 KO cells to ionizing radiation. (A) Clonogenic Survival assay of H1299T negative control cells after treatments with P-AscH^-^ (1 hour) and 2 Gy irradiation (X-ray). (B) Clonogenic Survival assay of H1299T KEAP1 KO cells after treatments with P-AscH^-^ (1 hour) and 4 Gy irradiation (X-ray). (C) Loewe synergy scores representing the synergistic effect of P-AscH^-^ (2.5 mM (5 pmol/cell), 5 mM (10 pmol/cell), and10 mM (30 pmol/cell)) plus radiation (IR, 2 or 4 Gy)). Cells were maintained and treated in 5% CO_2_, 21% O_2_, and humidified in a 37°C incubator. Errors represent ± SEM, *n =* 3. *p ≤ 0.05, **p ≤ 0.01, ***p ≤ 0.001, ****p ≤ 0.0001 by Two-Way ANOVA. Synergistic scores below -10 are considered antagonistic, the scores from -10 to 10 are considered additive, and the scores above 10 are considered synergistic interactions [49].

### CRISPR-mediated KO of KEAP1 or NRF2 in NSCLC cells lead to changes in total and oxidized glutathione contents, GPx1 activity, and TrxR activity

To identify if thiol antioxidants play a role in changes in response to the therapies in KEAP1 KO and NRF2 KO cells, total and GSSG, GPx1 activity, and TrxR activity was determined. H1299T KEAP1 KO cells showed increases in total GSH and GSSG levels, and at least one KEAP1 KO cell line showed significant increase in GPx1 or TrxR activities (**Fig. 8A-F**). Additionally, A549 NRF2 KO cells showed significant decreases in total GSH levels, with that of isoform specific KO cells being significantly greater than that of complete KO cells (**Fig. 8G**). The total GSSG level was not significantly changed in A549 sgCTRL and NRF2 KO cells, but GSH/GSSG and %GSSG were significantly greater in A549 NRF2 Complete KO cells, compared to A549 sgCTRL cells and A549 NRF2 isoform specific KO cells (**Fig. 8H-J**). The similar trends in total GSH levels and % GSSG were observed in the metabolomics data in which drift corrected raw data was normalized to total ion current or total sum of metabolites radiometrically for analysis (**Fig. 10J-L** **and Fig. S17J-L**). Moreover, GPx1 activity was significantly lower in A549 NRF2 Complete KO cells, compared to A549 sgCTRL cells and slightly reduced in A549 NRF2 Isoform Specific KO cells (**Fig. 8K**), and TrxR activities were lower in both A549 NRF2 KO cells (**Fig. 8L**). These results provide evidence that KEAP1-NRF2 pathway mediates resistance to auranofin, P-AscH^-^ and radiation through thiol-and peroxide-dependent metabolism.

**Figure 7.**
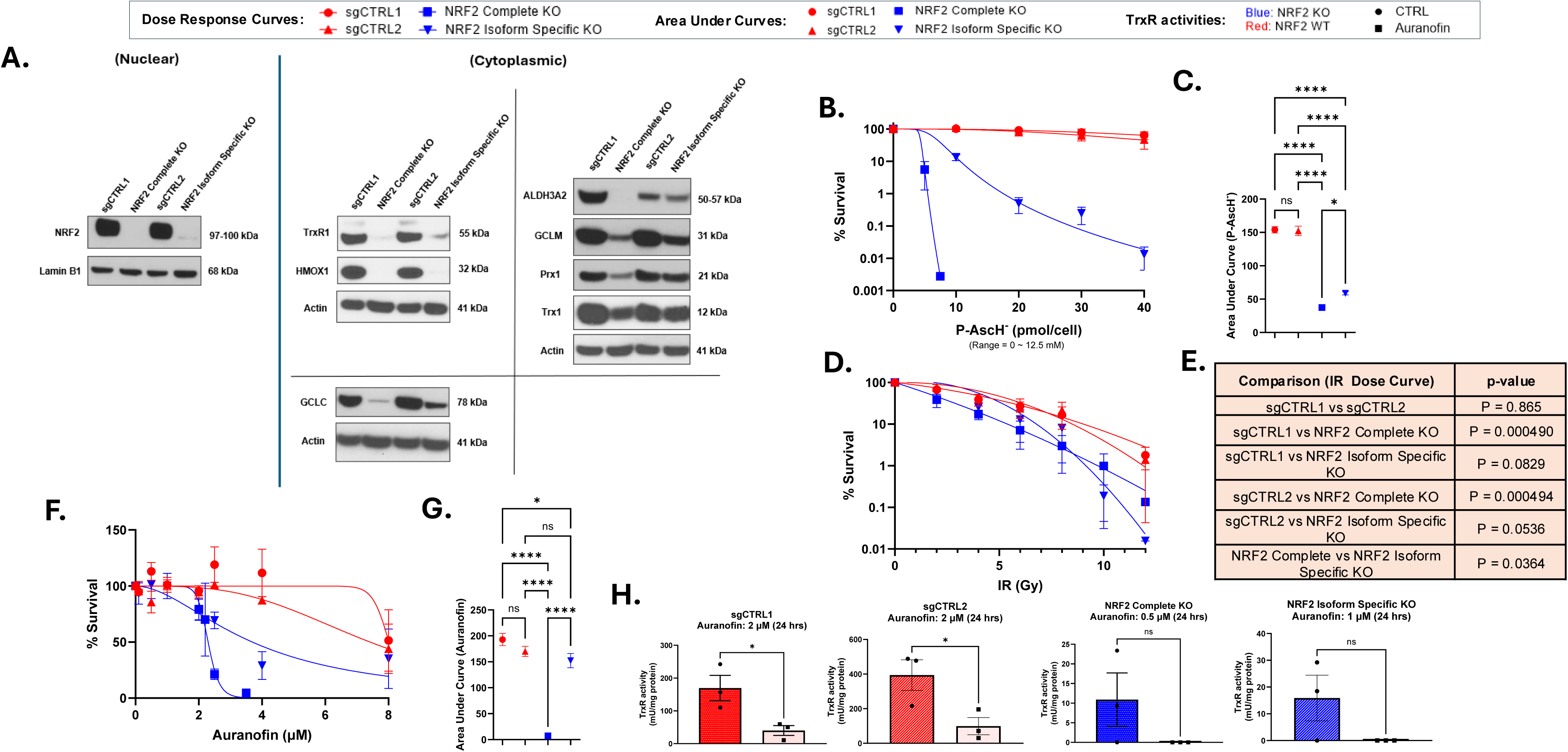
KO of NRF2 in A549 results in enhanced sensitivity to P-AscH^-^, Auranofin, and radiation. (A) Western Blot confirmation of NRF2 KO and increased expressions of NRF2 targets. (B) Dose response curves of A549 negative controls and A549 NRF2 KO cells after treatment with P-AscH^-^ for 1 hour at the indicated concentrations. (C) Quantification of area under curve from P-AscH^-^ dose-response curves in H1299T negative controls and A549 NRF2 KO cells. (D) Dose response curves of A549 negative controls and A549 NRF2 KO cells after irradiation of indicated doses. (E) The statistical analysis of the radiation dose response curves. (F) Dose response curves of A549 negative controls and A549 NRF2 KO cells after treatment with Auranofin for 24 hours at the indicated concentrations. (G) Quantification of area under curve from Auranofin dose-response curves in A549 negative controls and A549 NRF2 KO cells. (H) TrxR activities in A549 sgCTRL and NRF2 KO cells with or without Auranofin 0.5 - 2 µM (24-hour treatment). Cells were maintained and treated in 5% CO_2_, 21% O_2_, and humidified in a 37°C incubator. For the dose response curves, concentrations of Auranofin and P-AscH^-^ were converted to log_10_ values before the area under curves were calculated for each group, representing overall sensitivity to the drugs. Errors represent ± SEM, *n* = 3. *p ≤ 0.05, **p ≤ 0.01, ***p ≤ 0.001, ****p ≤ 0.0001 by Welch’s T-test. The statistical analysis of the radiation dose response curves were performed using IGOR Pro. Errors represent ± SEM, *n* = 3 – 4, *p ≤ 0.05, **p ≤ 0.01, ***p ≤ 0.001, ****p ≤ 0.0001. For TrxR activity assays errors represent ± SEM, *n =* 3. *p ≤ 0.05, **p ≤ 0.01, ***p ≤ 0.001, ****p ≤ 0.0001 by Student’s T-test.

**Figure 8.**
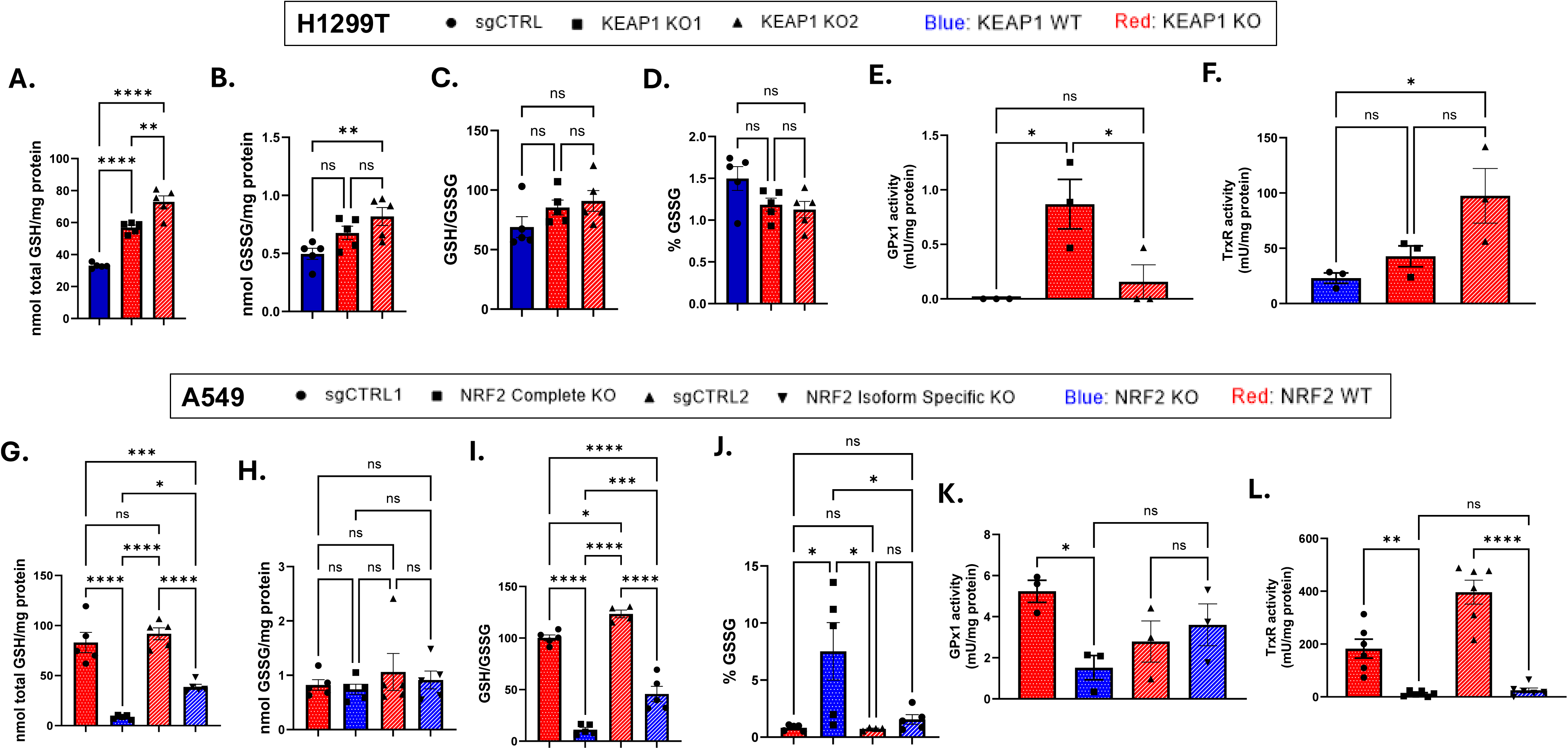
KO of KEAP1 or NRF2 in NSCLC cells lead to changes in total and oxidized glutathione contents, GPx1 activity, and TrxR activity. (A) Total GSH levels in H1299T sgCTRL and KEAP1 KO cells. (B) GSSG levels in H1299T sgCTRL and KEAP1 KO cells. (C) GSH/GSSG ratios in H1299T sgCTRL and KEAP1 KO cells. (D) %GSSG in H1299T sgCTRL and KEAP1 KO cells. (E) GPx1 activity in H1299T sgCTRL and KEAP1 KO cells. (F) TrxR activities in H1299T sgCTRL and KEAP1 KO cells. (G) Total GSH levels in A549 sgCTRL and NRF2 KO cells. (G) GSSG levels in A549 sgCTRL and NRF2 KO cells. (H) GSH/GSSG ratios in A549 sgCTRL and NRF2 KO cells. (I) GSH/GSSG ratios in A549 sgCTRL and NRF2 KO cells. (J) %GSSG in A549 sgCTRL and NRF2 KO cells. (K) GPx1 activities in A549 sgCTRL and NRF2 cells. (L) TrxR activities in A549 sgCTRL and NRF2 KO cells. Cells were maintained and treated in 5% CO_2_, 21% O_2_, and humidified in a 37°C incubator. Errors represent ± SEM, For total and oxidized GSH assays, *n =* 4 – 5, For GPx1 activity assays, *n =* 3, and for TrxR activity assays, *n =* 3 – 6, *p ≤ 0.05, **p ≤ 0.01, ***p ≤ 0.001, ****p ≤ 0.0001 by One-way ANOVA. Outliers were detected and removed from the analysis using ROUT (Q = 1 %) method.

**Figure 9.**
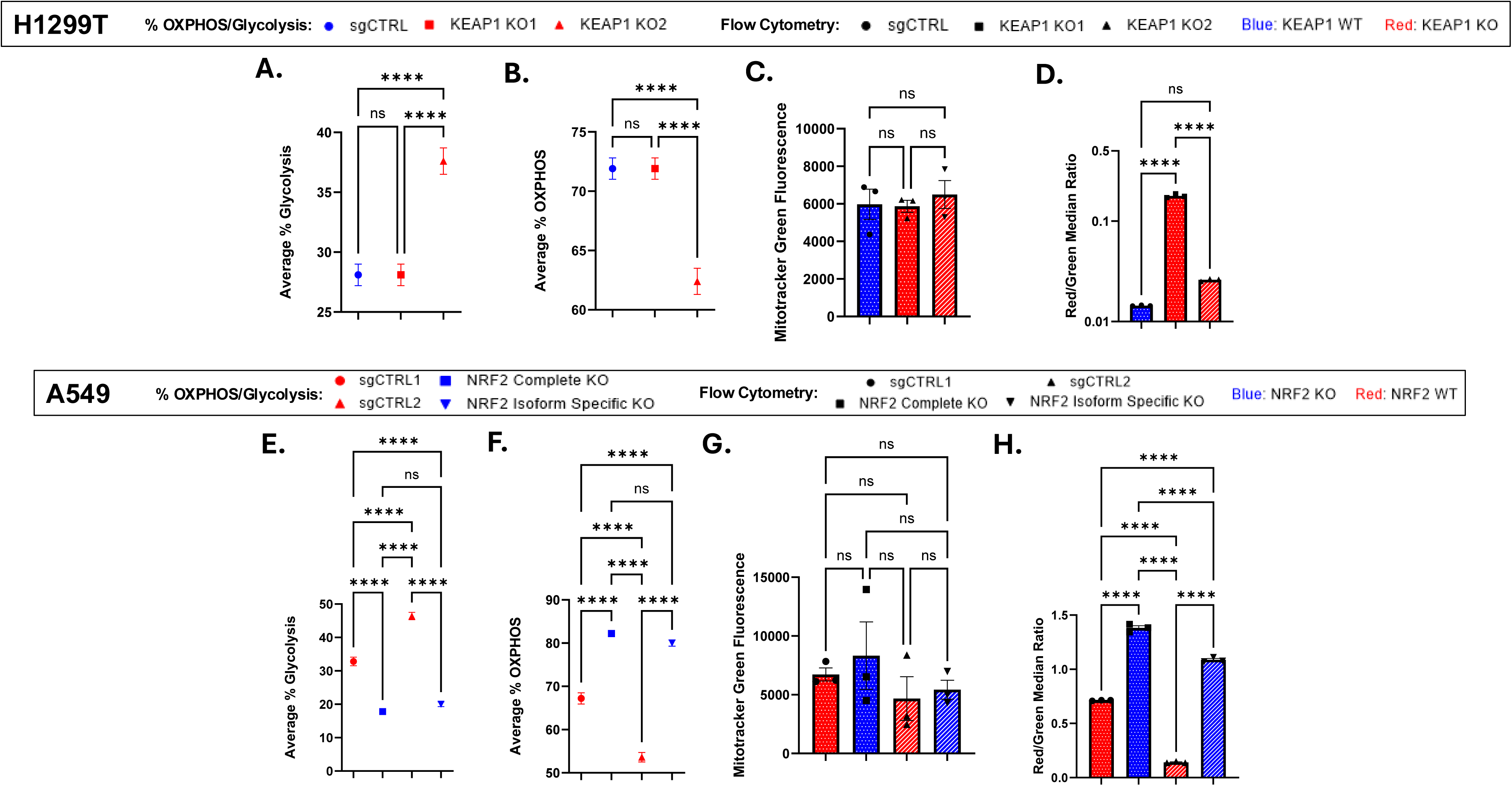
CRISPR-mediated KO of KEAP1 or NRF2 in NSCLC cells cause changes in mitochondrial membrane potential, glycolysis and oxidative phosphorylation, while causing no changes in quantity of mitochondria. (A and B) Average % Glycolysis and Average % OXPHOS in H1299T sgCTRL and H1299T KEAP1 KO cells. (C) Quantification of MitoTracker Green fluorescent H1299T sgCTRL and H1299T KEAP1 KO cells. (D) Quantification of the ratio of red and green JC-1 fluorescence in H1299T sgCTRL and H1299T KEAP1 KO cells. (E and F) Average % Glycolysis and Average % OXPHOS in A549 sgCTRL and NRF2 KO cells. (G) Quantification of MitoTracker Green fluorescent A549 sgCTRL and A549 NRF2 KO cells. (H) Quantification of the ratio of red and green JC-1 fluorescent in A549 sgCTRL and NRF2 KO cells. Errors represent ± SEM, *n* = 3. *p ≤ 0.05, **p ≤ 0.01, ***p ≤ 0.001, ****p ≤ 0.0001 by One-way ANOVA.

**Figure 10.**
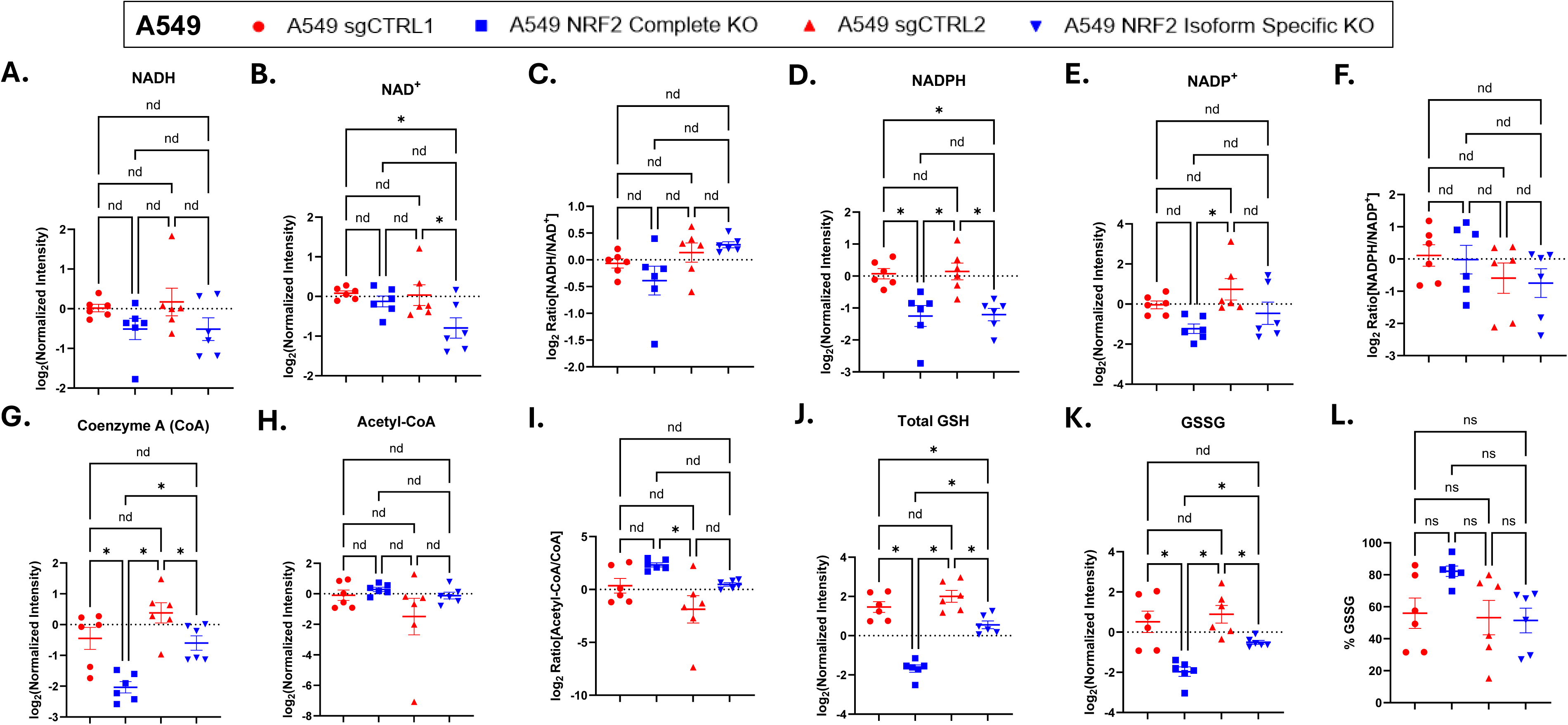
KO of NRF2 in A549 results in decrease in NADPH, NADP^+^, Coenzyme A, total GSH land GSSG evels, and increase in Acetyl-CoA/Coenzyme A ratio and % GSSG, but not in NADH and NAD^+^ levels, and NADH/NAD^+^ and NADPH/NADP^+^ ratios. (A) The log_2_-transformed normalized NADH levels in A549 sgCTRL and A549 NRF2 KO cells. (B) The log_2_-transformed normalized NAD^+^ levels in A549 sgCTRL and A549 NRF2 KO cells. (C) The log_2_-transformed NADH/NAD^+^ ratios in A549 sgCTRL and A549 NRF2 KO cells. (D) The log_2_-transformed normalized NADPH levels in A549 sgCTRL and A549 NRF2 KO cells. (E) The log_2_-transformed normalized NADP^+^ levels in A549 sgCTRL and A549 NRF2 KO cells. (F) The log_2_-transformed NADPH/NADP^+^ ratios in A549 sgCTRL and A549 NRF2 KO cells. (G) The log_2_-transformed normalized Coenzyme A (CoA) levels in A549 sgCTRL and A549 NRF2 KO cells. (H) The log_2_-transformed normalized Acetyl-CoA levels in A549 sgCTRL and A549 NRF2 KO cells. (I) The log_2_-transformed Acetyl-CoA/CoA in A549 sgCTRL and A549 NRF2 KO cells. (J) The log_2_-transformed total GSH levels in A549 sgCTRL and A549 NRF2 KO cells. (K) The log_2_-transformed GSSG levels in A549 sgCTRL and A549 NRF2 KO cells. (L) % GSSG in A549 sgCTRL and A549 NRF2 KO cells. Cell samples were submitted to Metabolomics Core Facility, Carver College of Medicine, Iowa City, IA 52242 for LC-MS profiling. Raw data was drift corrected, the values of the metabolites were normalized to total ion current as a proxy of biomass per sample, and log_2_-transformed to stabilize variance and normalize the distributions. Errors represent ± SEM, *n* = 5 - 6. *p ≤ 0.05, **p ≤ 0.01, ***p ≤ 0.001, ****p ≤ 0.0001 by One-way ANOVA. To control for multiple comparisons, p-values from post-hoc pairwise comparisons were adjusted using the Benjamini-Hochberg False Discovery Rate (FDR) procedure. The FDR threshold of Q < 0.05 was used to determine the statistical significance. Furthermore, outliers were detected and removed from the analysis using ROUT (Q = 1 %) method.

### CRISPR-mediated KO of KEAP1 or NRF2 in NSCLC Cells Cause Changes in Mitochondrial Membrane Potential, Glycolysis, and Oxidative Phosphorylation, but not in Mitochondrial Mass or Intracellular Prooxidant Levels

In previous studies, cancer cells were shown to cope with increased ROS production, compared to normal cells, by upregulating glycolysis and the pentose phosphate pathway (PPP) to supply critical reductants, particularly NADPH, for peroxide removal and antioxidant activities [58–63]. Moreover, the KEAP1-NRF2 pathway is known to not only upregulate some enzymes that are involved in PPP and thereby NADPH production but also upregulate glycolytic enzymes and increases in glycolytic flux [10, 64, 65]. To assess if CRISPR-mediated KO of KEAP1 and NRF2 induce differences in ATP production rate and changes in metabolism, Seahorse-XF Real-time ATP Rate Assay was performed. At least one H1299T KEAP1 KO cell line showed a decrease in average % oxidative phosphorylation (OXPHOS) and an increase in average % glycolysis relative to total cellular ATP production rate (**Fig. 9AB**), and the opposite trend was observed in A549 NRF2 KO cells compared to A549 sgCTRL cells (**Fig. 9EF**). The real-time changes in OCR and ECAR, which are essential for ATP rate calculation, are shown in **Figure S15**. To determine if these differences come from changes in mitochondrial mass or mitochondrial membrane potential, we performed MitoTracker Green Staining and JC-1 labeling (for mitochondrial membrane potential) followed by Flow Cytometry. No difference in MitoTracker Green fluorescence was observed among the H1299T sgCTRL and H1299T KEAP1 KO cells, or A549 sgCTRL and A549 NRF2 KO cells (**Fig. 9CG**), indicating similar mitochondrial mass. In contrast, JC-1 Red/Green median fluorescence ratio was significantly higher in one of the H1299T KEAP1 KO cell lines compared H1299T sgCTRL cells, while the difference in the values was only around 0.16, and there was no significant difference between the other H1299T KEAP1 KO cell line and H1299T sgCTRL cells (**Fig. 9D**). The Red/Green ratios were consistently and significantly higher in A549 NRF2 KO cells compared to A549 sgCTRL cells (0.6 – 1 in values) and was significantly higher in A549 NRF2 complete KO cells as well as in A549 NRF2 isoform specific KO cells (**Fig. 9H**). Interestingly, there were no differences in intracellular and mitochondrial prooxidant production in H1299T KEAP1 KO cells compared to the H1299T sgCTRL cells, and in A549 NRF2 KO cells compared to the A549 sgCTRL cells (**Fig. S14**). And notably, the metabolomics data in which metabolite values were normalized to total ion current on a per sample basis showed no consistent changes in Acetyl-CoA and nicotinamide adenine dinucleotide (NAD^+^ and NADH) levels among the A549 sgCTRL and A549 NRF2 KO cells, while A549 NRF2 KO cells showed decreases in Coenzyme A (CoA) and nicotinamide adenine dinucleotide phosphate (NADP^+^ and NADPH) levels, with NRF2 complete KO cells showing more significant decreases in NADP^+^ level compared to NRF2 isoform specific KO cells (**Fig. 10A-I**). Notably, total metabolite value sum normalization on a per sample basis of the drift corrected metabolites values resulted in comparable trends while both NRF2 KO cell lines showing significant decrease in NADP^+^ (**Fig. S17**). The results from metabolomics analysis indicate that NRF2 KO cells potentially upregulated OXPHOS as the result of adaptation mechanisms, through changes in mitochondrial enzyme activity that lead to increased consumption of CoA into Acetyl-CoA. In the previous studies, isoform 3 of NAD kinase and NDUFS7 (the core subunit of mitochondrial complex I) were suggested to be upregulated by NRF2 activity, and it was also suggested that NRF2 suppresses ATP-citrate lyase activity, which catalyzes production of Acetyl-CoA from CoA while producing oxaloacetate from citrate at the same time [66–68]. However, more studies are needed to identify the mechanisms of upregulation of OXPHOS in NRF2 KO NSCLC cells. Overall, the results suggest that KEAP1-NRF2 pathway plays an important role in maintaining the capacity of cancer cells to utilize glycolysis instead of OXPHOS to cope with the intracellular peroxide production.

## Discussion

Previous studies with NSCLCs showed that auranofin (through inhibition of TrxR) and P-AscH^-^ (through H_2_O_2_-and iron-based mechanisms) induce cancer cell specific toxicity [13, 19, 69]. However, these studies did not compare NSCLCs and normal human lung epithelial cells (HBEpCs) with auranofin + P-AscH^-^ or comparing KEAP1 mutant and WT cells with physiologically relevant 24-hour AUR dose responses. The results in the current report showed that 160 pmol/cell (∼40 mM) P-AscH^-^ was non-toxic to HBEpCs and KEAP1 mutant NSCLCs at physiological O_2_ concentration (4% O_2_) but were cytotoxic to KEAP1 WT NSCLCs. Furthermore, KEAP1 mutant and WT NSCLCs at physiological concentration (4% O_2_) were sensitized to P-AscH^-^ with auranofin pretreatment, but sensitization was not seen in HBEpCs.

In addition, at 21% O_2,_ all KEAP1 mutant NSCLCs (relative to KEAP1 WT) were resistant to P-AscH^-^ and AUR but both phenotypes demonstrated synergistic sensitization when treated with AUR + P-AscH^-^ demonstrating that KEAP1 mutations leading to NRF2 activation is a biomarker for resistance to ascorbate that can be overcome by combining with an FDA approved agent for combined modality cancer therapy development. Furthermore, the mechanism for how these agents act synergistically, was demonstrated to involve thiol oxidation and the formation of H_2_O_2_ as evidenced by the protection seen with NAC pretreatment and doxy-induced overexpression of catalase. These findings support the original hypothesis that KEAP1-NRF2 pathway regulates resistance to auranofin, P-AscH^-^, and radiation through thiol-and H_2_O_2_-based mechanisms in NSCLCs. In addition, our findings suggest that these two FDA approved drugs can be used as adjuvants for radio-chemotherapy in NSCLCs. However, in NSCLCs including A549, H2030, and H460, KEAP1 mutations frequently co-occur with mutations of KRAS and STK11 that regulate metabolic pathways and antioxidant expression [54–57].

To further identify if relative contributions of KEAP1 and NRF2 in mediating resistance to auranofin, P-AscH^-^ and radiation, CRISPR-mediated KEAP1 KO (in isogenic KEAP1 WT NSCLCs) or NRF2 KO cells (in KEAP1 mutant NSCLS) were generated and treated with auranofin, P-AscH^-^, and radiation dose responses, relative to sgCTRL and the CRISPR-mediated KOs. Western blot and transcriptome analysis demonstrated successful CRISPR-mediated targeting of each gene. Additionally, glutathione, GPx1, and TrxR analysis showed that CRISPR-mediated KO of KEAP1 in H1299T (KEAP1 WT) resulted in upregulation of nuclear NRF2 as well as antioxidant expression and activity. Similar analysis in the A549 NRF2 complete KO demonstrated significant downregulation of glutathione, GPx1, and TrxR metabolism which resulted in compromised cell growth and increased %GSSG demonstrating stress in redox metabolism. Furthermore, in A549 NRF2 KO that retained the isoform 5/6, showed less downregulation of glutathione, GPx1, and TrxR metabolism as well as no effect on cell growth supporting the hypothesis that isoform 5/6 was partially functional in reversing the effects of the complete KO.

H1299T KEAP1 KO cells showed increased resistance to P-AscH^-^ and radiation but not to auranofin, in comparison to H1299T sgCTRL cells (**Fig. 4B-G**). Survival of H1299T KEAP1 KO cells were approximately 1.4–2.1-fold less than that of A549, A549 sgCTRL, H2030, and H460 cells under the same dosages of P-AscH^-^, suggesting the KEAP1 deletion alone cannot recapitulate the phenotype of the KEAP1 mutant cells tested. However, this suggests that mutations in STK11 and KRAS may contribute to the resistance phenotype independent of upregulation of NRF2, but this hypothesis will be tested in future studies (**Fig. S1**).

A549 NRF2 KO1 cells that showed significant effects on cell growth were more sensitive to auranofin, P-AscH^-^, and radiation in comparison to A549 sgCTRLs. Again, consistent with the hypothesis that NRF2 isoform 5/6 retained significant function, NRF2 KO2 cells were more resistant to auranofin, P-AscH^-^, and radiation, relative to NRF2 Complete KO1 (**Fig. 7B-G**). As is identified in **Figure 7A** and **Figure S6**, A549 NRF2 Isoform Specific KO cells retained residual expression of NRF2, which is theoretically NRF2 isoform 5 and 6 that is suggested to be missing the N-terminal regulatory domain that interacts with KEAP1 and controls the stability, but retains the basic-leucine zipper domain in C-terminal, which plays an important role in DNA binging. These isoforms have been poorly studied, and to our knowledge, this is the first report supporting the hypothesis that these isoforms retain NRF2 function, relative to the complete KO of NRF2.

Additionally, auranofin sensitized both H1299T sgCTRL and KEAP1 KO cells to P-AscH^-^, and P-AscH^-^ as well as sensitizing the combinations with radiation **(Fig. 5 and 6**). These results further provided evidence that supports the original hypothesis. In future studies, H1299T sgCTRL, H1299T KEAP1 KO, A549 sgCTRL, and A549 NRF2 KO cells will be treated with auranofin, P-AscH^-^, and radiation and chemotherapeutic agents *in vivo* to test the hypothesis that these combinations could be used effectively to enhance therapeutic response. In addition, KEAP1 KO cells with Tet-On system for conditional induction of KEAP1 expression, and A549 NRF2 KO cells with Tet-On system for conditional induction of NRF2 expression will also be generated to test if restored expression of KEAP1 or NRF2 can reverse the biology seen in the CRISPR KOs. Furthermore, Phase II clinical trials in stage IIIBNSCLC (XACT-LUNG) utilizing Radio-chemotherapy + P-AscH^-^ are ongoing (NCT02905591), and genetic profiles of the patients will be analyzed in future studies to further identify the relative value of these mutations in therapeutic strategies for NSCLCs.

Surprisingly, in spite of the observed significant upregulation of thiol antioxidants levels and activities in H1299T KEAP1 KO cells compared to H1299T sgCTRL cells and the significant decrease in these in A549 NRF2 KO cells compared to A549 sgCTRL cells, intracellular and mitochondrial ROS levels were not significantly different in H1299T KEAP1 KO cells compared to H1299T sgCTRL cells, or A549 NRF2 KO cells compared to A549 sgCTRL cells (**Fig. S14**). Additionally, there were no differences in the quantity of mitochondria (**Fig. 9CG**). But interestingly, consistent increases in mitochondrial membrane potential were identified in A549 NRF2 KO cells compared to A549 sgCTRL cells, while only one KEAP1 KO cell line showed increases in mitochondrial membrane potential (**Fig. 9DH**). A549 NRF2 KO cells showed increased % OXPHOS compared to A549 sgCTRL cells, while at least one H1299T KEAP1 KO cell line showed increased % Glycolysis compared to H1299T sgCTRL cells (**Fig. 9ABEF**). Moreover, the metabolomics data showed no consistent changes in Acetyl-CoA and nicotinamide adenine dinucleotide (NAD^+^ and NADH) levels among the A549 sgCTRL and A549 NRF2 KO cells, while A549 NRF2 KO cells showing decrease in Coenzyme A (CoA) and nicotinamide adenine dinucleotide phosphate (NADP^+^ and NADPH) levels, with NRF2 complete KO cells showing more significant decrease in NADP^+^ level compared to NRF2 isoform specific KO cells (**Fig. 10A-I**). Moreover, we also performed total sum normalization of the drift corrected raw rata radiometrically for the metabolomics analysis, which yielded comparable trends while both NRF2 KO cell lines showing significant decrease in NADP^+^ (**Fig. S17**). The consistency of trends across both methods of normalization indicates that the observed metabolic profiles are robust to sample-wise scaling variations. Overall, these results suggest that KEAP1-NRF2 pathway plays an important role in maintaining the capacity of cancer cells to utilize glycolysis more than OXPHOS to cope with peroxide production. In our future studies, we will compare the expressions of ATP-Citrate lyase, NAD kinases, and NDUFS7 by western blot, and genetic manipulations that induce loss, suppression, or induction of these factors will be performed to further assess underlying mechanisms of the metabolic profiles of H1299T sgCTRL, H1299T KEAP1 KO, A549 sgCTRL, and A549 NRF2 KO.

This study also showed that KEAP1-NRF2 pathway may cause changes in cell growth and survival, and cell cycle progression (**Fig. S9 and S10**). Despite we have shown that the expression andactivity of the thiol antioxidants that are involved in peroxide removal in cells change significantly in KEAP1 KO and NRF2 KO NSCLC cells compared to their sgCTRL cells, more studies need to be pursued to identify if thiol antioxidant and peroxide cause the difference in survival and growth that were seen in A549 NRF2 Complete KO cells compared to A549 sgCTRL cells. In the future studies, clonogenic survival assay with A549 sgCTRL1 and A549 Complete KO cells will be performed, in which both cell lines would be incubated with NAC and PEGylated Catalase (#C4963; Sigma-Aldrich) after plating for 7 – 9 days to assess if providing these will cause changes in clonogenic survivals. Additionally, the growth rate of H1299T sgCTRL, H1299T KEAP1 KO, A549 sgCTRL, and A549 NRF2 KO cells will be compared *in vivo* mouse model.

## Conclusion

This study provides strong evidence supporting the hypothesis that the KEAP1-NRF2 pathway regulates resistance to auranofin, P-AscH^-^, and radiation through thiol-and H_2_O_2_-based mechanisms in NSCLCs. The results support the hypothesis the FDA approved auranofin and P-AscH^-^ could be used in combined modality therapies overcome KEAP1-NRF2-mediated redox-dependent resistance to radio-chemotherapy in NSCLC by targeting thiol and peroxide metabolism.

SS, MAF, MF, BGA, EBT, AJD, DRS: Conceptualization; SS, AMT, BAW, DLM, SAW, ANL, MQMG, TJN, CFP: Data curation; SS, AMT, DLM, BAW, SAW, ANL, MQMG, TJN, CFP, BGA, TIAE, AJD: Formal analysis; DRS: Funding acquisition; SS, AMT, BAW, DLM, SAW, ANL, MQMG, TJN, CFP: Investigation; SS, EBT, AJD, DRS: Methodology; AJD, DRS: Project administration; DRS, AJD: Resources; DRS, AJD: Supervision; DRS, AJD: Validation; SS, AJD, DRS: Visualization; SS, AJD, DRS: Roles/Writing - original draft; and SS, AMT, BAW, DLM, SAW, ANL, MQMG, TJN, CFP, FAM, TIAE, FM, BGA, EBT, AJD, DRS: Writing - review & editing.

## Acknowledgements

The Authors would like to thank Radiation and Free Radical Research Core at the University of Iowa for providing P-AscH^-^, ionizing radiation services, and antioxidant analysis. Additionally, the authors would like to thank Iowa Institute of Human Genetics at the University of Iowa for the RNA sequencing service, and Metabolomics Core at the University of Iowa for the metabolomics analysis.

## Funding

This work was supported by P01 CA217797, T32 CA078586, R01 CA298628, R01 DK138664, P50 CA302572, P30 CA086862, and the Holden Comprehensive Cancer Center.

^1^Glucose-6-phosphate (G-6-P), 6-phosphogluconate (6-P-G), glucose-6-phosphate dehydrogenase (G6PD), nicotinamide adenine dinucleotide phosphate (NADP^+^/NADPH), glutathione (GSH), oxidized glutathione (GSSG), glutathione peroxidase (GPx), glutathione reductase (GR), thioredoxin (Trx), peroxiredoxin (Prx), Kelch-like ECH-associated protein 1 (KEAP1), nuclear factor erythroid 2-related factor (NRF2), non-small cell lung cancer (NSCLC), human bronchial epithelial cells (HBEpC), Pharmacological ascorbate (P-AscH^-^), iron (Fe^3+^/Fe^2+^), hydrogen peroxide (H_2_O_2_), Organic peroxide (ROOH), hydroxyl radical (HO^●^), alkoxyl radical (RO^●^) thioredoxin reductase (TrxR), N-Acetyl-L-Cystein (NAC), doxycycline (doxy), Catalase (CAT), ionizing radiation (IR), single guide (sg), knockout (KO), aldehyde dehydrogenase 3A2 (ALDH3A2), glutamate-cysteine ligase catalytic subunit (GCLC), glutamate-cysteine ligase modifier subunit (GCLM), heme oxygenase (HMOX1), RNA Sequencing (RNA-Seq), oxygen consumption rate (OCR), extracellular acidification rate (ECAR), proton efflux rate (PER), oxidative phosphorylation (OXPHOS), MitoTracker Green (MTG), dihydroethidium (DHE), dimethyl sulfoxide (DMSO), propidium iodide (PI), Coenzyme A (CoA), nicotinamide adenine dinucleotide (NAD^+^/NADH), chromosome (chr)

## References

[1] R.L. Siegel, T.B. Kratzer, N.S. Wagle, H. Sung, A. Jemal, Cancer statistics, 2026, CA Cancer J Clin 76(1) (2026) e70043. 10.3322/caac.70043.

[2] R. Chen, R. Manochakian, L. James, A.G. Azzouqa, H. Shi, Y. Zhang, Y. Zhao, K. Zhou, Y. Lou, Emerging therapeutic agents for advanced non-small cell lung cancer, J Hematol Oncol 13(1) (2020) 58. 10.1186/s13045-020-00881-7.

[3] C. Putzu, S. Canova, P. Paliogiannis, R. Lobrano, L. Sala, D.L. Cortinovis, F. Colonese, Duration of Immunotherapy in Non-Small Cell Lung Cancer Survivors: A Lifelong Commitment?, Cancers (Basel) 15(3) (2023). 10.3390/cancers15030689.

[4] M.C. Jaramillo, D.D. Zhang, The emerging role of the Nrf2-Keap1 signaling pathway in cancer, Genes Dev 27(20) (2013) 2179–91. 10.1101/gad.225680.113.

[5] A. Di Federico, A. De Giglio, C. Parisi, F. Gelsomino, STK11/LKB1 and KEAP1 mutations in non-small cell lung cancer: Prognostic rather than predictive?, Eur J Cancer 157 (2021) 108–113. 10.1016/j.ejca.2021.08.011.

[6] Y. Jeong, J.A. Hellyer, H. Stehr, N.T. Hoang, X. Niu, M. Das, S.K. Padda, K. Ramchandran, J.W. Neal, H. Wakelee, M. Diehn, Role of KEAP1/NFE2L2 Mutations in the Chemotherapeutic Response of Patients with Non-Small Cell Lung Cancer, Clin Cancer Res 26(1) (2020) 274–281. 10.1158/1078-0432.CCR-19-1237.

[7] Y. Jeong, N.T. Hoang, A. Lovejoy, H. Stehr, A.M. Newman, A.J. Gentles, W. Kong, D. Truong, S. Martin, A. Chaudhuri, D. Heiser, L. Zhou, C. Say, J.N. Carter, S.M. Hiniker, B.W. Loo, Jr., R.B. West, P. Beachy, A.A. Alizadeh, M. Diehn, Role of KEAP1/NRF2 and TP53 Mutations in Lung Squamous Cell Carcinoma Development and Radiation Resistance, Cancer Discov 7(1) (2017) 86–101. 10.1158/2159-8290.CD-16-0127.

[8] E.B. Krall, B. Wang, D.M. Munoz, N. Ilic, S. Raghavan, M.J. Niederst, K. Yu, D.A. Ruddy, A.J. Aguirre, J.W. Kim, A.J. Redig, J.F. Gainor, J.A. Williams, J.M. Asara, J.G. Doench, P.A. Janne, A.T. Shaw, R.E. McDonald Iii, J.A. Engelman, F. Stegmeier, M.R. Schlabach, W.C. Hahn, Correction: KEAP1 loss modulates sensitivity to kinase targeted therapy in lung cancer, Elife 6 (2017). 10.7554/eLife.33173.

[9] D. Marinelli, M. Mazzotta, S. Scalera, I. Terrenato, F. Sperati, L. D’Ambrosio, M. Pallocca, G. Corleone, E. Krasniqi, L. Pizzuti, M. Barba, S. Carpano, P. Vici, M. Filetti, R. Giusti, A. Vecchione, M. Occhipinti, A. Gelibter, A. Botticelli, F. De Nicola, L. Ciuffreda, F. Goeman, E. Gallo, P. Visca, E. Pescarmona, M. Fanciulli, R. De Maria, P. Marchetti, G. Ciliberto, M. Maugeri-Sacca, KEAP1-driven co-mutations in lung adenocarcinoma unresponsive to immunotherapy despite high tumor mutational burden, Ann Oncol 31(12) (2020) 1746–1754. 10.1016/j.annonc.2020.08.2105.

[10] C. Tonelli, I.I.C. Chio, D.A. Tuveson, Transcriptional Regulation by Nrf2, Antioxid Redox Signal 29(17) (2018) 1727–1745. 10.1089/ars.2017.7342.

[11] C.F. Pulliam, M.A. Fath, S. Sho, S.T. Johnson, B.A. Wagner, M. Singhania, A.L. Kalen, K. Bayanbold, S.R. Solst, B.G. Allen, B.N. George, J.M. Caster, G.R. Buettner, D.P. Riley, J.L. Keene, R.A. Beardsley, D.R. Spitz, Pharmacological ascorbate combined with rucosopasem selectively radio-chemo-sensitizes NSCLC via generation of H(2)O(2), Redox Biol 80 (2025) 103505. 10.1016/j.redox.2025.103505.

[12] J.D. Schoenfeld, M.S. Alexander, T.J. Waldron, Z.A. Sibenaller, D.R. Spitz, G.R. Buettner, B.G. Allen, J.J. Cullen, Pharmacological Ascorbate as a Means of Sensitizing Cancer Cells to Radio-Chemotherapy While Protecting Normal Tissue, Semin Radiat Oncol 29(1) (2019) 25–32. 10.1016/j.semradonc.2018.10.006.

[13] J.D. Schoenfeld, Z.A. Sibenaller, K.A. Mapuskar, B.A. Wagner, K.L. Cramer-Morales, M. Furqan, S. Sandhu, T.L. Carlisle, M.C. Smith, T. Abu Hejleh, D.J. Berg, J. Zhang, J. Keech, K.R. Parekh, S. Bhatia, V. Monga, K.L. Bodeker, L. Ahmann, S. Vollstedt, H. Brown, E.P.S. Kauffman, M.E. Schall, R.J. Hohl, G.H. Clamon, J.D. Greenlee, M.A. Howard, M.K. Schultz, B.J. Smith, D.P. Riley, F.E. Domann, J.J. Cullen, G.R. Buettner, J.M. Buatti, D.R. Spitz, B.G. Allen, O(2)(⋅-) and H(2)O(2)-Mediated Disruption of Fe Metabolism Causes the Differential Susceptibility of NSCLC and GBM Cancer Cells to Pharmacological Ascorbate, Cancer Cell 32(2) (2017) 268. 10.1016/j.ccell.2017.07.008.

[14] G.R. Buettner, The pecking order of free radicals and antioxidants: lipid peroxidation, alpha-tocopherol, and ascorbate, Arch Biochem Biophys 300(2) (1993) 535–43. 10.1006/abbi.1993.1074.

[15] B.G. Allen, K.L. Bodeker, M.C. Smith, V. Monga, S. Sandhu, R. Hohl, T. Carlisle, H. Brown, N. Hollenbeck, S. Vollstedt, J.D. Greenlee, M.A. Howard, K.A. Mapuskar, S.N. Seyedin, J.M. Caster, K.A. Jones, J.J. Cullen, D. Berg, B.A. Wagner, G.R. Buettner, M.J. TenNapel, B.J. Smith, D.R. Spitz, J.M. Buatti, First-in-Human Phase I Clinical Trial of Pharmacologic Ascorbate Combined with Radiation and Temozolomide for Newly Diagnosed Glioblastoma, Clin Cancer Res 25(22) (2019) 6590–6597. 10.1158/1078-0432.CCR-19-0594.

[16] K.L. Bodeker, B.J. Smith, D.J. Berg, C. Chandrasekharan, S. Sharif, N. Fei, S. Vollstedt, H. Brown, M. Chandler, A. Lorack, S. McMichael, J. Wulfekuhle, B.A. Wagner, G.R. Buettner, B.G. Allen, J.M. Caster, B. Dion, M. Kamgar, J.M. Buatti, J.J. Cullen, A randomized trial of pharmacological ascorbate, gemcitabine, and nab-paclitaxel for metastatic pancreatic cancer, Redox Biol 77 (2024) 103375. 10.1016/j.redox.2024.103375.

[17] M. Furqan, T. Abu-Hejleh, L.M. Stephens, S.M. Hartwig, S.L. Mott, C.F. Pulliam, M. Petronek, J.B. Henrich, M.A. Fath, J.C. Houtman, S.M. Varga, K.L. Bodeker, A.D. Bossler, A.M. Bellizzi, J. Zhang, V. Monga, H. Mani, M. Ivanovic, B.J. Smith, M.M. Byrne, W. Zeitler, B.A. Wagner, G.R. Buettner, J.J. Cullen, J.M. Buatti, D.R. Spitz, B.G. Allen, Pharmacological ascorbate improves the response to platinum-based chemotherapy in advanced stage non-small cell lung cancer, Redox Biol 53 (2022) 102318. 10.1016/j.redox.2022.102318.

[18] V. Monga, B.J. Miller, M. Tanas, S. Boukhar, B. Allen, C. Anderson, L. Stephens, S. Hartwig, S. Varga, J. Houtman, L. Wang, W. Zhang, O. Jaber, J. Thomason, D. Kuehn, M. Rajput, C. Metz, K.D. Zamba, S. Mott, C. Abanonu, S. Bhatia, M. Milhem, Intratumoral talimogene laherparepvec injection with concurrent preoperative radiation in patients with locally advanced soft-tissue sarcoma of the trunk and extremities: phase IB/II trial, J Immunother Cancer 9(7) (2021). 10.1136/jitc-2021-003119.

[19] M.A. Fath, I.M. Ahmad, C.J. Smith, J. Spence, D.R. Spitz, Enhancement of carboplatin-mediated lung cancer cell killing by simultaneous disruption of glutathione and thioredoxin metabolism, Clin Cancer Res 17(19) (2011) 6206–17. 10.1158/1078-0432.CCR-11-0736.

[20] L. Li, M.A. Fath, P.M. Scarbrough, W.H. Watson, D.R. Spitz, Combined inhibition of glycolysis, the pentose cycle, and thioredoxin metabolism selectively increases cytotoxicity and oxidative stress in human breast and prostate cancer, Redox Biol 4 (2015) 127–35. 10.1016/j.redox.2014.12.001.

[21] F. Radenkovic, O. Holland, J.J. Vanderlelie, A.V. Perkins, Selective inhibition of endogenous antioxidants with Auranofin causes mitochondrial oxidative stress which can be countered by selenium supplementation, Biochem Pharmacol 146 (2017) 42–52. 10.1016/j.bcp.2017.09.009.

[22] T.P. Keeley, G.E. Mann, Defining Physiological Normoxia for Improved Translation of Cell Physiology to Animal Models and Humans, Physiol Rev 99(1) (2019) 161–234. 10.1152/physrev.00041.2017.

[23] K.C. Falls-Hubert, A.L. Butler, K. Gui, M. Anderson, M. Li, J.M. Stolwijk, S.N. Rodman, 3rd, S.R. Solst, A. Tomanek-Chalkley, C.C. Searby, V.C. Sheffield, V. Sandfort, H. Schmidt, M.L. McCormick, B.R. Wels, B.G. Allen, G.R. Buettner, M.K. Schultz, D.R. Spitz, Disulfiram causes selective hypoxic cancer cell toxicity and radio-chemo-sensitization via redox cycling of copper, Free Radic Biol Med 150 (2020) 1–11. 10.1016/j.freeradbiomed.2020.01.186.

[24] S.R. McKeown, Defining normoxia, physoxia and hypoxia in tumours—implications for treatment response, The British Journal of Radiology 87(1035) (2014). 10.1259/bjr.20130676.

[25] K.E. Brandt, K.C. Falls, J.D. Schoenfeld, S.N. Rodman, Z. Gu, F. Zhan, J.J. Cullen, B.A. Wagner, G.R. Buettner, B.G. Allen, D.J. Berg, D.R. Spitz, M.A. Fath, Augmentation of intracellular iron using iron sucrose enhances the toxicity of pharmacological ascorbate in colon cancer cells, Redox Biol 14 (2018) 82–87. 10.1016/j.redox.2017.08.017.

[26] M. Singhania, A. Zaher, C.F. Pulliam, K. Bayanbold, C.C. Searby, J.D. Schoenfeld, K.A. Mapuskar, M.A. Fath, B.G. Allen, D.R. Spitz, M.S. Petronek, Ǫuantitative MRI Evaluation of Ferritin Overexpression in Non-Small-Cell Lung Cancer, Int J Mol Sci 25(4) (2024). 10.3390/ijms25042398.

[27] G.R. Buettner, B.A. Jurkiewicz, Ascorbate free radical as a marker of oxidative stress: an EPR study, Free Radic Biol Med 14(1) (1993) 49–55. 10.1016/0891-5849(93)90508-r.

[28] J. Du, J.J. Cullen, G.R. Buettner, Ascorbic acid: chemistry, biology and the treatment of cancer, Biochim Biophys Acta 1826(2) (2012) 443–57. 10.1016/j.bbcan.2012.06.003.

[29] D. Erudaitius, J. Mantooth, A. Huang, J. Soliman, C.M. Doskey, G.R. Buettner, V.G.J. Rodgers, Calculated cell-specific intracellular hydrogen peroxide concentration: Relevance in cancer cell susceptibility during ascorbate therapy, Free Radic Biol Med 120 (2018) 356–367. 10.1016/j.freeradbiomed.2018.03.044.

[30] G.G. Kramarenko, S.G. Hummel, S.M. Martin, G.R. Buettner, Ascorbate reacts with singlet oxygen to produce hydrogen peroxide, Photochem Photobiol 82(6) (2006) 1634–7. 10.1562/2006-01-12-RN-774.

[31] K.A. Mapuskar, K.H. Flippo, J.D. Schoenfeld, D.P. Riley, S. Strack, T.A. Hejleh, M. Furqan, V. Monga, F.E. Domann, J.M. Buatti, P.C. Goswami, D.R. Spitz, B.G. Allen, Mitochondrial Superoxide Increases Age-Associated Susceptibility of Human Dermal Fibroblasts to Radiation and Chemotherapy, Cancer Res 77(18) (2017) 5054–5067. 10.1158/0008-5472.CAN-17-0106.

[32] B.G. Slane, N. Aykin-Burns, B.J. Smith, A.L. Kalen, P.C. Goswami, F.E. Domann, D.R. Spitz, Mutation of succinate dehydrogenase subunit C results in increased O2.-, oxidative stress, and genomic instability, Cancer Res 66(15) (2006) 7615–20. 10.1158/0008-5472.CAN-06-0833.

[33] D.T. Erudaitius, G.R. Buettner, V.G.J. Rodgers, The latency of peroxisomal catalase in terms of effectiveness factor for pancreatic and glioblastoma cancer cell lines in the presence of high concentrations of H(2)O(2): Implications for the use of pharmacological ascorbate in cancer therapy, Free Radic Biol Med 156 (2020) 20–25. 10.1016/j.freeradbiomed.2020.05.023.

[34] B.A. Wagner, C.B. Evig, K.J. Reszka, G.R. Buettner, C.P. Burns, Doxorubicin increases intracellular hydrogen peroxide in PC3 prostate cancer cells, Arch Biochem Biophys 440(2) (2005) 181–90. 10.1016/j.abb.2005.06.015.

[35] M. Anderson, R. Greenwald, Handbook of methods for oxygen radical research, Florida: CRC Press Inc (1985) 317-23.

[36] O.W. Griffith, Determination of glutathione and glutathione disulfide using glutathione reductase and 2-vinylpyridine, Anal Biochem 106(1) (1980) 207–12. 10.1016/0003-2697(80)90139-6.

[37] A. Holmgren, M. Fagerstedt, The in vivo distribution of oxidized and reduced thioredoxin in Escherichia coli, J Biol Chem 257(12) (1982) 6926–30.

38. R.A. Greenwald, CRC Handbook of Methods for Oxygen Radical Research, CRC Press 1985.

[39] R.A. Lawrence, R.F. Burk, Glutathione peroxidase activity in selenium-deficient rat liver, Biochem Biophys Res Commun 71(4) (1976) 952–8. 10.1016/0006-291x(76)90747-6.

[40] J.M. Stolwijk, K.C. Falls-Hubert, C.C. Searby, B.A. Wagner, G.R. Buettner, Simultaneous detection of the enzyme activities of GPx1 and GPx4 guide optimization of selenium in cell biological experiments, Redox Biol 32 (2020) 101518. 10.1016/j.redox.2020.101518.

[41] N.L. Bray, H. Pimentel, P. Melsted, L. Pachter, Near-optimal probabilistic RNA-seq quantification, Nat Biotechnol 34(5) (2016) 525–7. 10.1038/nbt.3519.

[42] H. Pimentel, N.L. Bray, S. Puente, P. Melsted, L. Pachter, Differential analysis of RNA-seq incorporating quantification uncertainty, Nat Methods 14(7) (2017) 687–690. 10.1038/nmeth.4324.

[43] J. Du, A.N. Pope, B.R. O’Leary, B.A. Wagner, P.C. Goswami, G.R. Buettner, J.J. Cullen, The role of mitochondria in pharmacological ascorbate-induced toxicity, Sci Rep 12(1) (2022) 22521. 10.1038/s41598-022-27185-9.

[44] B.A. Wagner, S. Venkataraman, G.R. Buettner, The rate of oxygen utilization by cells, Free Radic Biol Med 51(3) (2011) 700–12. 10.1016/j.freeradbiomed.2011.05.024.

[45] J.R. Cantor, M. Abu-Remaileh, N. Kanarek, E. Freinkman, X. Gao, A. Louissaint, Jr., C.A. Lewis, D.M. Sabatini, Physiologic Medium Rewires Cellular Metabolism and Reveals Uric Acid as an Endogenous Inhibitor of UMP Synthase, Cell 169(2) (2017) 258–272 e17. 10.1016/j.cell.2017.03.023.

[46] W.B. Dunn, D. Broadhurst, P. Begley, E. Zelena, S. Francis-McIntyre, N. Anderson, M. Brown, J.D. Knowles, A. Halsall, J.N. Haselden, A.W. Nicholls, I.D. Wilson, D.B. Kell, R. Goodacre, C. Human Serum Metabolome, Procedures for large-scale metabolic profiling of serum and plasma using gas chromatography and liquid chromatography coupled to mass spectrometry, Nat Protoc 6(7) (2011) 1060–83. 10.1038/nprot.2011.335.

[47] J.A. Petsche, S.C. Huang, D.J. Foster, C.C. Searby, V.C. Sheffield, D.R. Spitz, C.S. Carter, Protocol to evaluate markers of reactiveoxygen species and labile iron from a single sample across in vitro and in in vivo lung cancer models, STAR Protoc 7(2) (2026) 104596. 10.1016/j.xpro.2026.104596.

[48] S.G. Menon, E.H. Sarsour, D.R. Spitz, R. Higashikubo, M. Sturm, H. Zhang, P.C. Goswami, Redox regulation of the G1 to S phase transition in the mouse embryo fibroblast cell cycle, Cancer Res 63(9) (2003) 2109–17.

[49] S. Zheng, W. Wang, J. Aldahdooh, A. Malyutina, T. Shadbahr, Z. Tanoli, A. Pessia, J. Tang, SynergyFinder Plus: Toward Better Interpretation and Annotation of Drug Combination Screening Datasets, Genomics Proteomics Bioinformatics 20(3) (2022) 587–596. 10.1016/j.gpb.2022.01.004.

[50] Y. Adachi, R. Kimura, K. Hirade, H. Ebi, Escaping KRAS: Gaining Autonomy and Resistance to KRAS Inhibition in KRAS Mutant Cancers, Cancers (Basel) 13(20) (2021). 10.3390/cancers13205081.

[51] A. Galan-Cobo, P. Sitthideatphaiboon, X. Ǫu, A. Poteete, M.A. Pisegna, P. Tong, P.H. Chen, L.K. Boroughs, M.L.M. Rodriguez, W. Zhang, F. Parlati, J. Wang, V. Gandhi, F. Skoulidis, R.J. DeBerardinis, J.D. Minna, J.V. Heymach, LKB1 and KEAP1/NRF2 Pathways Cooperatively Promote Metabolic Reprogramming with Enhanced Glutamine Dependence in KRAS-Mutant Lung Adenocarcinoma, Cancer Res 79(13) (2019) 3251–3267. 10.1158/0008-5472.CAN-18-3527.

[52] F. Skoulidis, L.A. Byers, L. Diao, V.A. Papadimitrakopoulou, P. Tong, J. Izzo, C. Behrens, H. Kadara, E.R. Parra, J.R. Canales, J. Zhang, U. Giri, J. Gudikote, M.A. Cortez, C. Yang, Y. Fan, M. Peyton, L. Girard, K.R. Coombes, C. Toniatti, T.P. Heffernan, M. Choi, G.M. Frampton, V. Miller, J.N. Weinstein, R.S. Herbst, K.K. Wong, J. Zhang, P. Sharma, G.B. Mills, W.K. Hong, J.D. Minna, J.P. Allison, A. Futreal, J. Wang, Wistuba, II, J.V. Heymach, Co-occurring genomic alterations define major subsets of KRAS-mutant lung adenocarcinoma with distinct biology, immune profiles, and therapeutic vulnerabilities, Cancer Discov 5(8) (2015) 860–77. 10.1158/2159-8290.CD-14-1236.

[53] M. Ghandi, F.W. Huang, J. Jane-Valbuena, G.V. Kryukov, C.C. Lo, E.R. McDonald, 3rd, J. Barretina, E.T. Gelfand, C.M. Bielski, H. Li, K. Hu, A.Y. Andreev-Drakhlin, J. Kim, J.M. Hess, B.J. Haas, F. Aguet, B.A. Weir, M.V. Rothberg, B.R. Paolella, M.S. Lawrence, R. Akbani, Y. Lu, H.L. Tiv, P.C. Gokhale, A. de Weck, A.A. Mansour, C. Oh, J. Shih, K. Hadi, Y. Rosen, J. Bistline, K. Venkatesan, A. Reddy, D. Sonkin, M. Liu, J. Lehar, J.M. Korn, D.A. Porter, M.D. Jones, J. Golji, G. Caponigro, J.E. Taylor, C.M. Dunning, A.L. Creech, A.C. Warren, J.M. McFarland, M. Zamanighomi, A. Kauffmann, N. Stransky, M. Imielinski, Y.E. Maruvka, A.D. Cherniack, A. Tsherniak, F. Vazquez, J.D. Jaffe, A.A. Lane, D.M. Weinstock, C.M. Johannessen, M.P. Morrissey, F. Stegmeier, R. Schlegel, W.C. Hahn, G. Getz, G.B. Mills, J.S. Boehm, T.R. Golub, L.A. Garraway, W.R. Sellers, Next-generation characterization of the Cancer Cell Line Encyclopedia, Nature 569(7757) (2019) 503–508. 10.1038/s41586-019-1186-3.

[54] E.L. Greer, P.R. Oskoui, M.R. Banko, J.M. Maniar, M.P. Gygi, S.P. Gygi, A. Brunet, The energy sensor AMP-activated protein kinase directly regulates the mammalian FOXO3 transcription factor, J Biol Chem 282(41) (2007) 30107–19. 10.1074/jbc.M705325200.

[55] X. Hou, J. Song, X.N. Li, L. Zhang, X. Wang, L. Chen, Y.H. Shen, Metformin reduces intracellular reactive oxygen species levels by upregulating expression of the antioxidant thioredoxin via the AMPK-FOXO3 pathway, Biochem Biophys Res Commun 396(2) (2010) 199–205. 10.1016/j.bbrc.2010.04.017.

[56] X.N. Li, J. Song, L. Zhang, S.A. LeMaire, X. Hou, C. Zhang, J.S. Coselli, L. Chen, X.L. Wang, Y. Zhang, Y.H. Shen, Activation of the AMPK-FOXO3 pathway reduces fatty acid-induced increase in intracellular reactive oxygen species by upregulating thioredoxin, Diabetes 58(10) (2009) 2246–57. 10.2337/db08-1512.

[57] M.S. Padanad, G. Konstantinidou, N. Venkateswaran, M. Melegari, S. Rindhe, M. Mitsche, C. Yang, K. Batten, K.E. Huffman, J. Liu, X. Tang, J. Rodriguez-Canales, N. Kalhor, J.W. Shay, J.D. Minna, J. McDonald, Wistuba, II, R.J. DeBerardinis, P.P. Scaglioni, Fatty Acid Oxidation Mediated by Acyl-CoA Synthetase Long Chain 3 Is Required for Mutant KRAS Lung Tumorigenesis, Cell Rep 16(6) (2016) 1614–1628. 10.1016/j.celrep.2016.07.009.

[58] N. Aykin-Burns, I.M. Ahmad, Y. Zhu, L.W. Oberley, D.R. Spitz, Increased levels of superoxide and H2O2 mediate the differential susceptibility of cancer cells versus normal cells to glucose deprivation, Biochem J 418(1) (2009) 29–37. 10.1042/BJ20081258.

[59] I.B. Bize, L.W. Oberley, H.P. Morris, Superoxide dismutase and superoxide radical in Morris hepatomas, Cancer Res 40(10) (1980) 3686–93.

[60] L.W. Oberley, G.R. Buettner, Role of superoxide dismutase in cancer: a review, Cancer Res 39(4) (1979) 1141–9.

[61] L.W. Oberley, T.D. Oberley, G.R. Buettner, Cell differentiation, aging and cancer: the possible roles of superoxide and superoxide dismutases, Med Hypotheses 6(3) (1980) 249–68. 10.1016/0306-9877(80)90123-1.

[62] D.R. Spitz, J.E. Sim, L.A. Ridnour, S.S. Galoforo, Y.J. Lee, Glucose deprivation-induced oxidative stress in human tumor cells. A fundamental defect in metabolism?, Ann N Y Acad Sci 899 (2000) 349–62. 10.1111/j.1749-6632.2000.tb06199.x.

[63] E.L. Springer, Comparative study of the cytoplasmic organelles of epithelial cell lines derived from human carcinomas and nonmalignant tissues, Cancer Res 40(3) (1980) 803–17.

[64] M.J. Bollong, G. Lee, J.S. Coukos, H. Yun, C. Zambaldo, J.W. Chang, E.N. Chin, I. Ahmad, A.K. Chatterjee, L.L. Lairson, P.G. Schultz, R.E. Moellering, A metabolite-derived protein modification integrates glycolysis with KEAP1-NRF2 signalling, Nature 562(7728) (2018) 600–604. 10.1038/s41586-018-0622-0.

[65] Y. Mitsuishi, K. Taguchi, Y. Kawatani, T. Shibata, T. Nukiwa, H. Aburatani, M. Yamamoto, H. Motohashi, Nrf2 redirects glucose and glutamine into anabolic pathways in metabolic reprogramming, Cancer Cell 22(1) (2012) 66–79. 10.1016/j.ccr.2012.05.016.

[66] G. Cascio, K.N. Aguirre, K.P. Church, R.O. Hughes, L.A. Nease, I. Delclaux, H.J. Davis, E. Piskounova, Transcriptional Isoforms of NAD(+) kinase regulate oxidative stress resistance and melanoma metastasis, Redox Biol 76 (2024) 103289. 10.1016/j.redox.2024.103289.

[67] X. Yan, S. Wu, X. Fan, Ǫ. Li, Y. Teng, N. Wang, J. Zhang, NRF2 Deficiency Disrupts Mitochondrial Homeostasis via NDUFS7 in Trabecular Meshwork, Research (Wash D C) 9 (2026) 1203. 10.34133/research.1203.

[68] M.H. Ludtmann, P.R. Angelova, Y. Zhang, A.Y. Abramov, A.T. Dinkova-Kostova, Nrf2 affects the efficiency of mitochondrial fatty acid oxidation, Biochem J 457(3) (2014) 415–24. 10.1042/BJ20130863.

[69] B.J. Sishc, L. Ding, T.K. Nam, C.D. Heer, S.N. Rodman, J.D. Schoenfeld, M.A. Fath, D. Saha, C.F. Pulliam, B. Langen, R.A. Beardsley, D.P. Riley, J.L. Keene, D.R. Spitz, M.D. Story, Avasopasem manganese synergizes with hypofractionated radiation to ablate tumors through the generation of hydrogen peroxide, Sci Transl Med 13(593) (2021). 10.1126/scitranslmed.abb3768.

