## Supplementary material for "Resistance to Pharmacological Ascorbate, Thioredoxin Reductase Inhibition, and Radiation in KEAP1 Mutant Lung Cancer Cells is Mediated by NRF2": Figure S1-S17

| Mutational Status | A549 | H2030 | H460 | Calu-6 | H1299 | H292 |
|---|---|---|---|---|---|---|
| **KRAS Status** | NP_004976.2: p.Gly12Ser (G12S) | NP_004976.2: p.Gly12Cys (G12C) | NP_004976.2: p.Gln61His (Q61H) | NP_004976.2: p.Gln61Lys (Q61K) | WT | WT |
| **STK11 (LKB1) Status** | NP_000446.1: p.Gln37Ter (Q37*) | NP_000446.1: p.Glu317Ter (E317*) | NP_000446.1: p.Gln37Ter (Q37*) | WT | WT | WT |
| **KEAP1 Status** | NP_036421.2: p.Gly333Cys (G333C) | NP_036421.2: p.Val568Phe (V568F) | NP_036421.2: p.Asp236His (D236H) | WT | WT | WT |
| **NRF2 (NFE2L2) Status** | WT | WT | WT | WT | WT | WT |

**Figure S1.** KRAS, STK11 (LKB1), KEAP1, and NRF2 (NFE2L2) statuses of the cell lines under investigation. Adapted from DepMap [53].

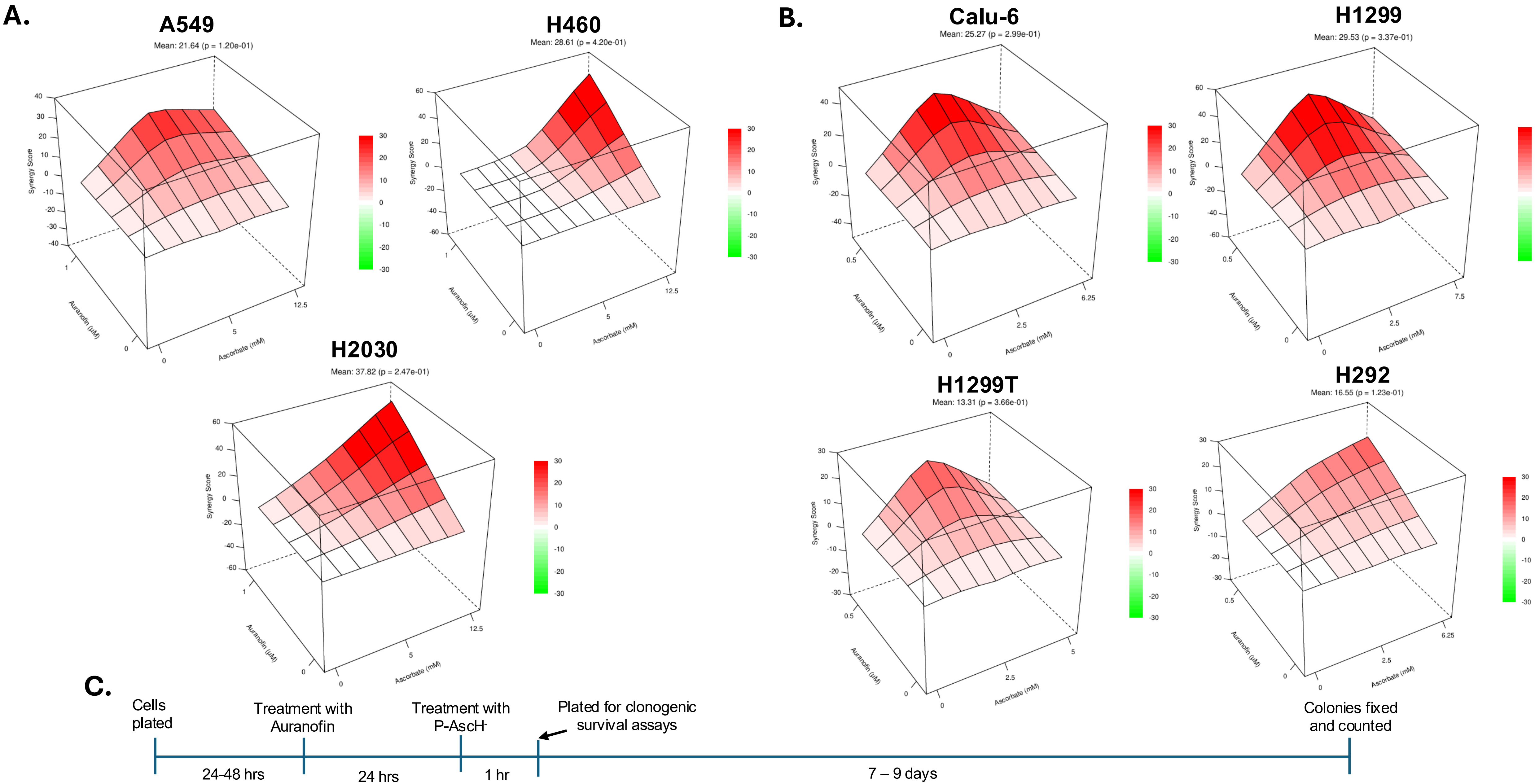


**Figure S2. Loewe synergy plots and scores represent the synergistic effect of Auranofin (0.5 -1 µM) plus P-AscH$^-$ (2.5 mM (5 pmol/cell) - 12.5 mM (40 pmol/cell)) in NSCLC cells.** (A) Loewe synergy plots showing the synergistic effect of Auranofin (1 µM , 24 hours) plus P-AscH$^-$ (5 mM (10 pmol/cell) and 12.5 mM (40 pmol/cell)) in A549, H460, and H2030 cells. (B) Loewe synergy plots showing the synergistic effect of Auranofin (0.5 µM, 24 hours) plus P-AscH$^-$ (2.5 mM (5 pmol/cell), 6.25 mM (15 pmol/cell), and 7.5 mM (20 pmol/cell), 1 hour) in Calu-6, H1299, H1299T, and H292 cells. Cells were maintained and treated in 5% $CO_2$, 21% $O_2$, and humidified in a 37°C incubator. Synergistic scores below -10 are considered antagonistic, the scores from -10 to 10 are considered additive, and the scores above 10 are considered synergistic interactions [49]. (C) Timeline of the treatments and the experimental procedures.

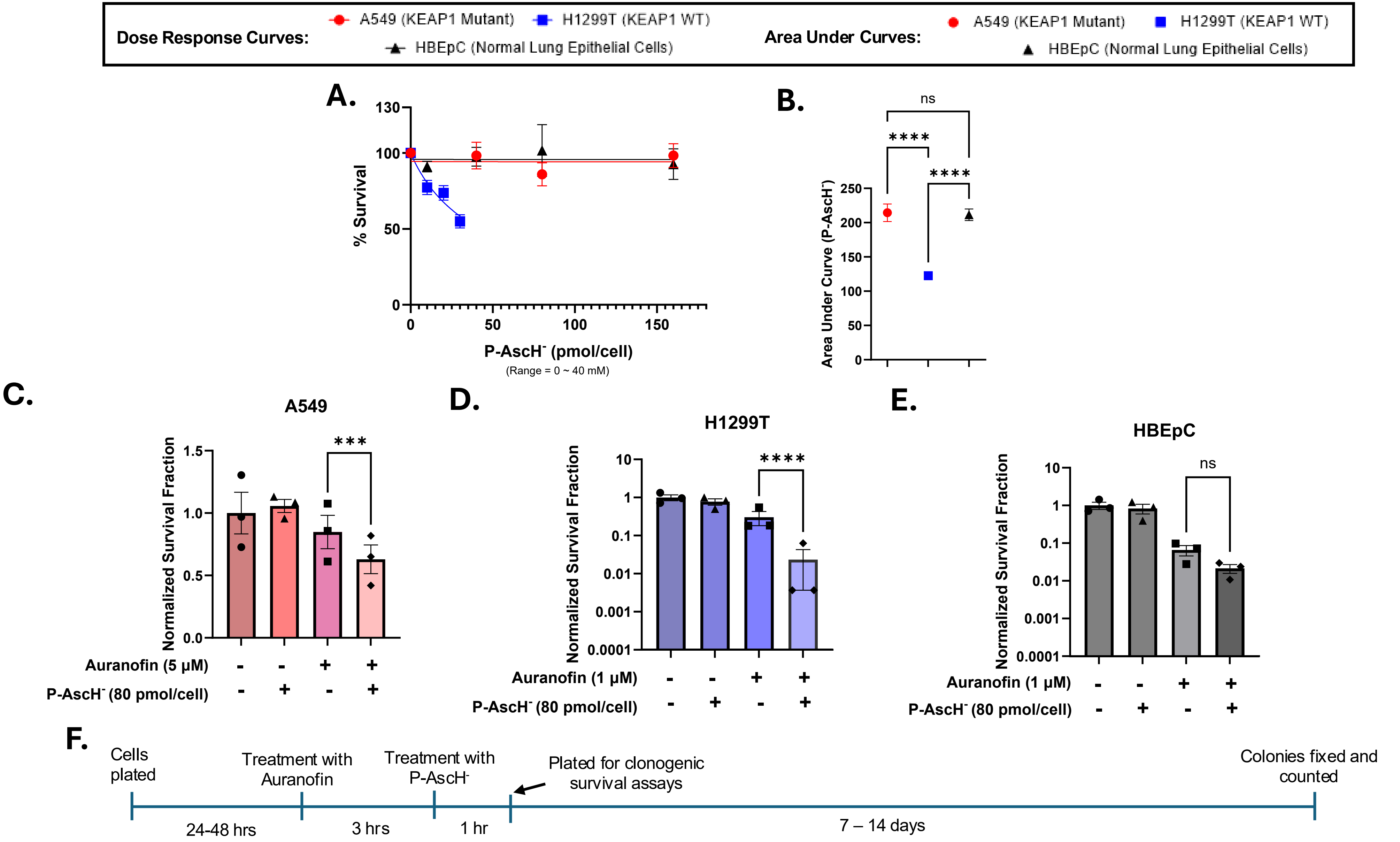


**Figure S3. P-AscH- is non-toxic to normal lung epithelial cells (HBEpC) and to A549 cells, and Auranofin does not sensitize HBEpC but NSCLC cells to P-AscH- in 4 % $O_2$.** (A) Dose response curves of A549, H1299T, and HBEpC cells toward 1-hour P-AscH- treatments at the indicated concentrations. (B) Quantification of area under curve from P-AscH- dose-response curves in NSCLC cells (A549 and H1299T) and normal lung epithelial cells (HBEpC). (C) Clonogenic Survival Assays of A549 cells after treatment with 5 µM Auranofin for 3 hours followed by treatments with P-AscH- (1 hour). (D) Clonogenic Survival Assays of H1299T cells after treatment with 1 µM Auranofin for 3 hours followed by treatments with P-AscH - (1 hour). (E) Clonogenic Survival Assays of HBEpC cells after treatment with 1 µM Auranofin for 3 hours followed by treatments with P-AscH- (1 hour). Cells were maintained and treated in 5% $CO_2$, 4% $O_2$, and humidified in a 37°C incubator. Concentrations of P-AscH- and were converted to $\log_{10}$ values before the calculation of area under curves of each cell line. Lower area under the curve values indicate higher sensitivity. Errors represent ± SEM, $n$ = 3. *p ≤ 0.05, **p ≤ 0.01, ***p ≤ 0.001, ****p ≤ 0.0001 by Welch's t test (area under curves) and Two-way ANOVA (clonogenic survival assays).

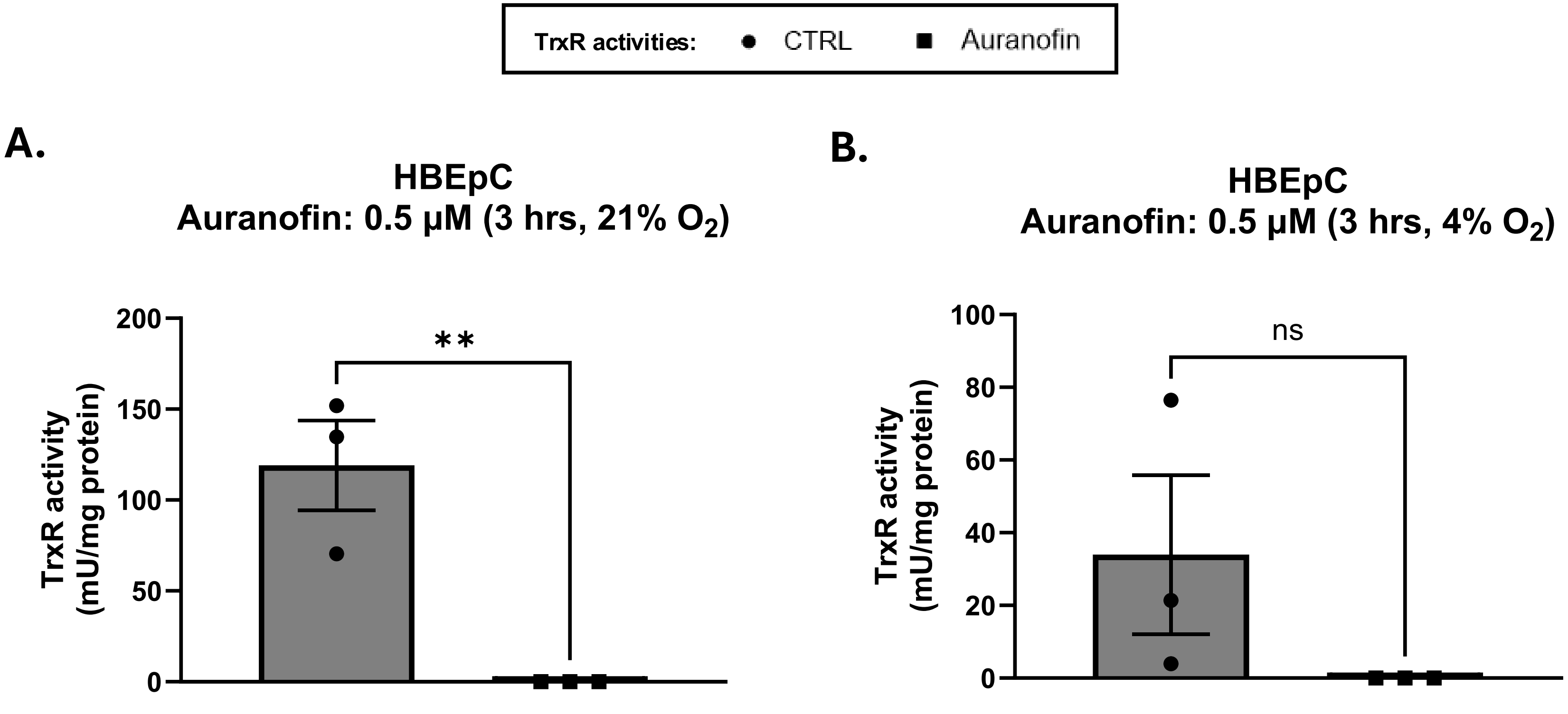


**Figure S4. 3-hour Auranofin treatment results in significant inhibition of TrxR activity in normal lung epithelial cells, and the TrxR activity changes in different $O_2$ levels. (**A and B) TrxR activities in HBEpC cells with or without Auranofin 0.5 µM (3-hour treatment). Cells were maintained and treated in 5% $CO_2$, humidified in a 37°C incubator, and 21 and 4 % $O_2$, respectively. Errors represent ± SEM, *n* = 3. *p $\leq 0.05$, **p $\leq 0.01$, ***p $\leq 0.001$, ****p $\leq 0.0001$ by Student's T-test.

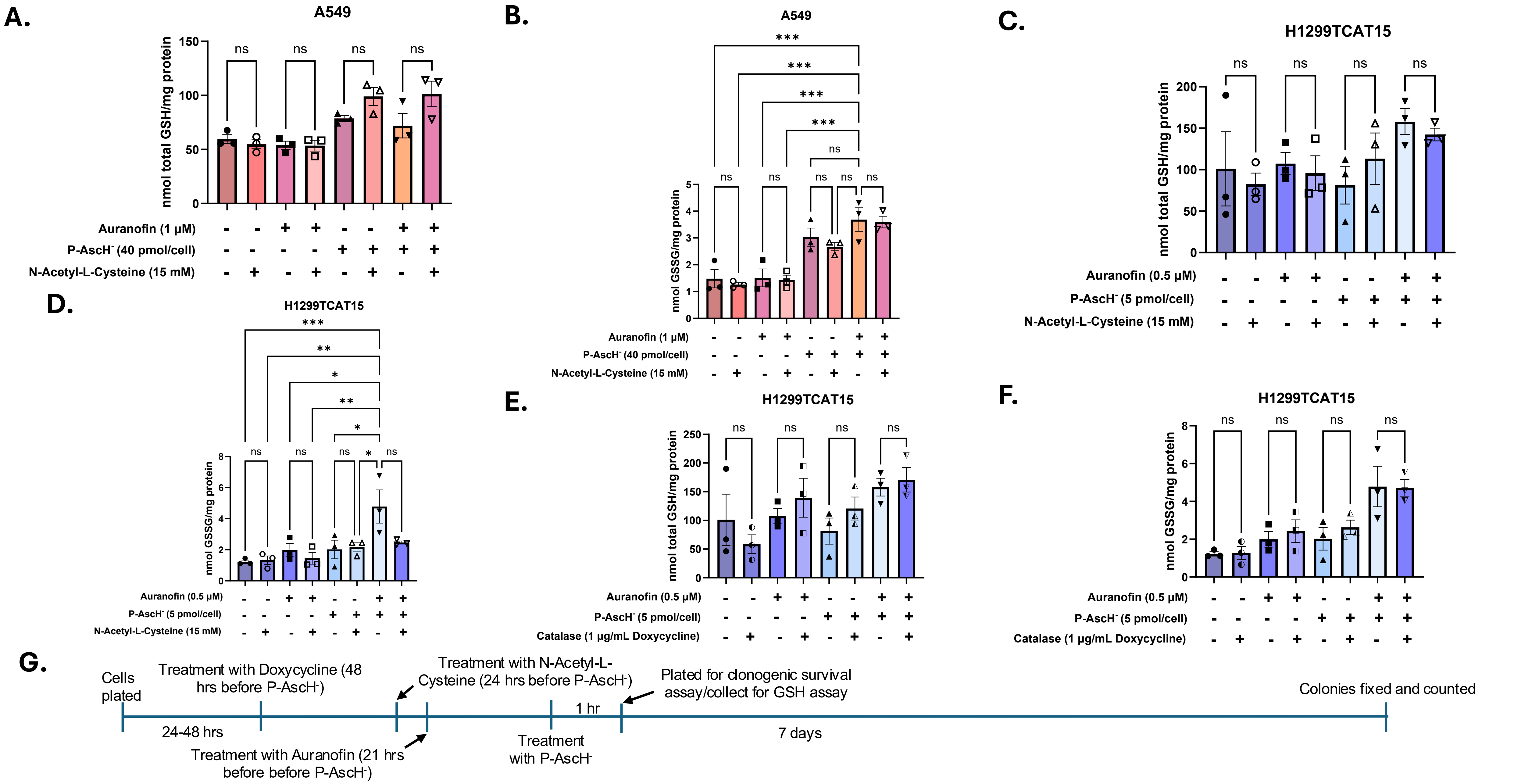


**Figure S5. N-Acetyl-L-Cysteine pretreatment does not cause change in total GSH levels but in % oxidized glutathione level in Auranofin + P-AscH$^-$ treatment group of A549 and H1299TCAT15 cells, and Catalase induction does not cause change in total or oxidized GSH levels.** (A and B): Total GSH levels and GSSG levels in A549 cells that were treated with NAC (15 mM, 24 hours), Auranofin (1 μM, 21 hours), and P-AscH$^-$ (40 pmol/cell, 1 hour). (C and D): Total GSH levels, and GSSG levels in H1299TCAT15 cells that were treated with NAC (15 mM, 24 hours), Auranofin (1 μM, 21 hours), and P-AscH$^-$ (40 pmol/cell, 1 hour). (E and F): Total GSH levels, and GSSG levels in H1299TCAT15 cells that were treated with doxycycline (1 μg/mL, 48 hours), Auranofin (0.5 μM, 21 hours), and P-AscH$^-$ (5 pmol/cell, 1 hour). (G) Timeline of the treatments and the experimental procedures. Errors represent ± SEM, *n* = 3. *$p \leq 0.05$, **$p \leq 0.01$, ***$p \leq 0.001$, ****$p \leq 0.0001$ by Two-way ANOVA.

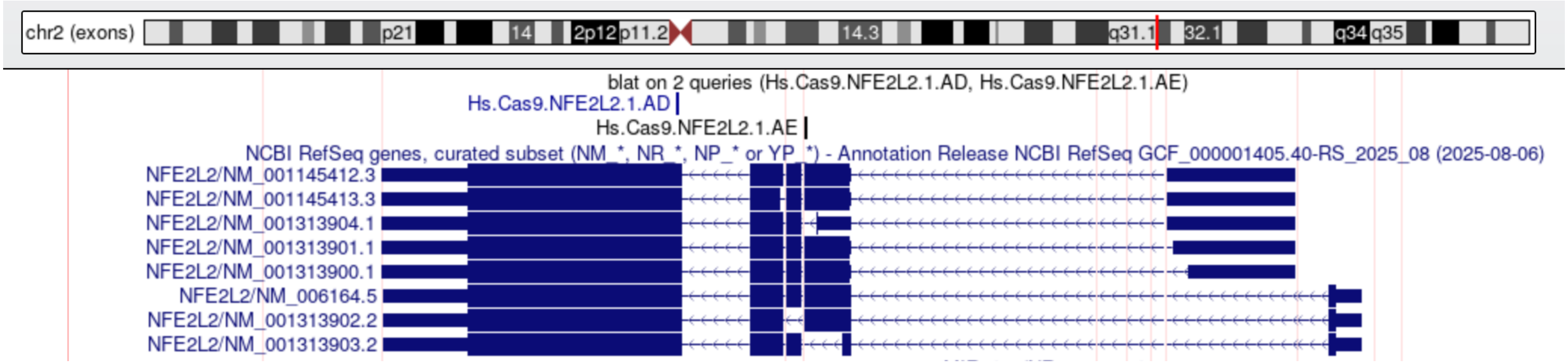


**Figure S6. NRF2 sgRNA 1.AD (Navy) targets all the identified transcript isoforms, while NRF2 sgRNA 1.AE (Black) does not target NM.001313903.2 and NM.001313904.1.** Genomic mapping of CRISPR-Cas9 guide RNAs (Hs.Cas9.NFE2L2.1.AD and 1.AE) targeting human NFE2L2 via the UCSC Genome Browser (hg38 assembly). Tracks display custom BLAT queries aligned against the NCBI RefSeq transcript subset. Introns are compressed using the exon view to highlight detail across all coding domains simultaneously. Hs.Cas9.NFE2L2.1.AD was used to create the A549 NRF2 Complete KO cell line, and Hs.Cas9.NFE2L2.1.AE was used to create the A549 Isoform Specific KO cell line.

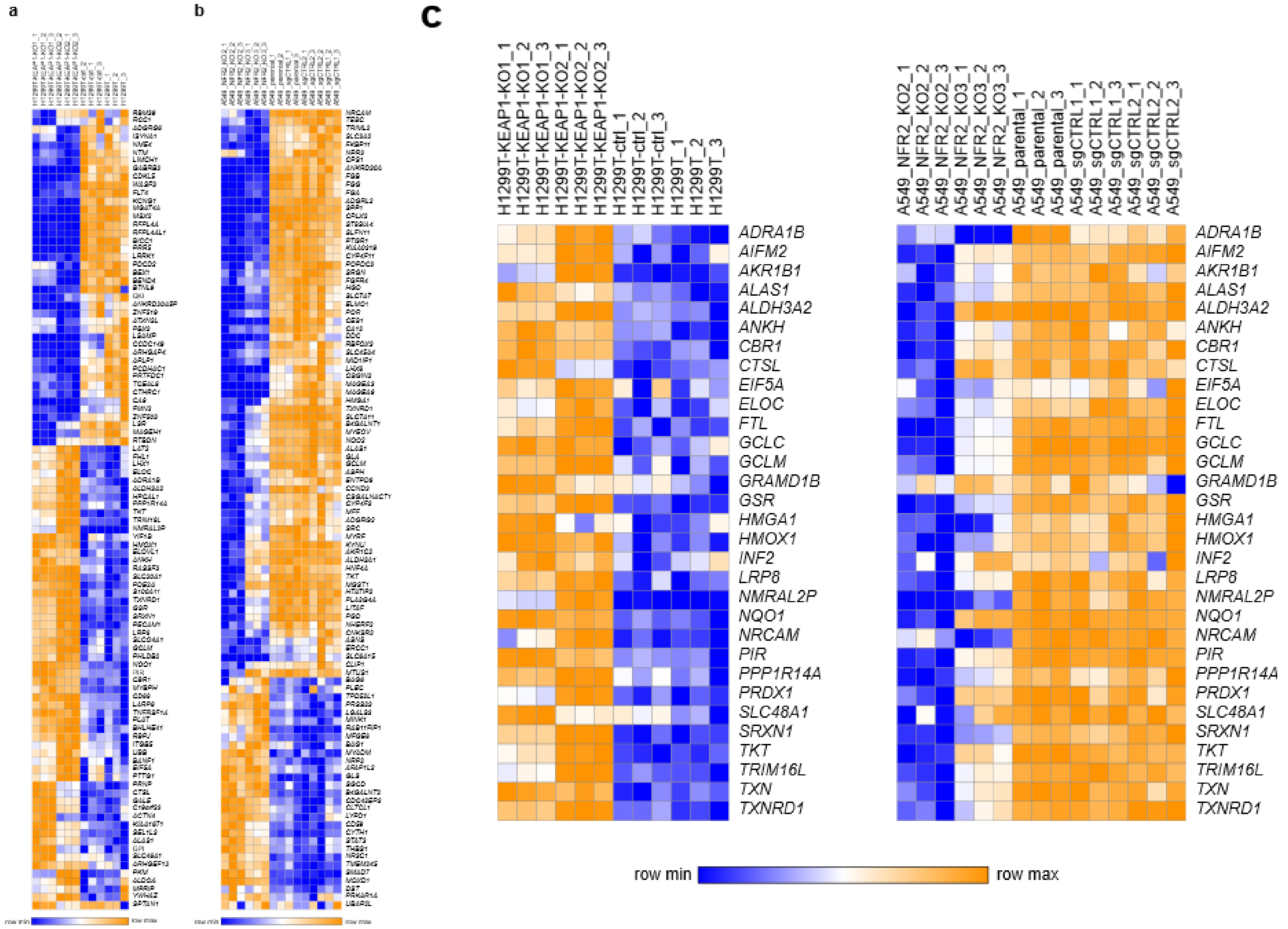


**Figure S7. Differential gene expression analysis of CRISPR edited HT1299T and A549 cells.** (**a**) Deletion of KEAP1 in H1299T cells significantly altered the expression of 192 genes (FDR ≤ 0.001, **Table S2**). The top 100 genes are shown here. (**b**) NRF2 deletion in A549 cells was associated with differential expression of 1736 genes (FDR ≤ 0.001, **Table S3**). Only the top 100 differentially expressed genes are shown. (**c**) Comparison of these two gene sets identified a signature of 31 genes that showed increased expression with KEAP1 deletion and decreased expression with NRF2 loss. As expected, this gene signature is enriched with genes involved with cellular redox homeostasis (GO:0045454, FDR = 6.12e-07), oxidoreductase activity (GO:0016491, FDR = 4.12e-07).

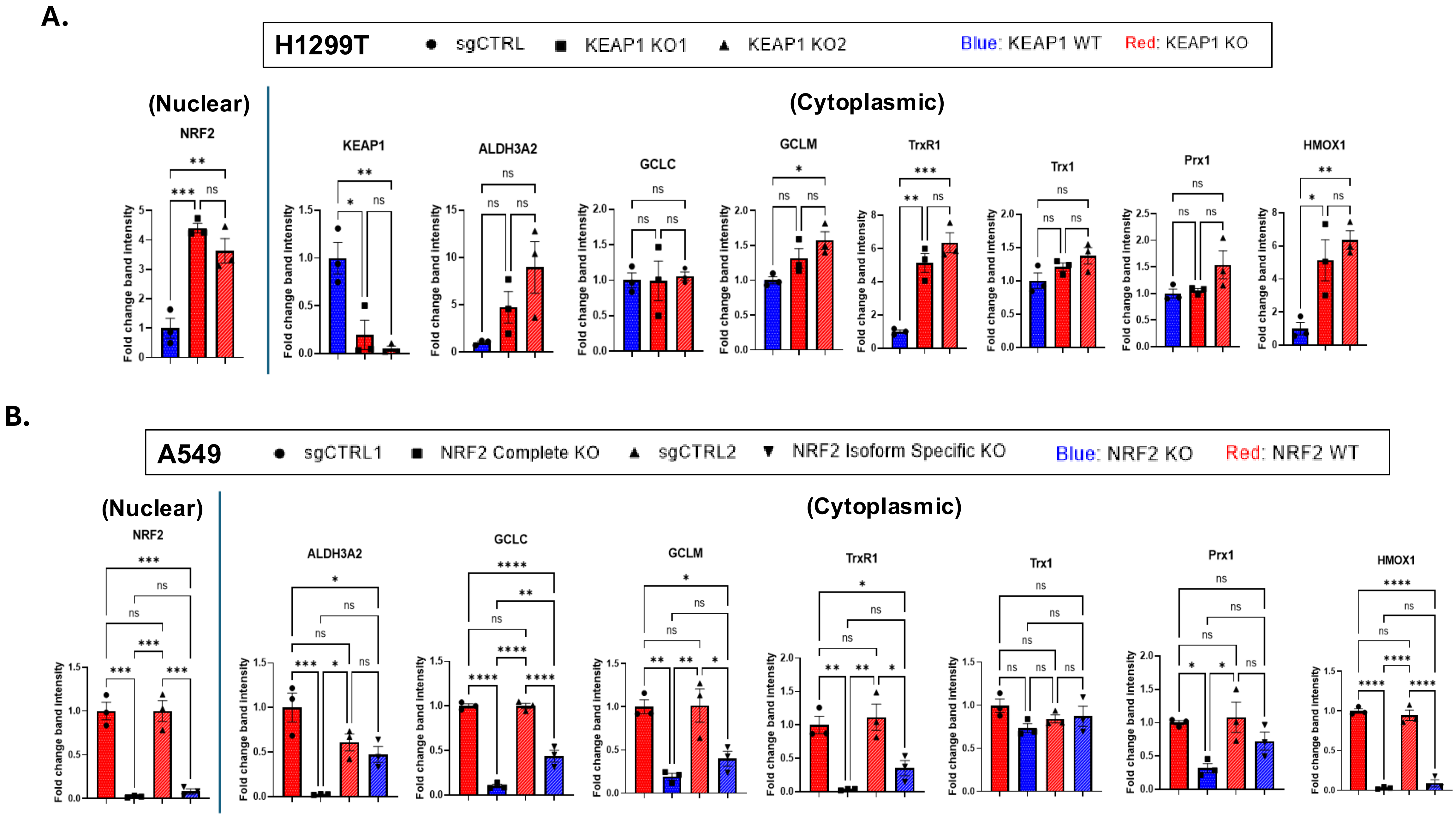


**Figure S8. KO of KEAP1 or NRF2 in NSCLC cells results changes in expressions of the NRF2 target proteins.** (A and B) Quantification of the Western Blot Images of each NRF2 target proteins (**Fig. 4A and 7A**). Errors represent ± SEM, *n* = 3. *p ≤ 0.05, **p ≤ 0.01, ***p ≤ 0.001, ****p ≤ 0.0001 by One-way ANOVA.

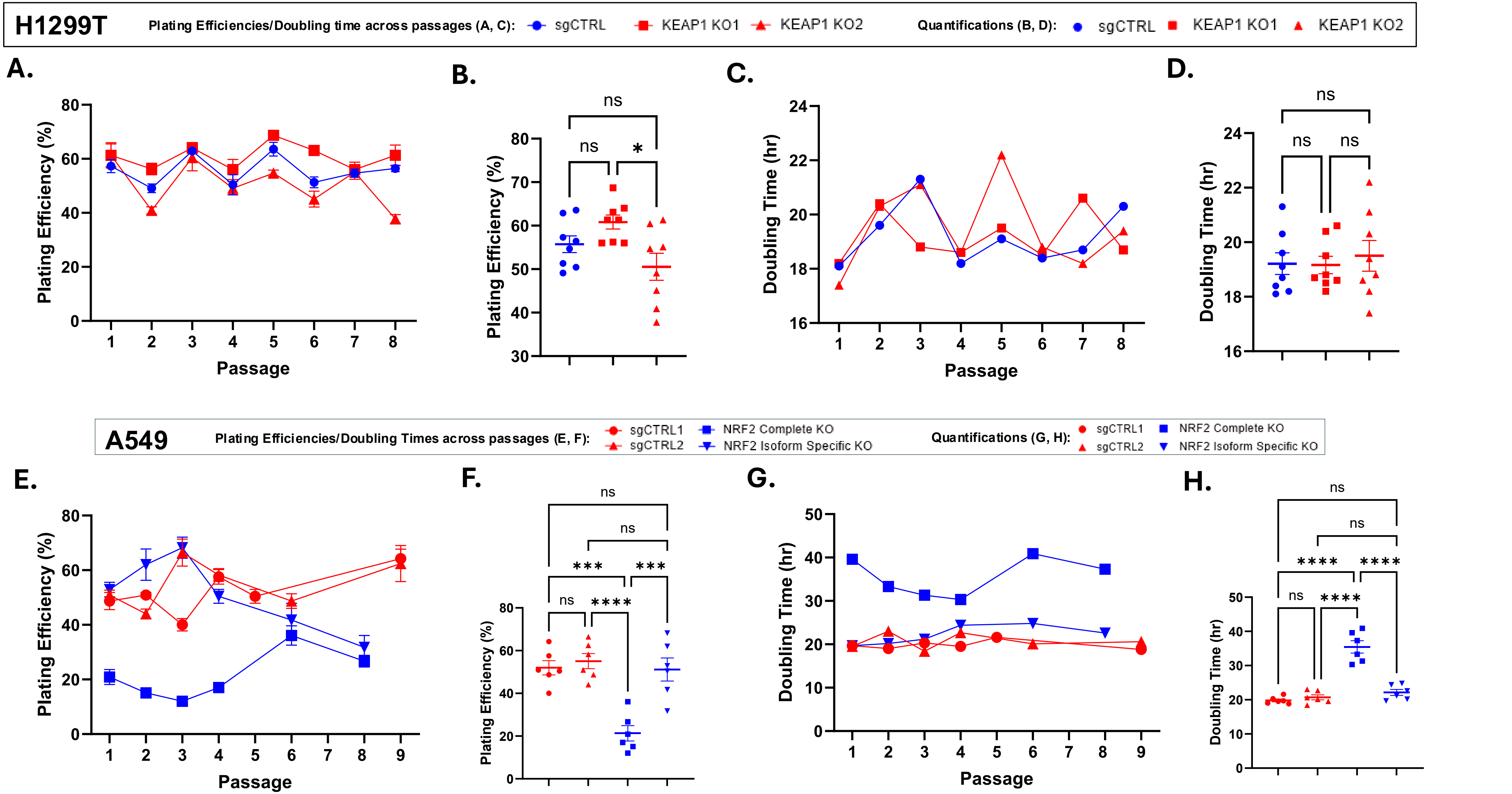


**Figure S9. KO of KEAP1 in H1299T does not cause change in plating efficiency and doubling time, but complete KO of NRF2 in A549 results in change in both.** (A and B) The changes and comparisons of plating efficiencies over the 8 passages in H1299T sgCTRL, and H1299T KEAP1 KO cells. (C and D) The changes and the comparisons in doubling time over the 8 passages in H1299T sgCTRL, and H1299T KEAP1 KO cells. (E and F) The changes and comparisons of plating efficiencies over the 8 passages in A549 sgCTRL, and A549 NRF2 KO cells. (G and H) The changes and the comparisons in doubling time over 8 – 9 passages in A549 sgCTRL, and A549 NRF2 KO cells. Cells were maintained in 5% $CO_2$, 21% $O_2$, and humidified in a 37°C incubator. Errors represent ± SEM, *n* = 3. *$p \leq 0.05$, **$p \leq 0.01$, ***$p \leq 0.001$, ****$p \leq 0.0001$ by One-Way ANOVA.

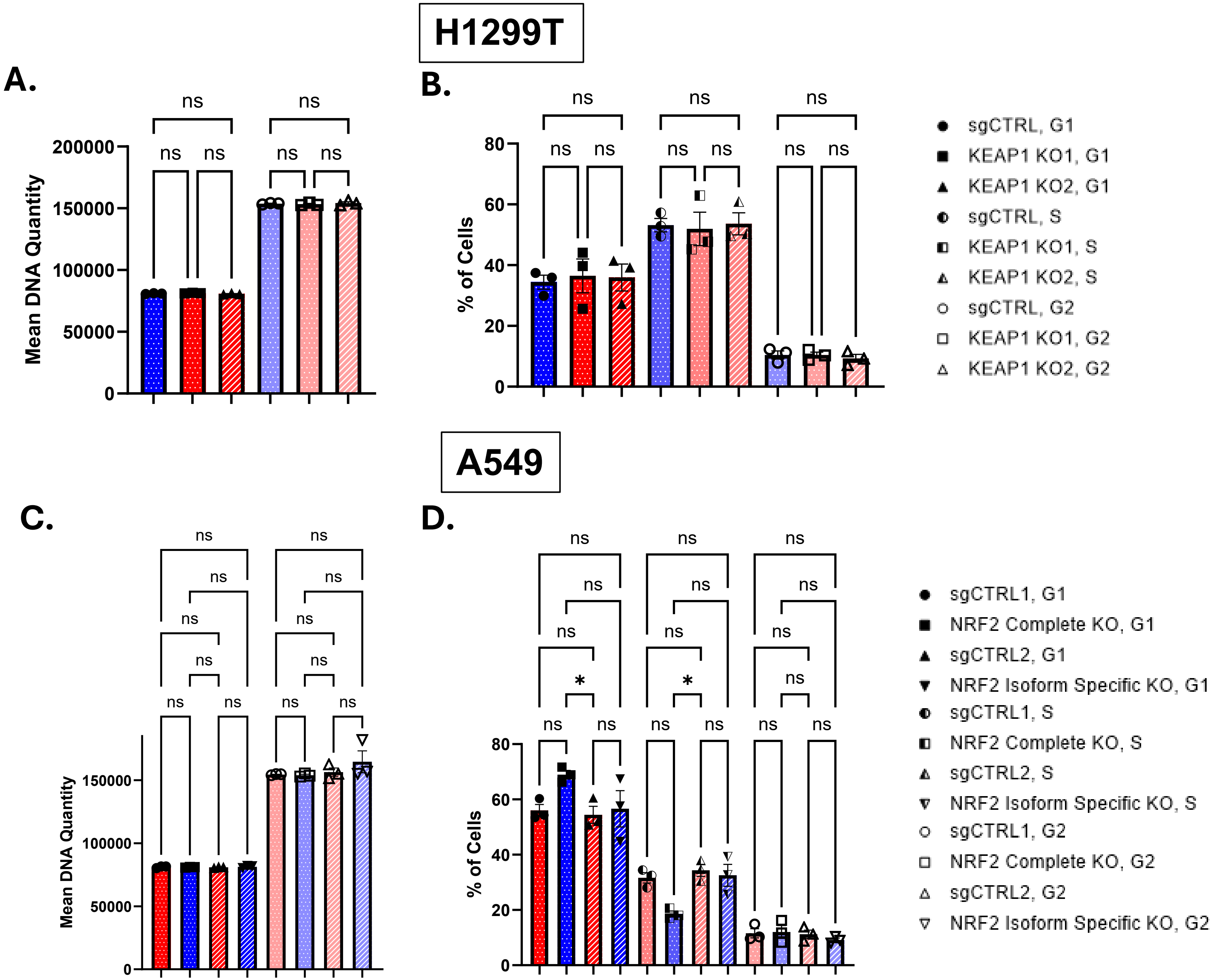


**Figure S10. KO of NRF2 in A549 results in higher percentage of cells in G1 cell cycle phase and lower percentage of cells in S phase, but KO of KEAP1 in H1299T results in no changes in the percentages.** (A and C) Mean DNA quantity of H1299TsgCTRL, H1299T KEAP1 KO, A549 sgCTRLs, and NRF2 KO cells in G1 and G2 phases. (B and D) percentage of H1299TsgCTRL, H1299T KEAP1 KO, A549 sgCTRLs, and NRF2 KO cells in G1, S, and G2 phases. Cells were maintained and treated in 5% $CO_2$, 21% $O_2$, and humidified in a 37°C incubator prior to staining with propidium iodide and Flow Cytometry. Errors represent ± SEM, *n* = 3. *$p \leq 0.05$, **$p \leq 0.01$, ***$p \leq 0.001$, ****$p \leq 0.0001$ by One-way ANOVA.

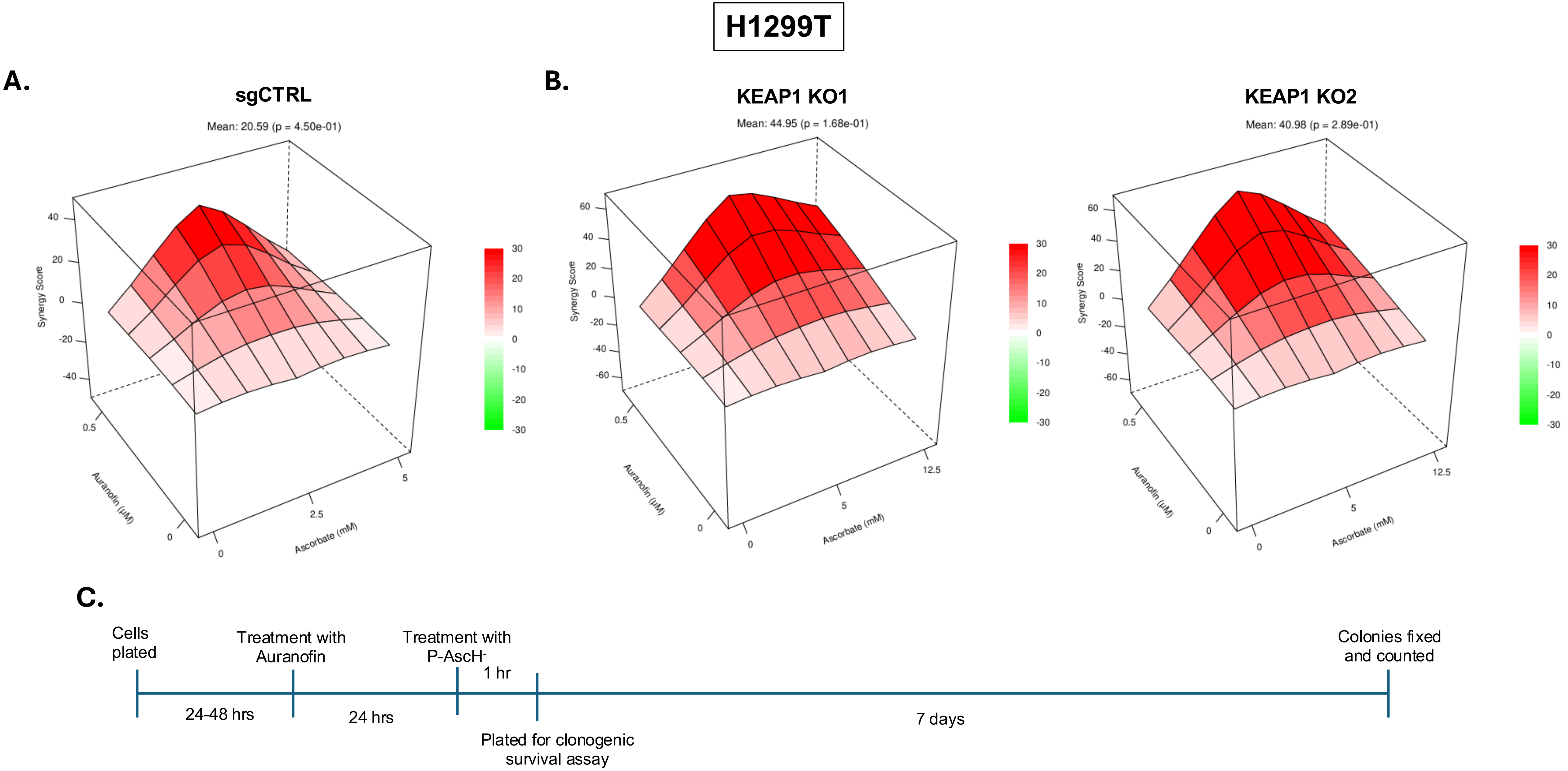


**Figure S11. Loewe synergy plots and scores represent the synergistic effect of Auranofin (0.5 µM) plus P-AscH$^-$ (2.5 mM (5 pmol/cell) - 12.5 mM (40 pmol/cell)) in H1299T sgCTRL and H129T KEAP1 KO cells.** (A) Loewe synergy scores representing the synergistic effect of Auranofin (0.5 µM, 24 hours) plus P-AscH$^-$ (2.5 mM (5 pmol/cell) and 5 mM (10 pmol/cell), 1 hour) toward H1299T sgCTRL cells. (B) Loewe synergy scores representing the synergistic effect of Auranofin (0.5 µM, 24 hours) plus P-AscH$^-$ (5 mM (10 pmol/cell) and 12.5 mM (40 pmol/cell), 1 hour) toward H1299T KEAP1 KO cells. (C)Timeline of the treatments and the experimental procedures. Cells were maintained and treated in 5% $CO_2$, 21% $O_2$, and humidified in a 37°C incubator. Synergistic scores below -10 are considered antagonistic, the scores from -10 to 10 are considered additive, and the scores above 10 are considered synergistic interactions [49].

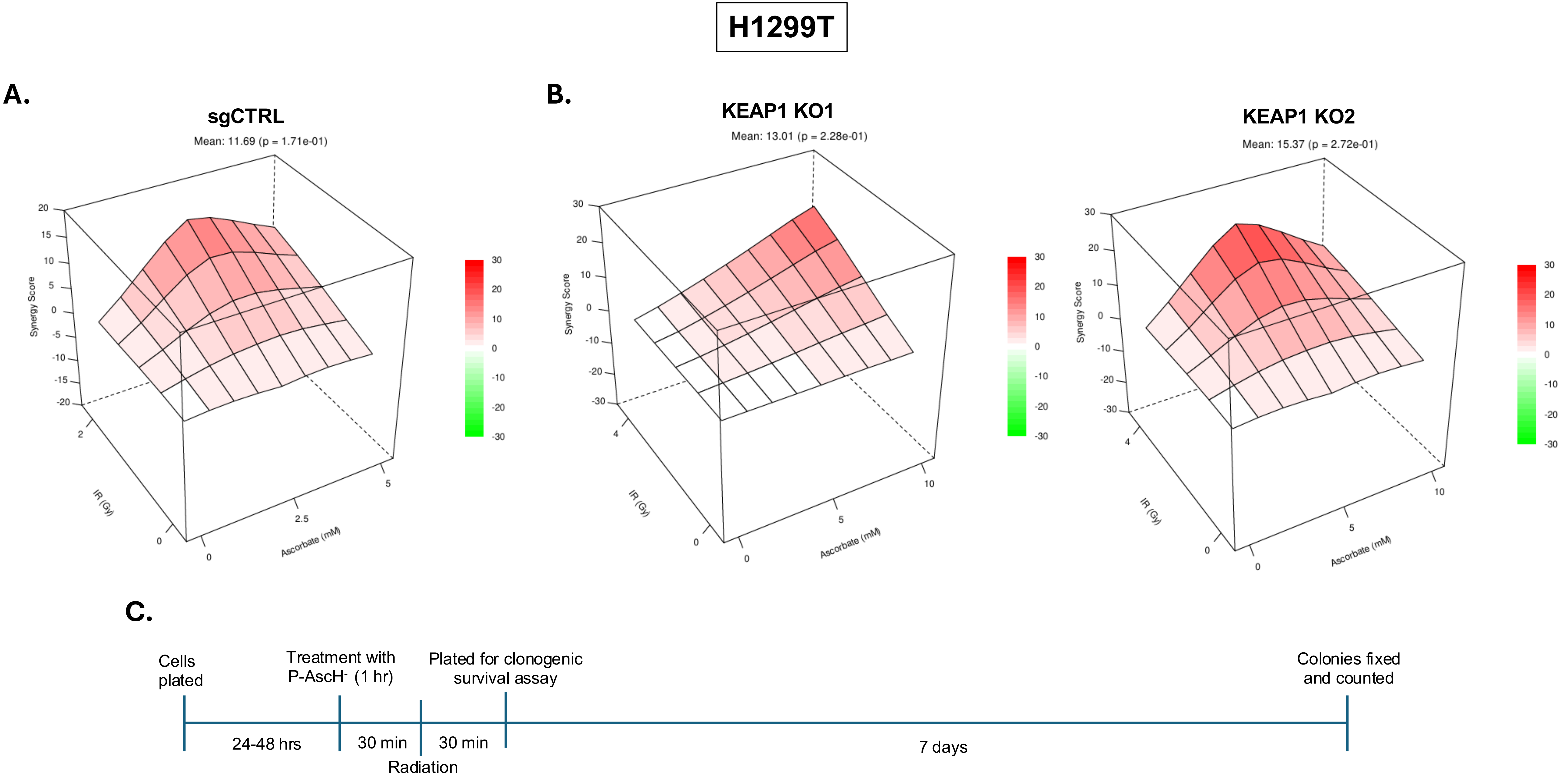


**Figure S12. Loewe synergy plots and scores represent the synergistic effect of P-AscH$^-$ (2.5 mM (5 pmol/cell) - 10 mM (30 pmol/cell)) plus radiation (2 or 4 Gy) in H1299T sgCTRL and H1299T KEAP1 KO cells.** (A) Loewe synergy scores representing the synergistic effect of P-AscH$^-$ (2.5 mM (5 pmol/cell) and 5 mM (10 pmol/cell), 1 hour) and radiation (IR, 2 Gy) in H1299T sgCTRL cells. (B) Loewe synergy scores representing the synergistic effect of P-AscH$^-$ (5 mM (10 pmol/cell) and 10 mM (30 pmol/cell), 1 hour) and radiation (IR, 4 Gy) in H1299T sgCTRL cells. (C) Timeline of the treatments and the experimental procedures. Cells were maintained and treated in 5% $CO_2$, 21% $O_2$, and humidified in a 37°C incubator. Synergistic scores below -10 are considered antagonistic, the scores from -10 to 10 are considered additive, and the scores above 10 are considered synergistic interactions [49].

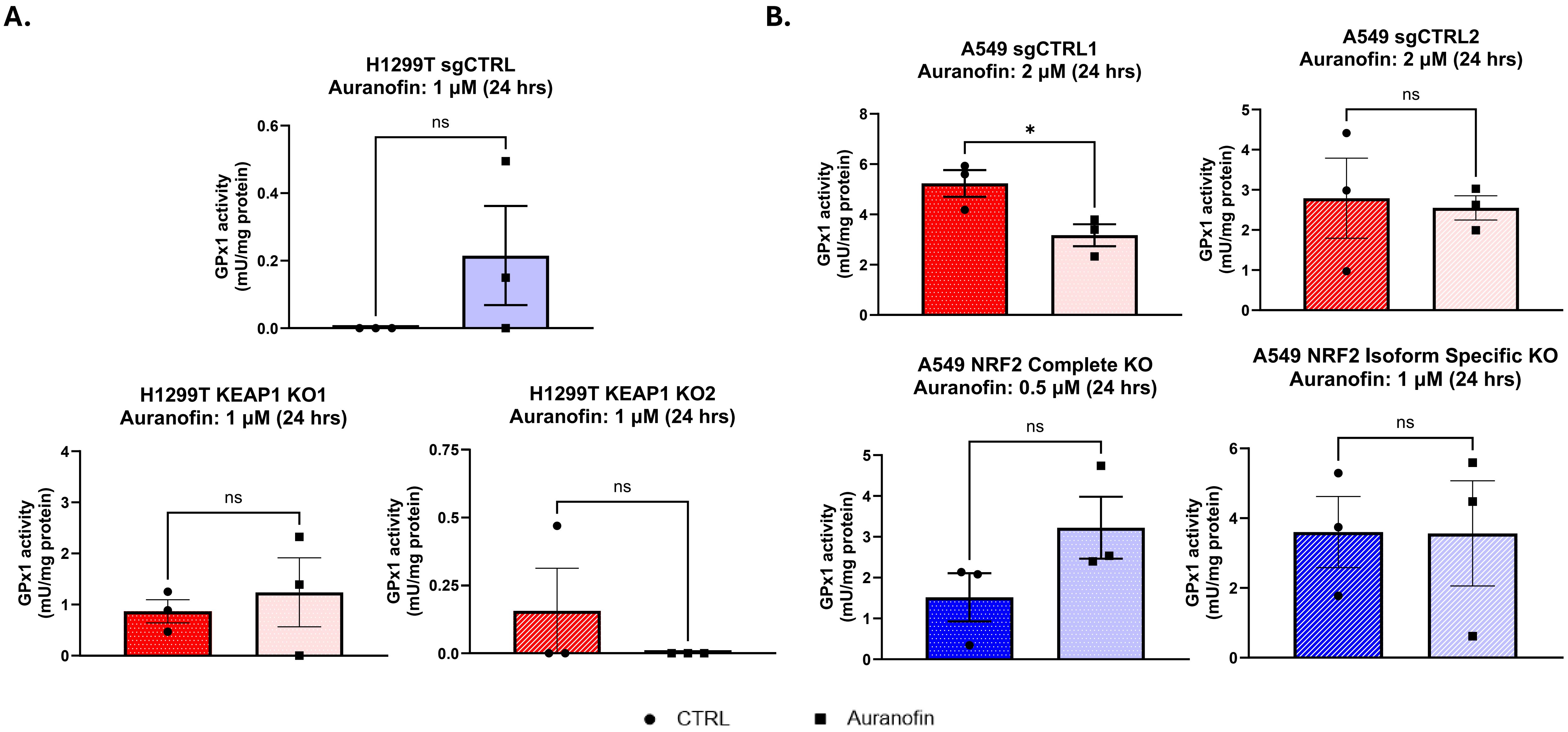


**Figure S13. Auranofin treatments does not result in inhibition of GPx1 activities in KEAP1 or NRF2 KO NSCLC cells. (**A) GPx1 activities in H1299T sgCTRL and KEAP1 KO cells with or without Auranofin 1 μM (24-hour treatment). (B) GPx1 activities in A549 sgCTRL and NRF2 KO cells with or without Auranofin (0 - 2 μM, 24-hour treatment). Cells were maintained and treated in 5% $CO_2$, 21% $O_2$, and humidified in a 37°C incubator. Errors represent ± SEM, *n* = 3. *$p \leq 0.05$, **$p \leq 0.01$, ***$p \leq 0.001$, ****$p \leq 0.0001$ by Student's T-test.

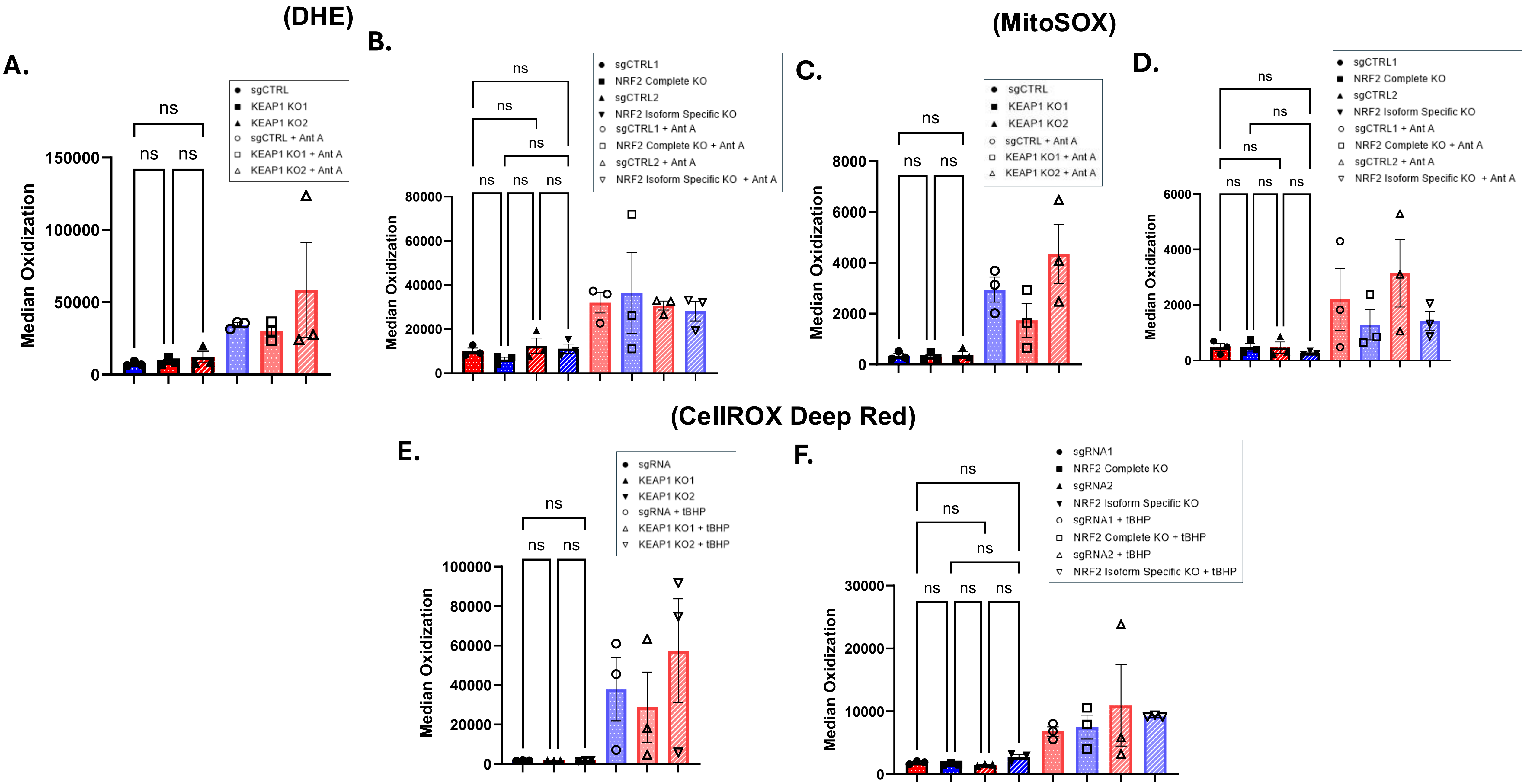


**Figure S14. KO of KEAP1 and NRF2 do not induce change in intracellular ROS levels, but NRF2 KO cells showed increased median oxidation when treated with tert-butyl hydroperoxide.** (A and B) DHE labeling of H1299T sgCTRL, H1299T KEAP1 KO, A549 sgCTRL, and A549 NRF2 KO cells. (C and D) MitoSOX labeling of H1299T sgCTRL, H1299T KEAP1 KO, A549 sgCTRL, and A549 NRF2 KO cells. (E and F) CellROX Deep Red labeling of H1299T sgCTRL, H1299T KEAP1 KO, A549 sgCTRL, and A549 NRF2 KO cells. Cells were maintained and treated in 5% $CO_2$, 21% $O_2$, and humidified in a 37°C incubator prior to staining and Flow Cytometry. Errors represent ± SEM, $n$ = 3. $^{*}p \leq 0.05$, $^{**}p \leq 0.01$, $^{***}p \leq 0.001$, $^{****}p \leq 0.0001$ by One-way ANOVA.

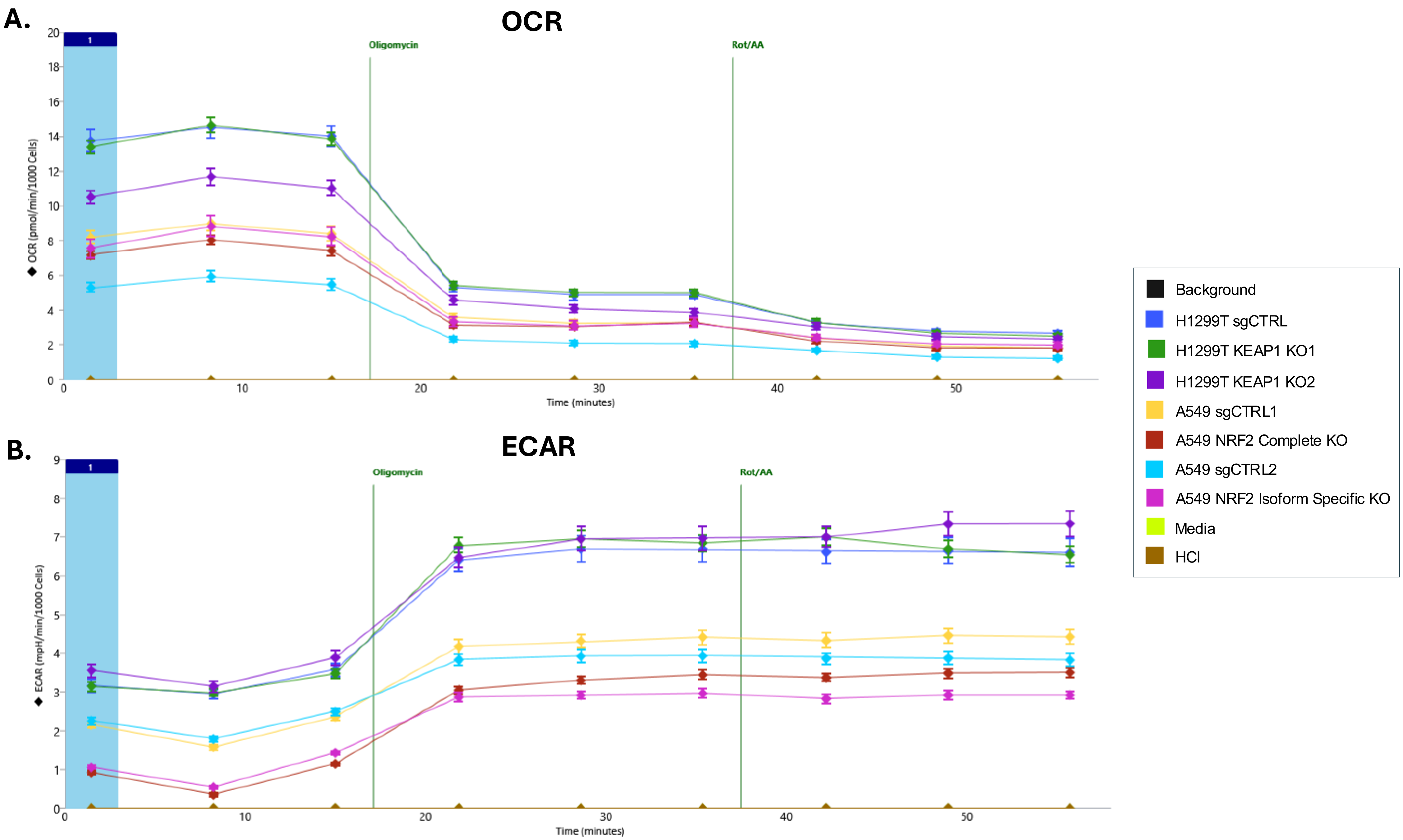


**Figure S15. KO of NRF2 results in decreased baseline ECAR but no change in baseline OCR, and KEAP1 KO2 show opposite effect in NSCLC cells.** Transitions of OCR (A) and ECAR (B) in H1299T sgCTRL, H1299T KEAP1 KO, A549 sgCTRL and NRF2 KO cells over time with Oligomycin, Rotenone, and Antimycin A treatments.

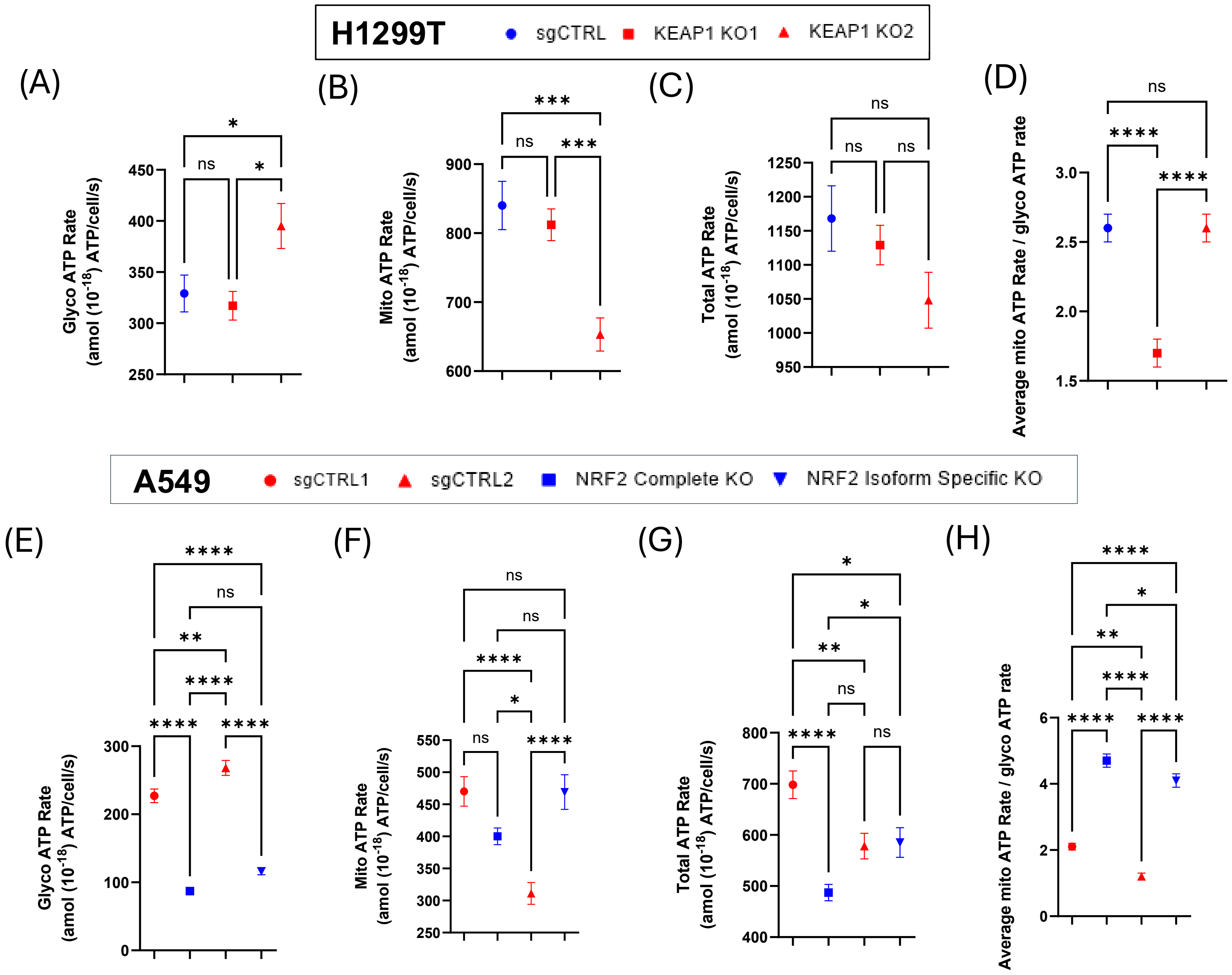


**Figure S16. KO of KEAP1 results in no change in total ATP rate, but complete KO of NRF2 results in decreased total ATP rate, and KEAP1 KO and NRF2 KO induce opposite effects on glycolytic ATP and OXPHOS ATP rates.** (A - D): Glycolytic ATP rates, mitochondrial ATP rates, total ATP rates, and the ratio of average mitochondrial and glycolytic ATP rates in H1299T sgCTRL and H1299T KEAP1 KO cells. (E-H) Glycolytic ATP rates, mitochondrial ATP rates, total ATP rates, and the ratio of average mitochondrial and glycolytic ATP rates in A549 sgCTRL and A549 NRF2 KO cells. Errors represent ± SEM, *n* = 3. *p ≤ 0.05, **p ≤ 0.01, ***p ≤ 0.001, ****p ≤ 0.0001 by One-way ANOVA.

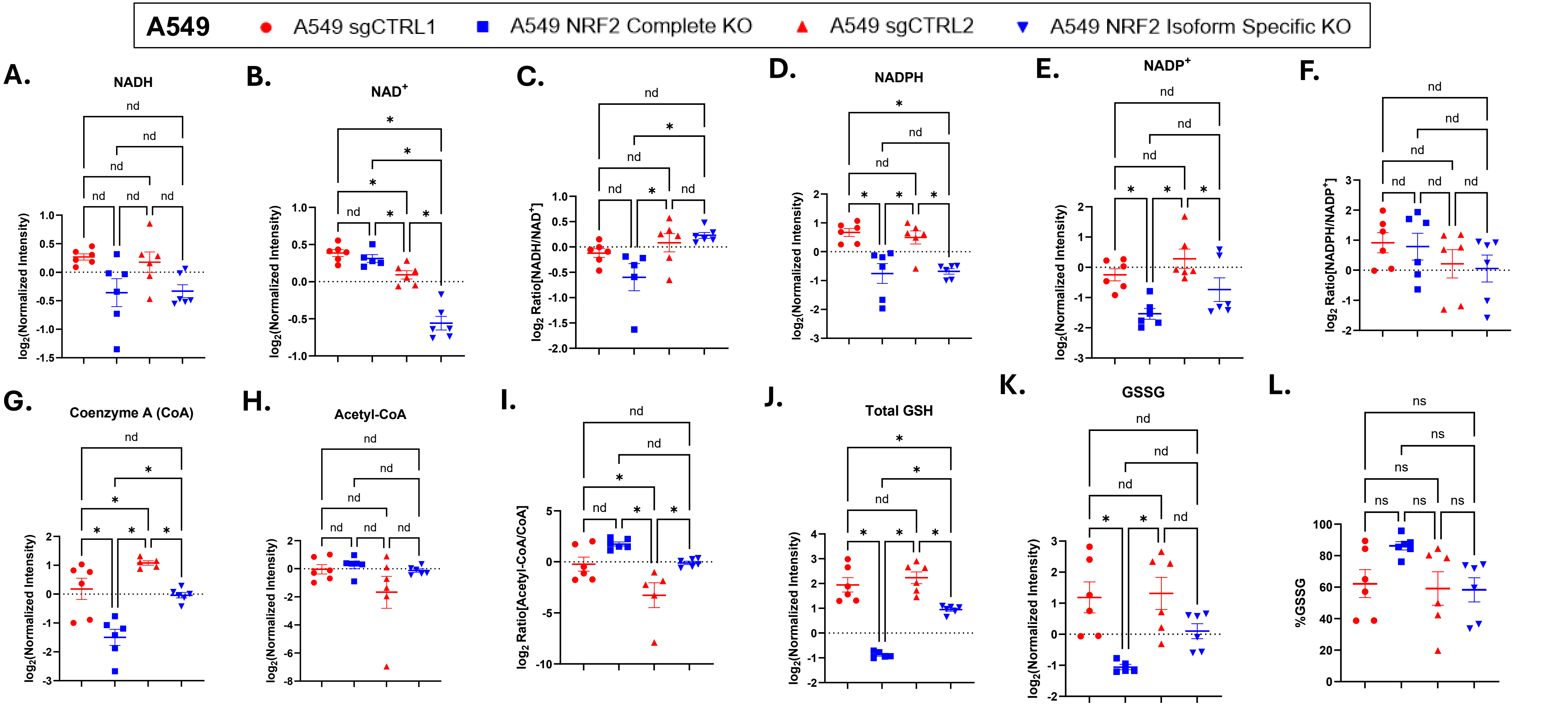


**Figure S17. KO of NRF2 in A549 results in decrease in NADPH, NADP⁺, Coenzyme A, total GSH land GSSG levels, and increase in Acetyl-CoA/Coenzyme A ratio, but not in NADH and NAD⁺ levels, and NADH/NAD⁺ and NADPH/NADP⁺ ratios.** (A) The $log_2$-transformed normalized NADH levels in A549 sgCTRL and A549 NRF2 KO cells. (B) The $log_2$-transformed normalized $NAD^+$ levels in A549 sgCTRL and A549 NRF2 KO cells. (C) The $log_2$-transformed $NADH/NAD^+$ ratios in A549 sgCTRL and A549 NRF2 KO cells. (D) The $log_2$-transformed normalized NADPH levels in A549 sgCTRL and A549 NRF2 KO cells. (E) The $log_2$-transformed normalized $NADP^+$ levels in A549 sgCTRL and A549 NRF2 KO cells. (F) The $log_2$-transformed $NADPH/NADP^+$ ratios in A549 sgCTRL and A549 NRF2 KO cells. (G) The $log_2$-transformed normalized Coenzyme A (CoA) levels in A549 sgCTRL and A549 NRF2 KO cells. (H) The $log_2$-transformed normalized Acetyl-CoA levels in A549 sgCTRL and A549 NRF2 KO cells. (I) The $log_2$-transformed Acetyl-CoA/CoA in A549 sgCTRL and A549 NRF2 KO cells. (J) The $log_2$-transformed total GSH levels in A549 sgCTRL and A549 NRF2 KO cells. (K) The $log_2$-transformed GSSG levels in A549 sgCTRL and A549 NRF2 KO cells. (L) % GSSG in A549 sgCTRL and A549 NRF2 KO cells. Cell samples were submitted to Metabolomics Core Facility, Carver College of Medicine, Iowa City, IA 52242 for LC-MS profiling. Raw data was drift corrected, the values of the metabolites were normalized to total sum of metabolites biometrically per sample, and $log_2$-transformed to stabilize variance and normalize the distributions. Errors represent ± SEM, $n$ = 5 - 6. *$p \leq 0.05$, **$p \leq 0.01$, ***$p \leq 0.001$, ****$p \leq 0.0001$ by One-way ANOVA. To control for multiple comparisons, p-values from post-hoc pairwise comparisons were adjusted using the Benjamini-Hochberg False Discovery Rate (FDR) procedure. The FDR threshold of $Q < 0.05$ was used to determine the statistical significance. Furthermore, outliers were detected and removed from the analysis using ROUT ($Q$ = 1 %) method.
